# Urban enrichment of *molestus* ancestry and opposing temperature clines in a metropolitan *Culex pipiens* hybrid zone

**DOI:** 10.64898/2026.10.30.763117

**Authors:** Benjamin P. Gregory, Arielle Arsenault-Benoit, Katherine L. Bell, Megan L. Fritz

## Abstract

Hybridization between ecologically distinct lineages can fuel rapid adaptation when introgressed alleles are locally favored. The North American *Culex pipiens* assemblage, a principal West Nile virus vector east of the Mississippi, comprises three lineages—temperate *Cx. pipiens* f. *pipiens*, urban-specialist *Cx. pipiens* f. *molestus*, and subtropical *Cx. quinquefasciatus*—that meet in a mid-Atlantic hybrid zone where temperature and urbanization gradients intersect. Using whole-genome Pool-seq of 24 field populations and three lineage-reference pools across the Baltimore–Washington corridor, we find a weakly differentiated, *Cx. pipiens* f. *pipiens*-like background, richer in *molestus*- and *quinquefasciatus*-like ancestry toward the warmer end of the corridor. Urbanization impacts allele frequencies across the genome: *molestus*-like ancestry rises with impervious surface, independently of temperature and position, while *Cx. quinquefasciatus*-like ancestry does not. An 11.6 Mb chromosome 3 inversion is *molestus*-like in every population, most of all in urban ones, and tracks genome-wide *molestus*-like ancestry. Temperature acts locus by locus. On chromosome 1, a cluster of six CYP6 paralogs, a family implicated in insecticide resistance and thermal-stress response, shows rising *Cx. quinquefasciatus*-like ancestry and read depth with temperature. A chromosome 2 interval centered on a *Yan* ortholog, a repressor of cell differentiation, shows non-*quinquefasciatus* ancestry increasing with temperature, consistent with selection against *Cx. quinquefasciatus*-like introgression. Urban populations therefore carry more ancestry from the human-commensal lineage, while temperature moves *Cx. quinquefasciatus* ancestry both ways at individual loci. Both *molestus* and *Cx. quinquefasciatus* donors carry human-contact traits, so these patterns bear on how warming and urbanization will reshape arboviral risk.

## Introduction

The footprint of mosquito-borne disease is shifting. Vector-borne zoonoses account for about a fifth of emerging human infections (Swei et al. 2020), and climate change and land-use change are complementary drivers of that trend (Patz et al. 2008). Because mosquitoes and their pathogens are ectotherms, temperature sets the outer bounds of where transmission is possible (Mordecai et al. 2019; Rocklöv and Dubrow 2020). Cities, where urban land is projected to keep expanding (Chen et al. 2020; Gao and O’Neill 2020), supply standing water for larvae (Gardner et al. 2012; LaDeau et al. 2013; Rochlin et al. 2019) and concentrate human hosts. In temperate regions, urban heat can increase larval development rates (LaDeau et al. 2015; Townroe and Callaghan 2014). Vectors adapt to these conditions. Population genomic studies have found signatures of local adaptation to climate in *Aedes aegypti* and *Anopheles gambiae* (Bennett et al. 2021; Cheng et al. 2012; Soudi et al. 2023). In *Ae. aegypti*, dry-season intensity and human population density predict most of the variation in preference for human hosts, a preference linked to a single genomic ancestry component whose spread into African cities dates to recent urbanization (Rose et al. 2020, 2023). Warming and urbanization may therefore do more than move the boundaries of transmission. They may favor the taxa, phenotypes, and alleles best suited to novel thermal and land-use conditions. How vector populations respond to these two forces, separately and together, is both an ecological question about where transmission happens and an evolutionary question about how the vectors themselves are changing.

Adaptive introgression, the selective retention of introgressed alleles that raise fitness in a recipient lineage (Burgarella et al. 2019), offers a faster route to environmental tracking than *de novo* mutation and is increasingly recognized as a major source of adaptive variation (Edelman and Mallet 2021; Pfennig 2021). In mosquitoes, adaptive introgression has been shown to take several forms. Selection on a pyrethroid-resistance mutation carried a multi-megabase genomic island along with it by hitchhiking from *An. gambiae* to *An. coluzzii* (Clarkson et al. 2014; Norris et al. 2015). Introgression can also act in tandem with gene duplication: the *Ace1* resistance haplotype, with a coding substitution and structural duplications, spread from *An. gambiae* into *An. coluzzii* (Grau-Bové et al. 2021). Inversions offer a further route: they can capture locally adapted multi-gene haplotypes and protect them from recombination (Kirkpatrick and Barrett 2015). In *An. darlingi*, a region that tops several selection scans and contains six cytochrome P450 genes lies within a segregating inversion (Tennessen et al. 2026).

The North American *Culex pipiens* assemblage is a natural system in which to ask whether warming and urbanization shape introgression. The assemblage maintains West Nile virus in birds and bridges transmission to humans across the eastern United States, and its members are tightly coupled to human habitats (Andreadis 2012; Ciota 2017; Kilpatrick et al. 2005, 2006; Rochlin et al. 2019). Its three lineages have niches that map onto the gradients of a modern city. Temperate, ornithophilic *Cx. pipiens* f. *pipiens* Linnaeus, 1758 overwinters through photoperiodically cued reproductive diapause (Denlinger and Armbruster 2014; Sim and Denlinger 2008). The bioform *molestus* (*Cx. pipiens* f. *molestus* Forskål, 1775) exploits below-ground and human-modified habitats, is facultatively autogenous, feeds preferentially on mammals including humans, and persists year-round in cities without obligate diapause (Bell et al. 2024; Fritz et al. 2015; Haba and McBride 2022). *Cx. quinquefasciatus* Say, 1823, is a subtropical sister species that also lacks diapause, feeds across host classes, and bridges avian and mammalian hosts (Andreadis 2012; Farajollahi et al. 2011). The three lineages hybridize where they meet, including across a *Cx. pipiens*–*Cx. quinquefasciatus* hybrid zone that extends into the mid-Atlantic (Cornel et al. 2003; Fonseca et al. 2004; Kothera et al. 2009; Saarman et al. 2025), and hybrids can differ from the parental forms in West Nile virus competence (Ciota et al. 2013), host preference (Fritz et al. 2015), and diapause capacity (Meuti et al. 2015). Because *molestus* and *Cx. quinquefasciatus* carry contrasting suites of disease-relevant traits, ancestry from one versus the other is not interchangeable: it determines which traits enter admixed populations.

The Baltimore–Washington corridor is an ideal setting in which to ask how environmental gradients shape introgression in this assemblage. It lies where all three lineages meet, at the northern edge of the *Cx. pipiens*–*Cx. quinquefasciatus* hybrid zone, classically placed between 36° and 39°N (Barr 1957; Cornel et al. 2003; Kothera et al. 2009), where hybrids still occur (Saarman et al. 2025; Yurchenko et al. 2020). It also includes two heavily urbanized cores where *molestus* has been recorded: historically in Baltimore (Twohy and Rozeboom 1957) and as a resident belowground population in Washington, DC (Arsenault-Benoit et al. 2021). Within a single ∼60 km region, the corridor spans the gradients that distinguish each lineage’s niche, summer thermal regime, indexed as maximum summer temperature (bio05) for *Cx. quinquefasciatus*, and urbanization, indexed as percent impervious surface for form *molestus*. Because it spans two metropolitan areas, the two vary separately: Baltimore and Washington differ in maximum summer temperature, while impervious surface varies independently within each.

Here, we ask how these environmental gradients structure introgression in this hybrid zone, treating *Cx. pipiens* f. *pipiens* as the recipient lineage and *Cx. pipiens* f. *molestus* and *Cx. quinquefasciatus* as the donors. We ask the question at two scales. First, across the genome, does each donor’s ancestry vary along the temperature and urbanization gradients, and does each gradient draw on a different donor? Second, at individual loci, does donor ancestry depart from that genome-wide level along a gradient? A genome-wide trend can arise from contact alone, because a population nearer a donor’s source carries more of that donor’s ancestry everywhere in its genome. Selection on introgressed material should appear at particular loci, as more or less donor ancestry than the same population’s genome carries, a population-specific excess or deficit we call Δ. Because the donors occupy contrasting niches, we predicted that Δ for *Cx. quinquefasciatus* ancestry would rise with summer temperature at some loci, and Δ for *molestus* ancestry would rise with impervious surface at others. We tested these predictions with whole-genome Pool-seq of 24 field populations spanning both gradients, together with a reference pool for each lineage (Fig. 1; Table S1).

**Figure 1.**
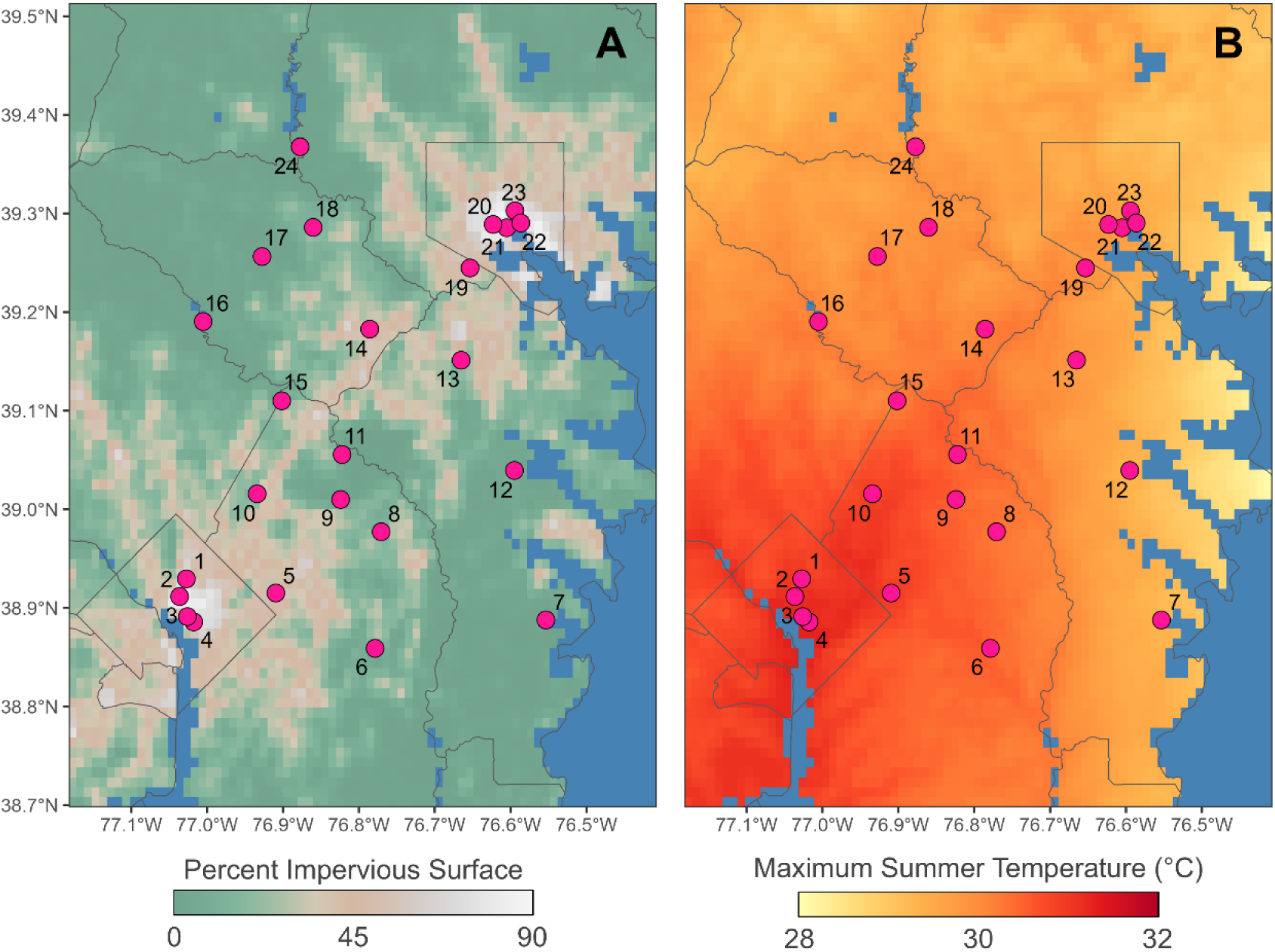
Field sampling sites across the Baltimore–Washington metropolitan corridor. Pink points mark the 24 *Culex pipiens* collection sites (numbered 1–24, with coordinates and pool details in Table S1) sampled for whole-genome Pool-seq. Both panels show the same sites overlaid on the two principal environmental gradients identified in the genotype-environment association analyses. (A) Percent impervious surface, the urbanization gradient. (B) Maximum summer temperature (°C). Rasters are displayed at 1km resolution from NLCD (A) and CHELSA Bioclim (B). Gray lines denote state and county boundaries.

## Materials and Methods

### Mosquito sampling, pool construction, and sequencing

We sampled adult female mosquitoes from 24 sites across the Baltimore, MD, and Washington, DC metropolitan areas, selected to span the regional urbanization and summer temperature gradients (Fig. 1; Table S1), with CDC gravid traps deployed overnight monthly in June, July, and August of 2023 and 2024. Specimens were confirmed as *Cx. pipiens* assemblage by an abdomen-maceration protocol (Gloor and Engels 1992) followed by a ribosomal DNA (ITS) PCR assay (Crabtree et al. 1995), preserving thoraces for sequencing. For each site we pooled thoracic tissue from 40 assemblage females, balanced across the six collection dates. Three reference pools were built identically from *Cx. quinquefasciatus* (Vero Beach, Florida; QUI), *Cx. pipiens* f. *pipiens* (Orono, Maine; PIP), and *Cx. p.* f. *molestus* (Calumet, Illinois; MOL). Libraries were sequenced on an Illumina NovaSeq S4 (150 bp paired- end) to a target of ∼20× mean coverage per pool. Trapping, extraction, and library details, like the software versions, parameters, and predictor sources for every analysis below, are given in Supplementary Methods under matching headings.

### Read processing, alignment, and variant calling

Reads were adapter- and quality-trimmed, aligned with BWA-MEM2 (Vasimuddin et al. 2019) to the *Cx. pipiens* reference assembly idCulPipi1 (Hesson et al. 2025), deduplicated with Picard (Broad Institute 2019), and filtered to properly paired reads with mapping quality ≥ 10. Variants were called across all 27 pools with PoolSNP (Kapun et al. 2020) from SAMtools pileups (Li et al. 2009), using only positions with coverage depths of at least 10× in every pool and below each pool’s per-chromosome 99th depth percentile.

### Population structure, differentiation, and isolation by distance

To detect population divergence in genome-wide structure, we conducted a principal component analysis (PCA) on the allele-frequency matrix (prcomp in R; Patterson et al. 2006). Pairwise F_ST_ among the 24 field pools and between each field pool and each reference was estimated in poolfstat with its Pool-seq analysis-of-variance estimator (Hivert et al. 2018), which extends that of Weir and Cockerham (1984), treating each pool as 80 chromosomes. Isolation by distance was tested by regressing linearized F_ST_/(1 − F_ST_) (Rousset 1997) on log_10_ geodesic distance across all 276 site pairs, with a Mantel test (9,999 permutations) in vegan (Oksanen et al. 2026). Eight site-level environmental predictors spanning the thermal and urbanization gradients were extracted within a 1 km diameter buffer around each site. Individual-level ddRADseq genotypes (553 mosquitoes, 7,258 SNPs) were analyzed by PCA to check that the field pools are composed of admixed individuals (Supplementary Methods).

### Genome-environment association with BayPass

To test for genome-environment associations, pool-seq read counts for the 24 field pools were analyzed in BayPass v3.1 (Gautier 2015). The core model estimated the population covariance matrix Ω and calibrated XtX_st_ statistics (Olazcuaga et al. 2020), and the covariate model tested association with eight centered, scaled predictors (elevation, impervious surface, tree canopy, water-table depth, population density, maximum summer temperature, temperature annual range, precipitation of the warmest quarter) using Ω. XtX_st_ thresholds were calibrated on 100,000 simulated neutral SNPs (Gautier 2015). Bayes factors (BF) of 10 and 30 dB marked minor and major outliers. Chain settings and thresholds are given in Supplementary Methods. We first identified SNPs that were at least minor outliers for both XtX_st_ and BF, then required each to belong to a chain of at least eleven SNPs that were at least minor outliers for either statistic, with consecutive positions no more than 10 kb apart. This spatial-support requirement excluded 14% of SNPs that met both statistical thresholds. In total, we retained 1,299 candidates for environmentally driven selection (hereafter, candidate SNPs), concentrated in three regions that we call CpL1a, CpL2a, and CpL3a.

### Genome-wide donor ancestry and environmental gradients

We measured donor ancestry with donor-private panels, defined as alleles at a frequency of at least 0.95 in one reference pool and no more than 0.05 in both others. The mean frequency of a panel’s alleles in a field pool, taken across the genome outside the three candidate regions, is that pool’s genome-wide ancestry from the donor (α). We regressed α for each lineage on maximum summer temperature and percent impervious surface, singly and together, across the 24 field pools. We repeated the regressions with a metropolitan-area term (assigned by nearest city center), with latitude and longitude, and with two winter-temperature layers from the same CHELSA climatologies (minimum temperature of the coldest month and mean temperature of the coldest quarter). We also tested impervious surface within each metropolitan area.

### Annotation liftover and gene-content surveys

To link candidate SNPs to genes, we projected the *Cx. quinquefasciatus* JHB2020 annotation onto the *Cx. pipiens* idCulPipi1 reference with Liftoff v1.6.3 (Shumate and Salzberg 2021) and used the lifted GFF3 for all annotation-based analyses. Genes overlapping each candidate interval were classified into candidate families, with orthology confirmed by BLASTP against SwissProt. Candidate SNPs were attributed to genes by counts and densities over the gene body (±2 kb flank) and CDS. Gene-body density was the primary ranking metric because adaptive variation often targets regulatory as well as coding sequence (Andolfatto 2005; Supplementary Methods).

### Targeted analysis of insecticide-resistance target sites

Because the genome scan retains only SNPs associated with a measured predictor, it cannot detect selection from an unmeasured pressure such as insecticide use. We therefore catalogued coding variation across 63 canonical insecticide-resistance target and detoxification genes independently of the scan, reading allele counts directly from the variant call set. The voltage-gated sodium channel (*Vgsc*) candidate was reconciled to standard housefly numbering by alignment to *Musca domestica* Vssc1 (UniProt Q94615). Gene lists, estimators, and the nulls used to test the *kdr* correlations are given in Supplementary Methods.

### Callable-position read-depth analysis

To test whether per-gene read depth covaries with summer temperature beyond genome-wide background, we fit a negative-binomial generalized linear mixed model to per-gene depth at PoolSNP-callable positions across the 24 field pools, summing callable depth within each gene’s CDS for 13,741 genes, with log per-pool genome-wide callable depth as an offset and a per-gene random intercept. Any mapping bias that varies with temperature across the corridor would shift depth at all genes, so we asked whether depth changed with temperature more at focal genes than across the rest of the genome. For each locus, we tested this with a focal-gene indicator and its interaction with temperature, comparing focal genes against the ∼13,740 non-focal genes. Reference pools were excluded.

### Locus-level diagnostics against a genome-wide null

To test whether each candidate region behaved as a coherent unit rather than a chance cluster of outlier SNPs, we compared its block structure, linkage disequilibrium, and donor ancestry against a common empirical null of 10,000 random autosomal regions of matched width, excluding the three candidate intervals, with one-sided empirical-percentile P-values. Because the candidate regions were selected for extreme differentiation, we repeated the block-structure tests against a stricter null additionally matched on differentiation. Block structure was summarized by the variance explained by PC1 of the per-population allele-frequency matrix and its correlation with each population’s mean F_ST_ to the donor reference. Linkage was the pairwise correlation (r²) between allele frequencies across the 24 field populations, computed for five SNP sets per locus and binned by distance. Donor references were QUI at CpL1a and MOL at CpL3a. The CpL2a candidate SNPs do not separate PIP from MOL, so we used QUI there and read the result on the QUI-versus-non-QUI axis.

### Per-population donor-ancestry test

Pooled data give allele frequencies, not genotypes, so donor ancestry in a pool could reflect introgression within individuals or unmixed individuals of different lineages pooled together. Only introgression should produce an excess of donor ancestry at a particular locus relative to the rest of the same pool’s genome. Using the donor-private panels, we computed α for each field pool inside each focal interval (α_locus_) and across the genome outside all three candidate regions (α_genome_). The difference Δ = α_locus_ − α_genome_ estimates locus-specific donor ancestry net of the genome-wide background. Because allele frequencies in an admixed population are a linear combination of its sources’ frequencies at every SNP (Patterson et al. 2012; Peter 2022), Δ has expectation zero at every locus under pure mixture, whereas environmentally structured introgression within individuals makes Δ exceed zero at the focal locus and track the environmental gradient. This expectation holds only if the panel marks donor ancestry equally well inside the locus and across the genome. Because inversions and other regions of suppressed recombination preserve fixed differences that recombination erodes elsewhere, we also regressed each locus’s donor ancestry on genome-wide donor ancestry and the predictor together. At CpL2a we used the QUI-deficit estimator 1 − α_QUI_, the combined non-QUI ancestry. We tested median Δ > 0 (one-sided Wilcoxon signed-rank) and the regression of Δ on the predictor against the width-matched null, and compared two-predictor models of Δ on maximum summer temperature and percent impervious surface with primary-predictor-only models by F-test. Panel thresholds and estimator algebra are given in Supplementary Methods.

### Windowed PCA and inversion breakpoints

To test whether the candidate regions could be detected without the genome scan or any environmental variable, we ran PCA on the 24 field pools in non-overlapping 500 kb and 1 Mb windows, ranked windows by the percentage of among-pool variance on the first principal component, and compared the windows’ local structure by multidimensional scaling (Li and Ralph 2019; Cebrián-Camisón et al. 2026). To locate inversion breakpoints, we called inversions from discordant read pairs in the duplicate-removed reference-pool alignments with DELLY v1.7.3 (paired-end support ≥ 5, MAPQ ≥ 30, inversions 50 kb–20 Mb), retaining breakpoint-spanning reads that a proper-pair filter would discard. Because Pool-seq does not resolve haplotypes, these calls were used only to locate breakpoints (Kapun et al. 2014).

### Use of artificial intelligence tools

We used Claude (Anthropic), a large language model, through the Claude Code interface to help write, revise, and document the R analysis code, including the figure scripts; to prepare the public code repository; and to edit the manuscript text. The authors decided which analyses to include, reviewed all AI-assisted code, results, and text, accepted or rejected every suggested change, and take full responsibility for the content of the paper.

## Results

### Mosquito collection and environmental variation across the corridor

The 24 sites spanned the two gradients that distinguish the niches of the three lineages, summer thermal regime and urbanization (Fig. 1; Table S1). Maximum summer temperature ranged from 29.2 °C to 31.2 °C, and percent impervious surface ranged from 0.5% to 85.3%. The two were only weakly correlated (r = 0.26; Fig. S1). Five of the eight predictors in the genome scan fell into two gradients, temperature (maximum summer temperature, temperature annual range) and urbanization (impervious surface, tree canopy, population density), and three were context variables (elevation, water-table depth, precipitation of the warmest quarter). Predictors within a gradient were far more strongly correlated with one another than with those of the other gradient, so a SNP’s association with one predictor could not be separated from its association with the others in that gradient. We therefore interpret each association as a response to its gradient and use maximum summer temperature and percent impervious surface as representatives (Supplementary Methods and Supplementary Results).

### Corridor populations are weakly differentiated and predominantly *Cx. p.* f. *pipiens*-like

Before asking whether any locus responds to the environment, we characterized the genomic background against which such a signal would have to stand out. Variant calling yielded 11,629,890 biallelic SNPs with complete allele-frequency data across pools, from 155.1 M positions that passed the depth filters. The first two principal components of the allele-frequency matrix separated the 24 field pools from the three reference pools (30.2% and 12.2% of variance; Fig. 2; additional axes in Fig. S2): the references sat at the extremes and the field pools formed a single tight cluster. Pools of unmixed individuals of different lineages would spread the field pools along the axes joining the references and make differentiation among them substantial, but differentiation was very low (mean pairwise F_ST_ 0.004; Fig. S3), and ddRADseq genotyping of 553 individuals from the region, including the source populations of all three references, places every field individual in a single continuous cloud spanning PIP and MOL, with none intermediate between it and the sharply separated QUI reference (Fig. S4; Supplementary Methods). Corridor populations are therefore composed of admixed individuals rather than mixtures of unmixed lineages or early-generation *Cx. quinquefasciatus* hybrids.

**Figure 2.**
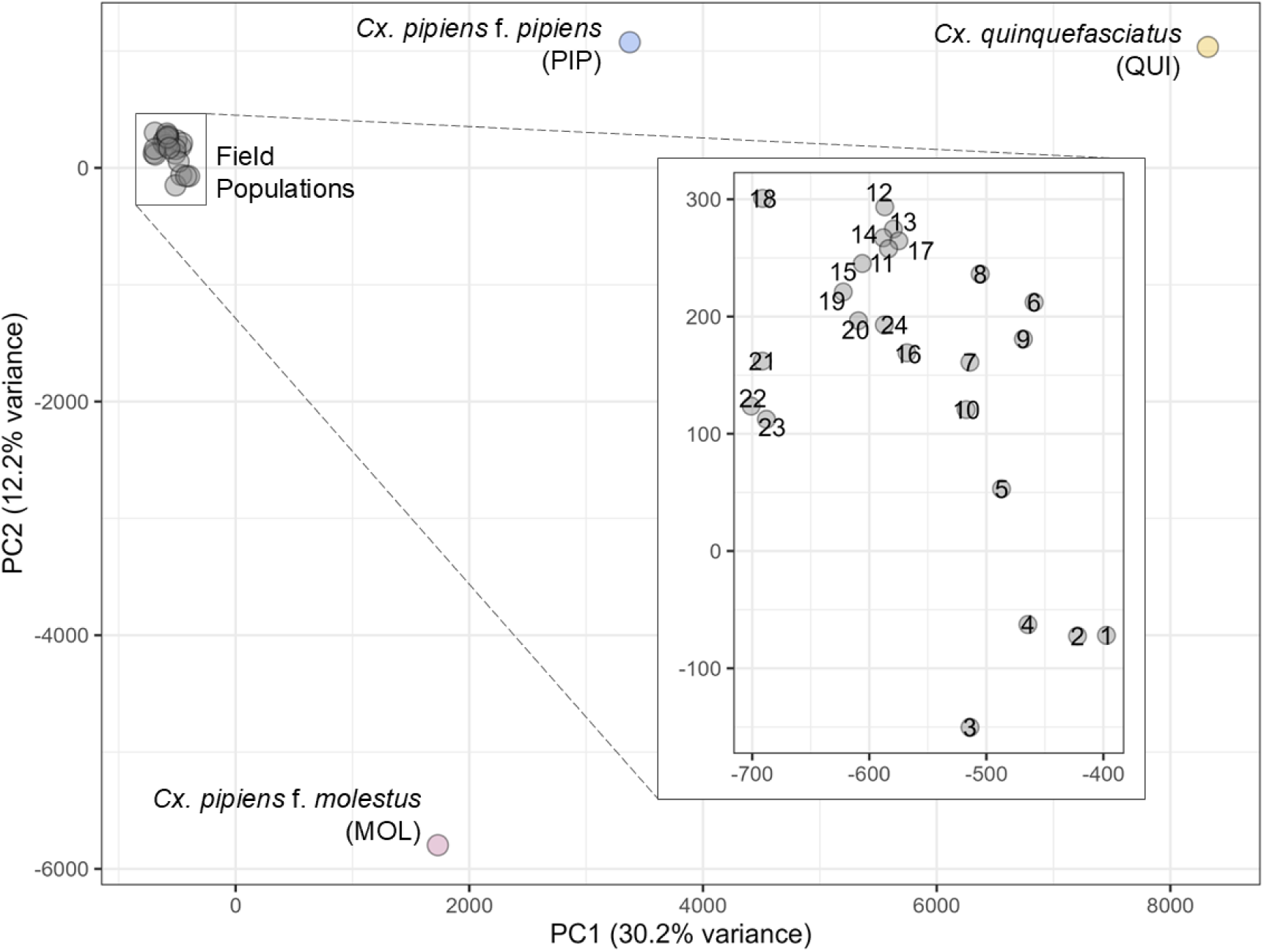
Genome-wide principal component analysis of field pools and reference taxa. PCA of genome-wide allele frequencies across all 27 Pool-seq libraries: the 24 field pools (gray, numbered 1–24) and the three reference pools—*Cx. pipiens* f. *pipiens* (PIP, blue), *Cx. quinquefasciatus* (QUI, yellow), and *Cx. pipiens* f. *molestus* (MOL, pink). PC1 (30.2% of variance) and PC2 (12.2%) are shown. The three references occupy the extremes of the ordination, and the field pools cluster tightly together near the origin, consistent with admixed corridor populations distinct from any single reference taxon. The inset rescales the field-pool cluster to resolve among-pool structure. Note the inset axes span only ≈300 PC1 units (≈ −700 to −400) and ≈450 PC2 units (≈ −150 to 300), a small fraction of the full ordination, so apparent separation among field pools is small relative to the field-versus-reference divergence.

The corridor background was predominantly *Cx. pipiens* f. *pipiens*-like. Across the field pools, the mean frequency of PIP-private alleles (α) was 0.366, against 0.158 for MOL-private and 0.108 for QUI-private alleles, an ordering that held when panel thresholds were varied (Supplementary Results). Genome-wide F_ST_ gave the same ordering (Table S2) but is deflated by a population’s own diversity, so we did not use it to measure ancestry (Fig. S5; Supplementary Results). Panel sizes also differed by more than an order of magnitude between donors, so we did not compare panel sizes or absolute ancestry magnitudes across donors (Supplementary Results).

Differentiation among field pools also tracks geographic distance: linearized F_ST_/(1 − F_ST_) rose with log_10_ geodesic distance across all 276 pairs (Pearson r = 0.538, one-sided P < 0.001; Mantel test P < 0.001; Fig. S6), with distance explaining roughly a third of the variation (adjusted R² = 0.29), consistent with active but distance-limited gene flow across the corridor.

### Genome-wide donor ancestry is structured by both temperature and urbanization

Using linear regressions across the 24 field pools, we asked whether each lineage’s genome-wide ancestry (α) varied along either gradient. QUI-like ancestry rose with maximum summer temperature (+0.004 per °C, adjusted R² = 0.45) but not with impervious surface, and temperature kept its effect with both gradients in the model. MOL-like ancestry rose with both temperature (+0.019 per °C, adjusted R² = 0.44) and impervious surface (adjusted R² = 0.41), each keeping its effect with the other in the model (joint adjusted R² = 0.68). PIP-like ancestry fell with temperature (−0.013 per °C, adjusted R² = 0.40) and was not correlated with impervious surface. Every rise or fall reported here has P < 0.001 (Supplementary Results). With metropolitan area (Baltimore or Washington, DC) added as a categorical covariate, the QUI-like and MOL-like temperature trends persisted but the PIP-like decline did not (Supplementary Results). Warmer sites therefore carry more ancestry from both donors and, across the corridor as a whole, less from the temperate background lineage, while more urban sites carry more *molestus*-like ancestry specifically.

These trends differ in what they can show. Summer temperature follows position along the corridor, warming toward Washington and the *Cx. quinquefasciatus* range (Supplementary Results), so the rise in QUI-like ancestry is consistent with proximity to that range. The rise in MOL-like ancestry has a different candidate source, the resident *molestus* populations of the two cities. Impervious surface has no spatial trend along the corridor (R² = 0.000 on latitude and longitude), and its association with MOL-like ancestry persisted with position, temperature, or metropolitan area in the model, held within Washington (r = 0.90, P < 0.001), and ran in the same direction within Baltimore (r = 0.55, P = 0.08; Supplementary Results).

### A genome scan identifies three candidate regions with distinct architectures

A locus under selection should track its gradient beyond these genome-wide trends. The BayPass scan for SNPs that were outliers for both differentiation (XtX_st_) and environmental association yielded 1,299 candidate SNPs concentrated in three regions (Fig. 3A), designated CpL1a, CpL2a, and CpL3a. Maximum summer temperature was by far the most frequently assigned predictor across candidate SNPs (64.4%; Supplementary Results). CpL1a, on chromosome 1 (∼108.5–111.0 Mb), is the densest: a compact ∼2.5 Mb peak of 654 candidate SNPs, half the genome-wide total, 84.4% of them most strongly associated with maximum summer temperature. CpL2a, on chromosome 2 (∼13.5–14.0 Mb), is a compact 500 kb cluster of 110 candidate SNPs with nearly the same predictor composition (81.8% maximum summer temperature). CpL3a, on chromosome 3 (149.4–161.0 Mb), is structurally distinct: 69 SNPs with broadly elevated XtX_st_ across the whole ∼11.6 Mb interval and no localized peak, driven mainly by the urbanization predictors. Tree canopy and percent impervious surface, which are near- collinear across our sites (r = −0.90), together accounted for just over half its candidate SNPs (Supplementary Results). The three loci differ systematically in donor lineage, environmental gradient, direction, and inferred mechanism (Table 1).

**Figure 3.**
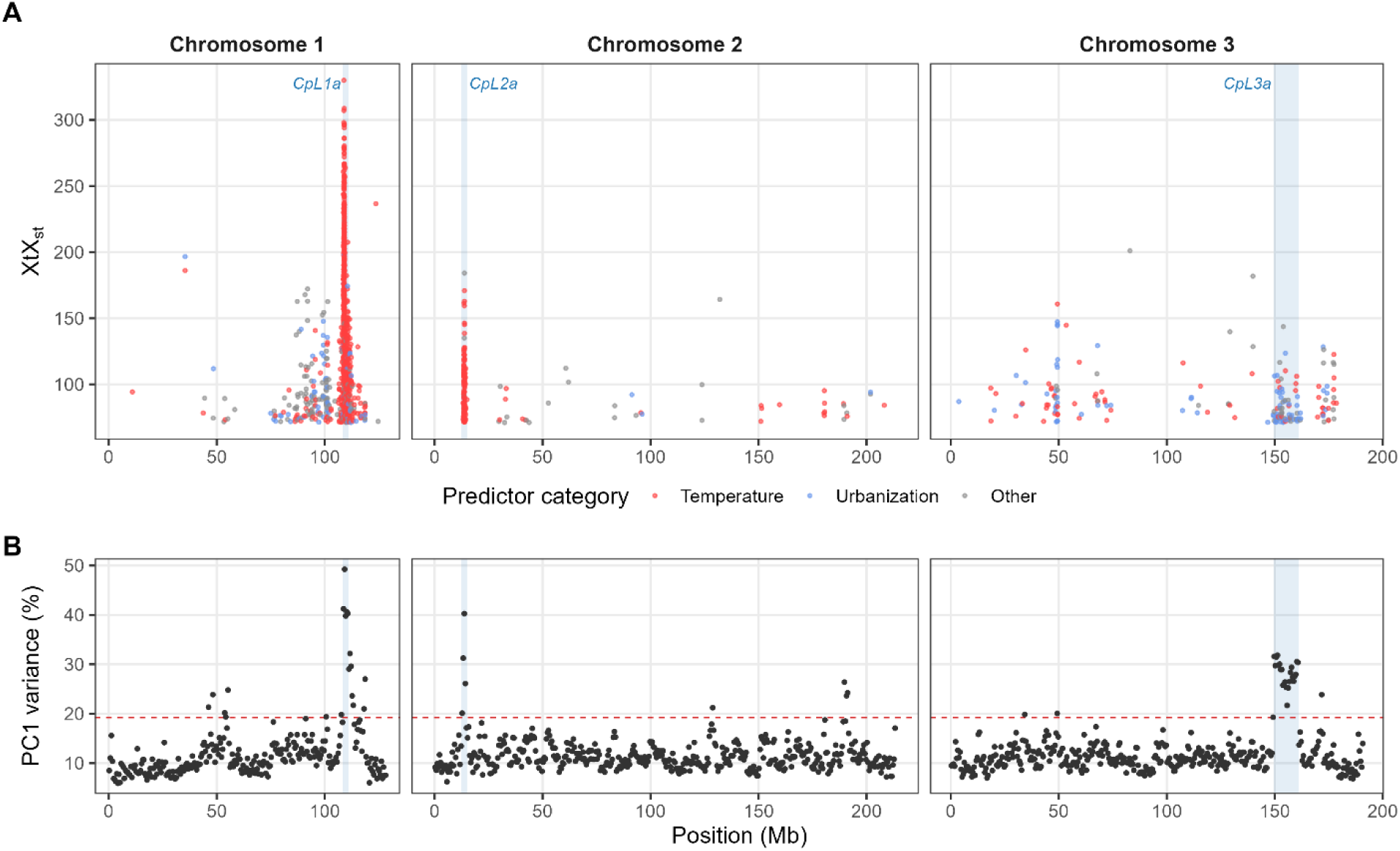
The genome scan and an environment-free windowed PCA identify the same three candidate loci. (A) Per-SNP XtX_st_ from the BayPass core model, plotted by genomic position for each chromosome. Only SNPs exceeding the POD-calibrated XtX_st_ significance threshold (XtX_st_ = 71.1, the lower axis limit) are shown, so every plotted point is an outlier. Higher XtX_st_ indicates stronger allele-frequency differentiation among the 24 field pools relative to the covariance-corrected null. Points are colored by predictor category: temperature (red), urbanization (blue), or other (grey). Shaded bands mark the three candidate loci: CpL1a (chr1, ∼108.5–111.0 Mb, CYP6 cluster), CpL2a (chr2, ∼13.5–14.0 Mb, Yan-centered interval, shaded at its 2.5 Mb block), and CpL3a (chr3, ∼149.4–161.0 Mb, inversion 3q). Because the y-axis is truncated at the significance threshold, the figure shows how outliers are distributed rather than how exceptional the three loci are relative to the genome as a whole. Scattered outliers elsewhere do not form comparable clusters, but they are not few: 466 of the 1,299 candidate SNPs (35.9%) fall outside all three candidate intervals—287 on chr1, 64 on chr2 and 115 on chr3—and the predictor composition of these unclustered outliers is given in Supplementary Results. (B) Percentage of among-pool variance on the first principal component in each non-overlapping 500 kb window, from PCA of the 24 field pools without environmental data. The dashed line marks the genome-wide 95th percentile, and shading marks the candidate intervals (CpL2a at its 2.5 Mb block). The 25 most structured windows in the genome lie within 2 Mb of a candidate locus.

**Table 1.** Summary of the three environmentally structured loci across the Baltimore– Washington *Culex pipiens* corridor. Each locus is a localized exception to a genome-wide background of low among-population differentiation, distinguished by donor lineage, environmental gradient, the direction of the ancestry-environment relationship, and the inferred mechanism. CpL1a and CpL2a track the same temperature gradient in opposite directions, whereas CpL3a follows urbanization independently of temperature, in step with genome-wide *molestus*-like ancestry.

| **Locus** | **Location** | **Donor ancestry** | **Environmental gradient** | **Inferred mechanism and direction** | **Focal feature** |
|---|---|---|---|---|---|
| CpL1a | chr1, ~108.5–111.0 Mb (~2.5 Mb) | QUI (*Cx. quinquefasciatus*) | Maximum summer temperature | Introgression with copy-number expansion. QUI-like ancestry, read depth, and sequence similarity rise with temperature. | CYP6 paralog cluster (6 paralogs, ~28.4 kb) |
| CpL2a | chr2, ~13.5–14.0 Mb core (~2.5 Mb block) | non-QUI (*Cx. pipiens* f. *pipiens* + *Cx. p.* f. *molestus*) | Maximum summer temperature | Selection against QUI-like alleles. Non-QUI ancestry rises with temperature, the opposite direction to CpL1a. | *Yan* ortholog (CQUJHB014704), regulatory signal |
| CpL3a | chr3, ~149.4–161.0 Mb (~11.6 Mb) | MOL (*Cx. p.* f. *molestus*) | Percent impervious surface (urbanization) | Carried with urban *molestus*-like ancestry. MOL-like ancestry rises with urbanization, in step with the genome-wide background. | Inversion 3q |

† Abbreviations: QUI, *Cx. quinquefasciatus*; PIP, *Cx. pipiens* f. *pipiens*; MOL, *Cx. pipiens* f. *molestus*; CpL1a, CpL2a, and CpL3a, the three candidate loci; chr, chromosome; Mb, megabase pairs; ~, approximately.

A scan that never sees an environmental variable finds the same three regions. Windowed PCA ranked the 1,066 non-overlapping 500 kb windows by the share of among-pool variance on the first axis, which measures how strongly allele frequencies within a window vary together across populations whatever the cause. Large mosquito inversions appear in such data as segregating haplotype blocks (Cebrián-Camisón et al. 2026; Tennessen et al. 2026). The 25 most structured windows all lay within 2 Mb of a candidate locus, with 27.4–49.2% of among-pool variance on the first axis against a genome-wide median of 10.8% (Fig. 3B), and the first window elsewhere ranked 26th, so the three regions are the most structured features of the genome, independently of the association scan.

A survey of 63 canonical insecticide-resistance genes (Table S3) found that the target-site allele *kdr* (*Vgsc* c.3105A>T, p.Leu1035Phe; L1014F in housefly numbering) was as strongly differentiated as the candidate SNPs (XtX_st_ = 128.2) but had no environmental support (best BF = −1.9 dB). Its frequency tracked neither maximum summer temperature (P = 0.32) nor percent impervious surface (P = 0.16; Fig. S7).

### *Cx. quinquefasciatus* ancestry and sequencing read depth at a CYP6 cluster rise with summer temperature

CpL1a spans ∼2.5 Mb on chromosome 1. A temperature cline there could reflect selection on locally segregating variation as readily as on introgressed variation, so we asked whether donor ancestry at the locus tracks temperature, and from which lineage, using donor-private panels defined across all SNPs in the interval, which scarcely overlap the candidate SNPs (Supplementary Methods). The interval held 123 QUI-private and 3,762 MOL-private SNPs but only one PIP-private SNP, so PIP-like ancestry could not be measured at this locus (Table S4). The two panels moved in opposite directions. QUI-like ancestry rose steeply with summer temperature, from a mean of 0.088 across the six coolest sites to 0.234 across the six warmest (adjusted R² = 0.64, P < 0.001), while MOL-like ancestry fell, from 0.117 to 0.089 (adjusted R² = 0.15, P = 0.033). Warmer sites therefore carry more *Cx. quinquefasciatus*-like and less *molestus*-like ancestry at this locus, although genome-wide they carry more of both, and the candidate SNPs show the same shift (Supplementary Results).

Lifting the *Cx. quinquefasciatus* gene annotation onto the *Cx. pipiens* reference allowed us to link candidate SNPs to genes. Six clustered CYP6 paralogs among CpL1a’s 79 protein-coding genes ranked first through sixth on candidate-SNP density, and the cluster was collinear between the two assemblies (Table S5; Figs. S8, S9; Supplementary Results). All six carried candidate SNPs within coding sequence, 13 of which changed the encoded amino acid, at least one in every paralog (Table S6). Those substitutions were near-fixed between QUI and MOL but not between QUI and PIP. At the coding sites under selection, the functional contrast was therefore QUI/PIP versus MOL.

Whether the candidate SNPs travel together as one introgressed tract or are independently selected sites that share an environmental association determines the unit of selection at CpL1a. Pairwise correlations (r²) in allele frequency across the 24 populations were strong among candidate SNPs across the whole 2.5 Mb cluster (median r² = 0.529, roughly 23 times baseline) but weak, and indistinguishable from baseline by 5–8 Mb, for pairs reaching outside the interval. CpL1a is therefore a tightly linked set of SNPs concentrated at the CYP6 cluster and embedded in freely recombining sequence, the pattern expected when selection maintains association among introgressed variants without dragging an extended haplotype (Fig. 4, CpL1a).

**Figure 4.**
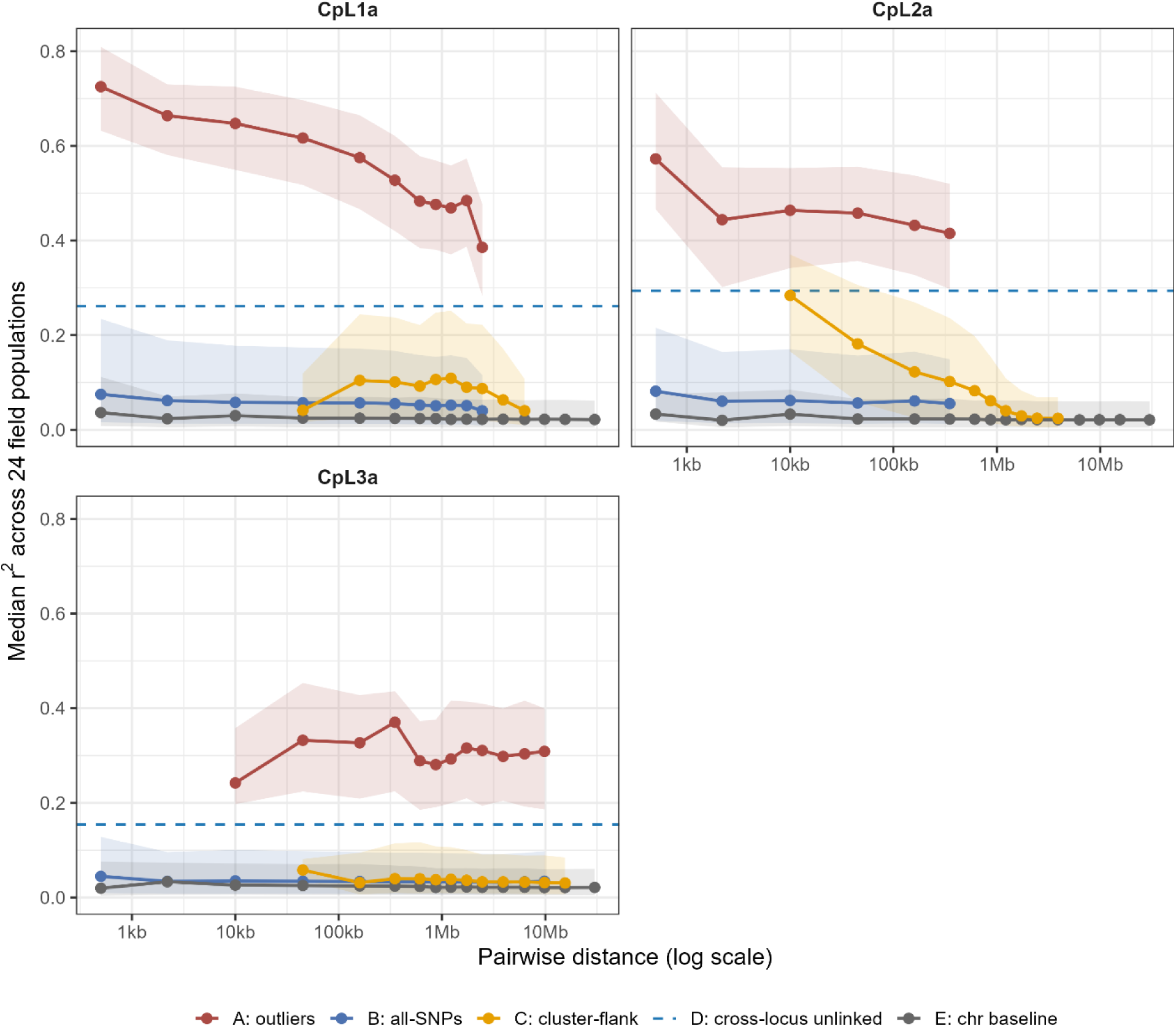
Linkage disequilibrium decay distinguishes three architectures at the candidate loci. Pairwise allele-frequency correlation (r²) across the 24 field pools as a function of genomic distance between SNP pairs (log scale), faceted by locus (CpL1a, CpL2a, CpL3a). Points are the median r² within each distance bin and shaded bands are the interquartile range (25th–75th percentile) of pairwise r² in that bin. Five SNP sets are shown: (A) within-cluster candidate SNPs (red), (B) all SNPs spanning the broader candidate interval (blue), (C) pairs formed between a candidate SNP and a SNP outside the cluster, binned by distance (gold), (D) pairs formed between this locus’s candidate SNPs and those of the other two loci, indexing the genome-wide donor-ancestry covariance expected between physically unlinked SNPs (dashed line); because set D is itself composed of environmentally associated SNPs it is inflated by shared ascertainment, so set E is the conservative baseline, and (E) a same-chromosome random baseline (gray). Because those pairs have no within-chromosome distance, D is shown as a single horizontal dashed line per panel rather than a decay curve. The diagnostic contrast is set A relative to D and E: correlation in excess of the cross-chromosome control reflects physical linkage beyond admixture covariance, and the distance behavior of A and C distinguishes the three architectures. CpL1a shows a tightly correlated focal cluster that decays only modestly across the cluster, with flank pairs weakly correlated at every distance (median r² 0.04–0.11) and indistinguishable from baseline by ∼5–8 Mb. CpL2a shows flank pairs beginning near median r² ≈ 0.28 beside the core and decaying smoothly to baseline by ∼2–3 Mb, delineating an extended donor haplotype block; the contrast between the two loci is therefore in amplitude beside the core rather than in the distance at which baseline is reached. CpL3a shows distance-independent correlation across the full ∼11.6 Mb interval that collapses abruptly to baseline at the candidate boundaries, the signature of suppressed recombination within an inversion.

These clines alone do not show that the signal was specific to CpL1a, because populations in warmer environments carrying more *Cx. quinquefasciatus* ancestry genome-wide would show the same clines. We therefore asked whether each population carried more QUI ancestry at CpL1a than it carried genome-wide (Δ on the QUI-private panel), and if any excess ancestry tracked summer temperature. Δ ranged from −0.038 to +0.207 and tracked summer temperature closely (adjusted R² = 0.62, P < 0.001, exceeding every width-matched and differentiation-matched comparison region; Supplementary Results). The four largest values fell at the four warmest urban Washington sites (Fig. 5, CpL1a). With both gradients in the model, temperature kept its effect (P < 0.001) while impervious surface contributed nothing (P = 0.16), so the locus-specific signal was structured by summer temperature, not urbanization.

**Figure 5.**
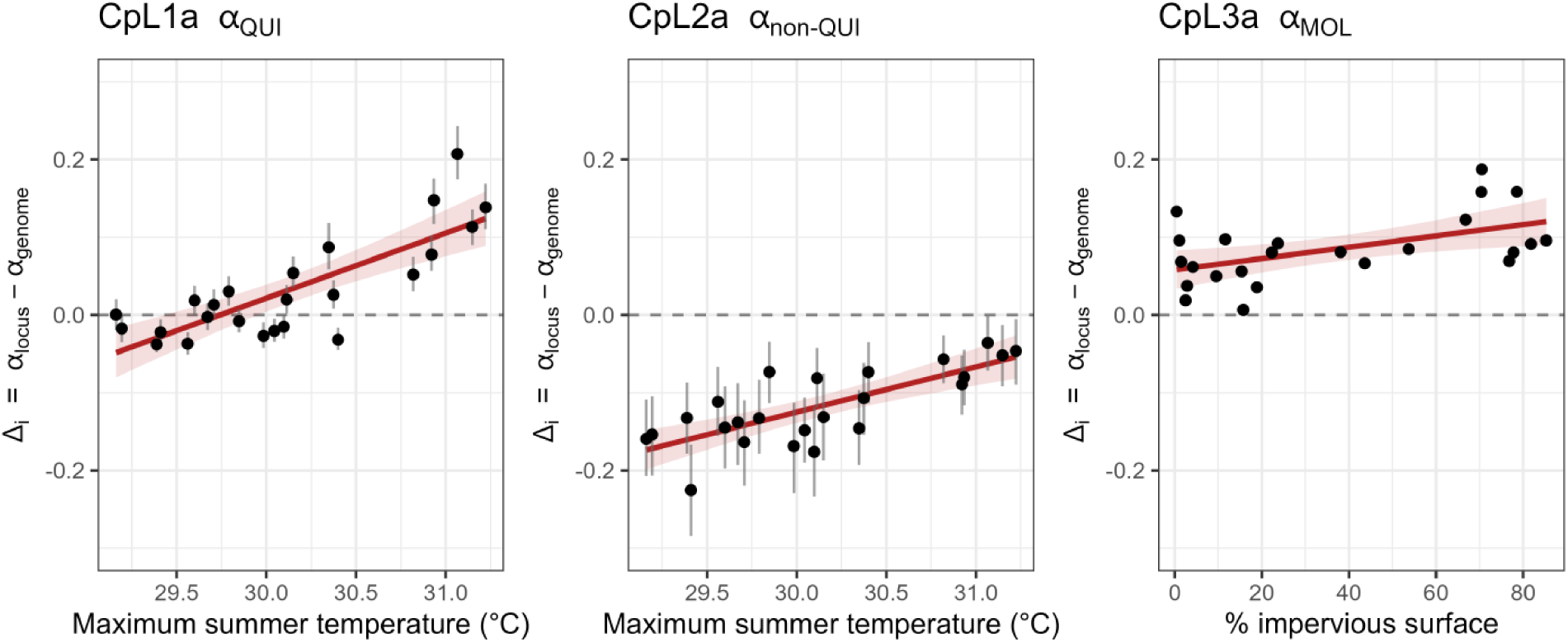
Locus-specific donor ancestry deviates from genome-wide background as a function of the relevant environmental gradient. Per-pool donor-ancestry deviation Δᵢ = α_locus,i_ − α_genome,i_ at each of the three candidate loci, where α is the mean pool allele frequency across donor-private panel SNPs and the subtraction removes each pool’s genome-wide donor ancestry (panel SNPs outside all three candidate regions). Each black point is one of the 24 field pools. (CpL1a) α_QUI_ versus maximum summer temperature. (CpL2a) combined non-QUI ancestry, estimated as the QUI deficit 1 − α_QUI_, versus maximum summer temperature. (CpL3a) α_MOL_ versus percent impervious surface. Gray vertical bars are per-pool 95% bootstrap confidence intervals on Δᵢ (B = 1,000 SNP resamples drawn independently within the focal and genome panels). The red line is an ordinary least-squares fit with its 95% confidence band, and the dashed line marks Δ = 0 (no locus-specific enrichment). The one-sided Wilcoxon signed-rank test of the marginal hypothesis (median Δ > 0) gave P = 0.051 at CpL1a, P = 1 at CpL2a and P = 6 × 10⁻⁸ at CpL3a, and the permutation P-value of the regression slope was 0.00013, 0.00013 and 0.0001, respectively. The permutation P-value is an empirical percentile against those width-matched random regions returning a valid statistic, which is fewer than the 10,000 drawn and differs among loci (7,679 to 9,677), so the finest attainable P-value differs accordingly. At CpL1a and CpL2a the signal is carried by the positive environmental slope rather than by a uniformly elevated median Δ; at CpL3a the median Δ is elevated across the corridor as well. At CpL3a, part of the slope of Δ on impervious surface is inherited from genome-wide *molestus*-like ancestry, which itself rises with impervious surface (Results).

Introgression can deliver extra gene copies as well as different alleles, and higher read depth at a gene in pooled sequencing may indicate higher copy number. As a ratio to the pool’s median gene depth, the mean depth of the six CYP6 paralogs rose with maximum summer temperature about 18 times more steeply than the genomic average (P < 0.001), and all six paralogs rose individually (Fig. 6; Supplementary Results). MOL carried the cluster at background depth, whereas PIP and QUI carried the higher-depth state. The field values therefore described a copy-number increase whose frequency rose with temperature. Cluster depth covaried with QUI-like ancestry at the locus (adjusted R² = 0.70) but not with MOL-like ancestry, which placed the extra copies on the introgressed sequence and mirrored the coding-SNP result (QUI/PIP versus MOL). Controls against a mapping artifact, and the shift of CYP6 consensus sequences toward QUI with rising temperature, are in Supplementary Results.

**Figure 6.**
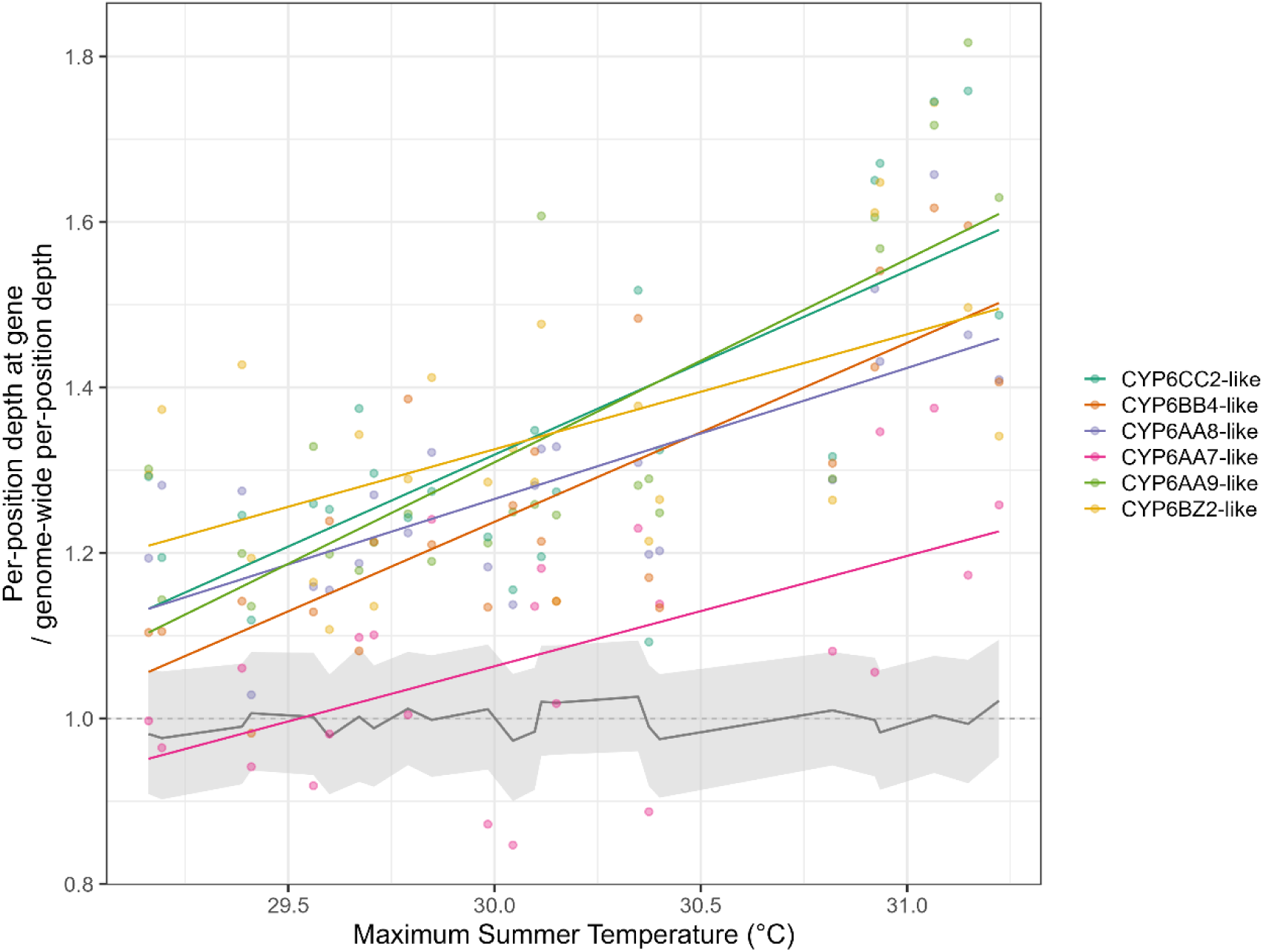
Relative read depth across the CpL1a CYP6 paralog cluster rises with summer temperature, consistent with a copy-number cline. For each field pool, per-position read depth at each of the six focal CYP6 paralogs is divided by that pool’s genome-wide per-position depth. A value of 1.0 (dashed line) therefore denotes genome-average depth, and values above 1.0 denote locally elevated depth. Points are per-pool, per-paralog ratios, and colored lines are per-paralog ordinary-least-squares fits shown for display. The grey line and ribbon give the median and interquartile range of the same ratio across all annotated background genes, which stay near 1.0 across the temperature range. Five of the six paralogs sit above the background across the temperature range and rise with maximum summer temperature; CYP6AA7-like sits at or near background at the cool end of the gradient and rises more shallowly. Statistical support is the focal-by-temperature interaction from a negative-binomial GLMM of per-gene total depth, fit with a genome-wide callable-depth offset and a per-gene random intercept on field pools only (β = 0.082, P < 0.001). The interaction tests whether depth at the cluster scales with temperature more steeply than the genome-wide background, isolating a focal copy-number response from any genome-wide depth-temperature trend.

### *Cx. quinquefasciatus* ancestry is excluded at a *Yan*-centered locus on the same gradient

The second region, CpL2a, is a compact 500 kb cluster on chromosome 2 whose candidate SNPs were associated with the same predictor as CpL1a. Applying the same private-panel test, we expected the same direction, but the association was reversed on both panels. QUI-like ancestry fell with summer temperature, against its genome-wide rise, from a mean of 0.325 across the six coolest sites to 0.146 across the six warmest (adjusted R² = 0.50, P < 0.001), while MOL-like ancestry rose, from 0.085 to 0.174 (adjusted R² = 0.66, P < 0.001). CpL1a and CpL2a are therefore mirror images on the same gradient. The QUI-private panel at CpL2a holds only six SNPs, three at consecutive positions and so effectively one marker, but relaxing the panel definition and resampling the markers leaves the slope essentially unchanged (Supplementary Results).

The candidate SNPs could not identify a donor lineage. None was fixed between PIP and MOL, and at the 27 that separated QUI from PIP (|Δp| ≥ 0.70), MOL nearly matched PIP (mean |Δp| = 0.11). These SNPs therefore distinguish QUI from non-QUI alleles, although PIP and MOL do differ elsewhere in the interval, with 39 of its 10,866 SNPs fixed between them (Supplementary Results). None of the 110 candidate SNPs fell in a private panel, so the panels measured the ancestry of the surrounding haplotype rather than the selected sites. On that measure, the background shifted toward a MOL-like state as sites warmed, but we do not identify the selected variant as *molestus*-derived (Supplementary Results).

Linkage also differed (Fig. 4, CpL2a; Supplementary Results). Candidate SNPs were correlated with one another (median r² = 0.444) and with the surrounding sequence, and that correlation decayed smoothly to baseline by 2–3 Mb. The locus therefore sits inside a donor haplotype block roughly 2.5 Mb wide. At CpL1a, by contrast, candidate SNPs were correlated with one another but only weakly with the surrounding sequence.

Because the candidate SNPs did not separate PIP from MOL, we measured the combined non-QUI fraction (one minus QUI-like ancestry). CpL2a carried less non-QUI ancestry than the genomic background at every one of the 24 sites, so the locus sits on an elevated baseline of QUI ancestry. Yet Δ varied among populations and tracked temperature (adjusted R² = 0.55, P < 0.001). Temperature retained its effect in a joint model, and impervious surface contributed nothing (P = 0.84). Warmer populations therefore carried more *Cx. pipiens s.l.*-like (PIP/MOL) ancestry despite the elevated QUI baseline (Fig. 5, CpL2a).

The signal concentrated most strongly on a single gene within the interval, an ortholog of *Yan* (CQUJHB014704). *Yan* is an ETS-family transcriptional repressor that keeps cells undifferentiated until receptor tyrosine kinase signaling switches it off (Rebay and Rubin 1995; Rogge et al. 1995; Webber et al. 2018). It carried 17 of the interval’s 110 candidate SNPs in its gene body, more than twice any other gene, as well as the highest single-SNP XtX_st_. Sixteen of the 17 SNPs fell in intronic or untranslated sequence, which suggested a regulatory signal (Table S7). Unlike CpL1a, read depth at *Yan* showed no relationship with temperature (P = 0.60), so this was allele-frequency variation without copy-number variation. The surrounding ∼2.5 Mb haplotype block also carries diapause-related genes, including a *takeout*-like ortholog.

### A *molestus* inversion carries urban *molestus*-like ancestry, independently of temperature

The third region is unlike the other two. Its uniformly elevated differentiation across ∼11.6 Mb of chromosome 3 suggested a segregating inversion, which would make the whole block, rather than any gene within it, the unit that changes in frequency. Three independent lines of evidence support that reading. First, the region behaved as a single haplotype block. Its primary axis of allele-frequency variation across the 24 populations corresponded almost exactly to how *molestus*-like each population was (PC1 = 28.1% of variance; correlation with mean F_ST_ to MOL, |r| = 0.995; both significant against the genome-wide null), so the broad differentiation reflected one block changing in frequency, not independent selection at the many genes it spans. In the windowed PCA, CpL3a was also the only extended region with a shared, distinct local structure (Fig. 3B, Fig. S10; Supplementary Results).

Second, linkage was flat across the entire interval. Candidate SNPs stayed correlated at median r² ≈ 0.30 at every distance across the block, with no decay, and then collapsed abruptly to baseline at the interval’s edges (Fig. 4, CpL3a), the signature of suppressed recombination inside an inversion and the opposite of the decay seen at CpL1a and CpL2a.

Third, the boundaries coincided with physical breakpoints recovered independently. Discordant read-pair mapping placed breakpoints at chromosome 3 ∼149.4 Mb and ∼161.0 Mb, with the strongest support in the MOL reference. These coordinates agree closely with inversion 3q identified by Cebrián-Camisón et al. (2026) through windowed PCA in a *Cx. pipiens* population from southwestern Spain (∼149–161 Mb on the same idCulPipi1 assembly), where all *molestus* specimens carried the inverted homokaryotype, so CpL3a is identifiable as that inversion (Supplementary Results).

We next asked whether the inversion tracked the environment, and which gradient. Applying the per-population ancestry test with *molestus*-private alleles, we found CpL3a consistently more MOL-like than the genomic background (median Δ = 0.081, robust to excluding corridor-common alleles; Supplementary Results). Unlike CpL1a, this elevation held across the corridor. Δ ranged from +0.007 at a suburban site to +0.188 in urban Washington and tracked percent impervious surface (adjusted R² = 0.245, one-sided P = 0.004). With both gradients in the model, impervious surface retained its effect while temperature contributed nothing (P = 0.20), and adding a metropolitan-area term did not change this (Supplementary Results). Part of that association was inherited from the genome-wide background that Δ subtracts. Δ assumes that a neutral locus gains donor ancestry one-for-one with the rest of the genome, and the inversion did not: its *molestus*-like ancestry rose 2.4 times as fast as the genome-wide level (adjusted R² = 0.52). With genome-wide *molestus*-like ancestry and impervious surface both in the model, the genome-wide level predicted the inversion’s *molestus*-like ancestry (P = 0.006) while impervious surface added no significant effect (P = 0.21). At CpL1a and CpL2a, temperature predicted locus ancestry beyond the background (P = 0.001 and 0.002). The inversion was therefore *molestus*-like throughout the corridor and most so where the genome as a whole was most *molestus*-like (Fig. 5, CpL3a).

Candidate SNPs were spread thinly across the inversion’s gene content, with no gene carrying more than two in coding sequence (Table S8), consistent with the block, not any single gene, being the unit that changed in frequency. The inversion contains a gustatory receptor and an odorant receptor, *Gr74* and *Or43* (Menna et al. 2026; Ryazansky et al. 2024). These are of note given host-feeding differences associated with *molestus*, although the block was not enriched for chemosensory receptors relative to the genomic background (Supplementary Results). We do not speculate which gene may be a target of selection, because the inversion captures all genes within it as one linked set.

## Discussion

Admixture supplies genomic variation to this ∼60 km metropolitan *Culex pipiens* assemblage hybrid zone, while urbanization and temperature sort that variation from different donors through distinct routes. *Molestus*-like ancestry rises with urbanization across a weakly differentiated, predominantly *Cx. pipiens* f. *pipiens*-like background, with a chromosome-scale inversion as its clearest marker, whereas temperature acts locus by locus, favoring *Cx. quinquefasciatus*-like ancestry at a compact CYP6 cluster and excluding it from an extended haplotype on chromosome 2 centered on *Yan*.

CpL1a provides the strongest case for introgression tied to the thermal gradient. QUI-like ancestry rises with summer temperature at CpL1a and, at the warm end of the corridor, exceeds each population’s genomic background (Fig. 5). The signal concentrates on six CYP6 paralogs with 13 amino acid substitutions and read depth that rises with temperature (Fig. 6).

The ancestry of the functional state is nevertheless more nuanced than a three-lineage label suggests. The nonsynonymous substitutions separate QUI and PIP from MOL rather than QUI from PIP, and the QUI and PIP references both carry the higher-depth state, whereas MOL sits at background depth. One explanation is that the amplified, protein-altering CYP6 haplotype entered *Cx. pipiens* f. *pipiens* through historical northward introgression from *Cx. quinquefasciatus*, and was already present across much of the temperate range before corridor populations encountered *molestus*. The cline we observe would then reflect a secondary process: at warmer corridor sites, the amplified haplotype is favored over the MOL low-copy state, and QUI/PIP-like ancestry at CpL1a displaces MOL-like ancestry.

Any phenotype that drives that cline remains unresolved, but two explanations fit it, alone or in combination. The first is thermal tolerance: southern *Cx. quinquefasciatus* larvae tolerate high temperatures better than northern *Cx. pipiens* larvae, and larvae from Maryland populations within and near the hybrid zone are intermediate (Gregory et al. 2026), so rising QUI-like ancestry at the warm end of the corridor is the direction a thermal advantage would produce. The second is insecticide resistance. Gene amplification is a recurrent mechanism of metabolic resistance (Bass and Field 2011), CYP6-family overexpression has repeatedly been tied to pyrethroid resistance in *Culex* (Liu et al. 2011; Zou et al. 2019, 2022), and amplification together with introgression at CpL1a recapitulates the *Ace1* architecture in *Anopheles* (Grau-Bové et al. 2021). The two are not exclusive, because mosquitoes can show cross-tolerance between high temperature and insecticides (Mack and Attardo 2024). CpL1a is best treated as a strong candidate for temperature-associated adaptive introgression with direct relevance to metabolic resistance, not as proof of thermal tolerance or of insecticide resistance alone.

On the same temperature gradient that favors QUI-like ancestry at CpL1a, QUI-like ancestry is progressively excluded at CpL2a (Fig. 5). The core signal centers on a *Yan* ortholog, an ETS-family transcriptional repressor of cellular differentiation (Webber et al. 2018), and the surrounding haplotype includes *takeout*-family genes, among them the ortholog of a *takeout*-like gene confirmed as a FOXO target in *Cx. pipiens* diapause (Sim et al. 2015), together with circadian and transporter genes. Diapause is therefore a plausible functional theme, because it declines in hybrids as the share of the genome from *Cx. quinquefasciatus* rises (Wilton and Smith 1985) and depends strongly on the maternal species in *Cx. pipiens* × *Cx. quinquefasciatus* crosses (Meuti et al. 2015). It does not establish the selected phenotype: maximum summer temperature is not the photoperiodic cue for diapause, most candidate SNPs are noncoding, and the candidate SNPs do not distinguish PIP from MOL. CpL2a is best read as selection for a non-QUI regulatory haplotype at a candidate diapause locus, and its interpretation is preliminary because it rests on fewer SNPs and lower reference differentiation than CpL1a. Its larger contribution is the evidence for exclusion: introgression is filtered out at individual loci even at the warm sites where *Cx. quinquefasciatus*-like ancestry rises across the rest of the genome.

Urbanization acts on this hybrid zone differently, not locus by locus, but through *molestus* ancestry as a whole. Genome-wide *molestus*-like ancestry rises with impervious surface independently of temperature, while *Cx. quinquefasciatus*-like ancestry shows no such association. This is the pattern the *molestus* bioform’s ecology predicts. It breeds in below-ground and human-made environments, persists year-round without obligate diapause, and prefers human over avian hosts (Bell et al. 2024; Fritz et al. 2015; Haba and McBride 2022), traits suited to densely built habitat. Urban sites therefore appear to either favor or concentrate the lineage best suited to them.

*Molestus*-like ancestry also rises toward the warmer end of the corridor, independently of impervious surface, and one proposed explanation is winter survival. Form *molestus* cannot diapause, which is thought to confine it to urban habitats in temperate regions (Aardema et al. 2022). Milder winters could therefore favor *molestus* aboveground (Haba and McBride 2022). Across our 24 sites, however, maximum summer temperature barely tracked winter temperature (r = −0.26 with the minimum temperature of the coldest month and r = 0.33 with the mean temperature of the coldest quarter). Genome-wide *molestus*-like ancestry followed neither winter measure (P = 0.58 and 0.32), consistent with the absence of a latitudinal trend in *molestus* introgression across the western Palearctic (Haba et al. 2025).

CpL3a’s abrupt linkage boundaries, breakpoint evidence, and uniform differentiation identify an 11.6 Mb inversion, the same inversion 3q that Cebrián-Camisón et al. (2026) found by sequencing 217 individual *Cx. pipiens* from southwestern Spain. There, every *molestus* carried the inverted arrangement on both chromosomes, and *pipiens* carried it as a polymorphism. The authors interpreted 3q as an ancestral polymorphism, older than the split between the forms, that *molestus* has fixed, and did not test the arrangement against urbanization. In our corridor, the inversion is *molestus*-like in every population, and *molestus*-like ancestry, both genome-wide and inside the inversion, is higher in more urban populations independently of temperature (Fig. 4, Fig. 5).

The rise of Δ with impervious surface at CpL3a does not by itself show selection on the inversion, because the inversion is a better marker of *molestus* ancestry than the rest of the genome. Our *molestus*-like ancestry estimate counts alleles private to one laboratory *molestus* colony. Across most of the genome, we expect wild *molestus* and *pipiens* to share much of their variation, as they do in Spain (Cebrián-Camisón et al. 2026), so many of those alleles are probably not fixed in wild *molestus*, and the genome-wide estimate would register only part of any real change in *molestus* ancestry. Inside the inversion, suppressed recombination keeps the two arrangements apart, so the panel’s alleles are mostly fixed differences and register the *molestus* arrangement almost in full. A rise in *molestus* ancestry that was the same everywhere in the genome would therefore still make the inversion’s estimate rise faster, as observed (2.4 times as fast as the genome-wide level).

Two explanations therefore fit the pooled data at CpL3a equally well. In the first, the inversion is a passenger: *molestus* ancestry is more common in cities for reasons that do not involve it, and the inversion marks that ancestry more faithfully than the rest of the genome. In the second, urban habitat favors the arrangement itself, and selection on it would also pull *molestus*-like ancestry across the rest of the genome in individuals that have not recombined it away. Pooled frequencies cannot distinguish the two, because both predict that the arrangement and genome-wide ancestry rise together. The second has a clear theoretical basis: inversions capture locally adapted alleles and shield them from recombination (Kirkpatrick and Barrett 2015), and although adaptive inversions usually assemble their content gradually (Schaal et al. 2022), one that evolved in a separate lineage can enter a new population already carrying it (Kirkpatrick and Barrett 2015). Form *molestus*, which carries the arrangement fixed in Spain (Cebrián-Camisón et al. 2026), has been evolving in human-modified environments for more than a thousand years (Haba et al. 2025), so the arrangement could enter the corridor pre-adapted. Separating the two requires genotyping the arrangement in individuals and setting it against each individual’s genome-wide ancestry. Because *pipiens* carries the arrangement too (Cebrián-Camisón et al. 2026), that test is feasible: if urban habitat favors the arrangement itself, urban individuals of mostly *pipiens* ancestry should carry it more often than rural ones. Whichever explanation holds, the inversion is the sharpest genomic marker of *molestus* ancestry in the corridor.

Several limitations bound these inferences, the most important of which is geographic. Maximum summer temperature is strongly collinear with position along the corridor, with 86% of its variation predicted by latitude and longitude alone. The design separates the thermal gradient from urbanization, but not from unmeasured variables that share the same geography, including proximity to the southern *Cx. quinquefasciatus* source (r = −0.75 with temperature), so the temperature results identify loci that track the thermal gradient without demonstrating that temperature is the causal agent. Replication in other metropolitan areas would strengthen these findings.

In addition, Pool-seq cannot phase ancestry, copy number, or inversions within individuals, and read depth is consistent with copy-number variation but is not a direct count of copies. Donor assignment relies on one laboratory reference pool per lineage (Table S9), with unequal colony histories and private-panel sizes, and mapping to a PIP-like assembly acts against CpL1a rather than for it by suppressing detection of QUI alleles. The Chicago-derived MOL reference may also not represent the *molestus* in Washington’s parking structures, because North American *molestus* populations from different cities differ (reviewed in Aardema et al. 2020). These features limit absolute ancestry estimates and leave CpL2a’s PIP-versus-MOL source unresolved, although environmental directions are robust to threshold choice and matched genomic nulls.

These limitations notwithstanding, the loci have implications for transmission and vector control. Because *Cx. quinquefasciatus* bridges avian and mammalian hosts (Andreadis 2012; Farajollahi et al. 2011) and *molestus* prefers human over avian hosts (Bell et al. 2024; Fritz et al. 2015; Noreuil and Fritz 2021), increasing temperature and urbanization may each move human-contact-associated ancestry into corridor populations, and each locus suggests a testable consequence. A copy-number-expanded CYP6 haplotype at CpL1a could alter metabolic resistance, and a diapause gene at CpL2a could affect seasonal activity. Common-garden assays of CYP6 expression, insecticide and heat tolerance, and diapause would test both. At CpL3a, the inversion carries gustatory and odorant receptors and could hold the host-preference differences between *molestus* and *pipiens* together as a linked unit (Bell et al. 2024; Fritz et al. 2015). Whether *molestus* ancestry raises biting of humans in nature remains unresolved (Hamer et al. 2009; Huang et al. 2009; Kilpatrick et al. 2006; Kothera et al. 2020). Individual long-read genomes sampled over time would phase donor ancestry with the CYP6 duplication, resolve the CpL2a haplotype, and genotype the inversion. The same loci could also serve as surveillance markers read from the pooled DNA that arboviral surveillance already tests (CDC 2024; Kothera et al. 2019).

The separation of the urban signal from the temperature-associated loci also matters beyond this corridor. Temperature and urbanization are often coupled through the urban heat-island effect (Fyie et al. 2023), which makes their evolutionary effects difficult to distinguish. The two-metropolitan design breaks that coupling: CpL1a and CpL2a follow the thermal gradient but not impervious surface, while the association of *molestus*-like ancestry with impervious surface, genome-wide and at CpL3a, holds with temperature in the model, so the urban response is not simply a thermal response inside a city. Nor will warming move a uniform *Cx. quinquefasciatus* genome northward, because CpL1a and CpL2a respond in opposite directions on the same gradient. Continental sampling already documents *Cx. quinquefasciatus* ancestry expanding across mid-latitudes (Saarman et al. 2025), and hybridization with *Cx. pipiens* f. *pipiens* could carry diapause into *Cx. quinquefasciatus* and let it move north (Aardema et al. 2022). Ultimately, our results indicate that climate change and urban development may reassemble trait combinations from ancestry already in contact, rather than shift the ranges of intact lineages.

## Acknowledgments

We thank Parker Grebe, Isabel Peñafiel, Ellen Osterman, and Nicole Guzman for help with mosquito collection and identification. Thank you to those who assisted us with site access: Al Greene (General Services Administration, retired), Arnaud Martin, Caroline Spiccioli, Rob McCann, David Serre, Josh Stift, Allison Dineen, Brenda Lee, Patuxent Research Refuge (USFWS), Smithsonian Environmental Research Center (SERC), the Smithsonian Institution, and University of Maryland Research and Education Centers at Beltsville, Upper Marlboro, and Clarksville. We also thank Brian Prendergast, who provided valuable insights into vector management practices in the State of Maryland. Maryland Genomics (Baltimore, MD) provided Illumina sequencing services.

This work was funded by NIH R01AI125622A to MLF. This work was also funded by USDA Extension Implementation Grant SZUSDA52O8900UMD to MLF. This publication was supported by the Centers for Disease Control and Prevention of the U.S. Department of Health and Human Services (HHS) as part of a financial assistance award (CDC-TEC 6NU50CK000633-01-01) totaling $21,600 with 100 percent funded by CDC/HHS. The contents are those of the author(s) and do not necessarily represent the official views of, nor an endorsement, by CDC/HHS, or the U.S. Government.

## Data Accessibility and Benefit-Sharing

### Data Accessibility Statement

Raw Pool-seq reads for the 27 pools (24 field and 3 reference) have been deposited in the NCBI Sequence Read Archive under BioProject PRJNA1551493 and will be released on publication. Site coordinates (decimal degrees) are given in Table S1 and, with the collection period, in the BioSample records linked to that BioProject. The derived inputs to the analyses (allele counts and frequencies, filtered variant calls, BayPass output, the lifted JHB2020 annotation, and individual-level ddRADseq genotypes) are available from Dryad (https://doi.org/10.5061/dryad.gmsbcc367; private for peer review, reviewer link provided at submission; public on publication). The *Culex pipiens* reference assembly idCulPipi1 is available from ENA under project PRJEB67967. Analysis code is available at https://github.com/BGregory98/culex-pipiens-hybrid-zone-genomics, and includes the per-site environmental predictors at full precision together with the metropolitan-area assignment used in every locus-level model, so that the within-area analyses reported here can be reproduced without rounding.

### Author contributions

B.G.: conceptualization, methodology, software, formal analysis, investigation, data curation, visualization, and writing of the original draft. A.A.-B.: investigation and resources, comprising the field collections and ddRADseq library preparation underlying the individual-level dataset, and review and editing of the manuscript. K.L.B.: formal analysis and data curation for the individual-level ddRADseq dataset, comprising read processing, variant calling and ancestry estimation, and review and editing of the manuscript. M.L.F.: conceptualization, supervision, resources, funding acquisition, and review and editing of the manuscript.

### Conflict of interest

The authors declare no competing interests.

## Notes

https://github.com/BGregory98/culex-pipiens-hybrid-zone-genomics

https://doi.org/10.5061/dryad.gmsbcc367

### Competing Interest Statement

The authors have declared no competing interest.

### Summary of Updates

Revised following submission to Molecular Ecology: text shortened and reorganized, CpL2a statements corrected, windowed f3 analysis removed from the supplement. Conclusions unchanged

