## Supplementary for "Urban enrichment of *molestus* ancestry and opposing temperature clines in a metropolitan *Culex pipiens* hybrid zone"

**Contents**

Supplementary Methods

Supplementary Results

Supplementary Figures S1–S10

Supplementary Tables S1, S2, S4, S6 and S9 (Tables S3, S5, S7 and S8 are provided as separate worksheets in Supplementary_Data.xlsx)

Supplementary References

### Supplementary Methods

#### Field collection, species identification and pool construction

At each of the 24 field sites, a single CDC gravid trap (John W. Hock Co., model 1712) baited with alfalfa-infused oviposition attractant was deployed overnight once per month in June, July, and August of 2023 and 2024, with additional trapping at a small number of 2024 events to meet pooling targets. Mosquitoes were cold-anesthetized at −20 °C, sorted within 24 h and stored at −80 °C. Specimens were sorted morphologically to *Culex*, then identified by a rapid abdomen-maceration DNA preparation adapted from Gloor and Engels (1992) followed by the ribosomal DNA (ITS) PCR assay of Crabtree et al. (1995), which assigns individuals to *Cx. restuans*, the *Cx. pipiens* assemblage, or *Cx. salinarius* by band size. Only abdominal tissue entered the PCR, so that thoraces were preserved for sequencing, and all PCR-confirmed *Cx. pipiens* assemblage females were retained.

Each field pool combined thoracic tissue from 40 females, balanced across collection dates to reduce temporal bias; the three reference pools were built identically from 40 thoraces each (collection coordinates in Table S9). Genomic DNA was extracted with the DNeasy Blood & Tissue Kit (Qiagen 69504), with two extraction blanks as contamination controls, and quantified on a Qubit 4 Fluorometer (ThermoFisher Scientific Q33238). Libraries were prepared and sequenced by Maryland Genomics on an Illumina NovaSeq S4 (150 bp paired-end), targeting approximately 20× mean coverage per pool and yielding approximately 1.37 billion read pairs (approximately 410 Gbp) across the 27 pools.

#### Read processing, alignment, variant calling and software versions

Reads were adapter- and quality-trimmed with Trimmomatic v0.33 (Bolger et al. 2014) and aligned with BWA-MEM2 v2.2.1 (Vasimuddin et al. 2019) to the *Cx. pipiens* reference assembly idCulPipi1 (Hesson et al. 2025). Duplicates were removed with Picard v3.4.0 and alignments were filtered to properly paired reads with mapping quality of at least 10. Pileups were generated with SAMtools v1.16 (mpileup -B -q 10 -Q 20 -d 500 -a; Danecek et al. 2021); read-pair overlap removal is enabled by default in this version, so bases sequenced twice within a fragment contribute a single observation. Variants were called across all 27 pools with PoolSNP v1.0.1 (Kapun et al. 2020) with min-cov 10 (every pool at depth ≥ 10), max-cov 0.99 (no pool above its own per-chromosome 99th depth percentile), min-count 20, min-freq 0.05, miss-frac 0.2 and base-quality 20; sites with any missing pool were then removed, yielding 11,629,890 biallelic SNPs with complete allele-frequency data across pools. The two depth filters define the callable genome on which every analysis rests: 155.1 M positions, 29.1% of the assembly, pass them in all 27 pools. R analyses used R 4.6.1 with poolfstat 3.1.0, terra 1.9-34, sf 1.1-2 (Pebesma 2018), lme4 2.0-6, lmerTest 3.2-1, glmmTMB 1.1.14, rtracklayer 1.72.0 and GenomicRanges 1.64.0 (full list in the code repository's renv.lock); the cluster aggregation steps used R 4.5.3.

**Environmental predictors**

Site-level predictors were extracted within a buffer of 1 km diameter around each site using terra (Hijmans 2026), a span close to the mean dispersal distances of 1.15–1.33 km measured for *Culex* (Ciota et al. 2012; Hamer et al. 2014). Percent impervious surface was taken from the National Land Cover Database 2021 (30 m; Dewitz 2023) and tree canopy from the companion USDA Forest Service NLCD tree canopy cover layer for 2021 (30 m; USDA Forest Service 2023), human population density from the WorldPop UN-adjusted population density surface (30 arc-second; WorldPop and Bondarenko 2020), as the natural log of persons $km^{-2}$, water-table depth from the simulated long-term average surficial groundwater system of the contiguous United States (250 m; Zell and Sanford 2020), and elevation from the USGS National Elevation Dataset (30 m; Gesch et al. 2018), alongside three BIOCLIM variables from CHELSA V2.1 (30 arc-second, 1981–2010; Karger et al. 2017, 2021): bio05, maximum temperature of the warmest month (maximum summer temperature here and in the main text); bio07, temperature annual range; and bio18, precipitation of the warmest quarter. The eight predictors were retained from a larger exploratory set of climatic, hydrological and land-cover layers after pruning for redundancy; they were not chosen a priori to separate the two gradients, although between them they represent both. Elevation was retained as a proxy for winter cold, and winter minimum temperature and mean temperature of the coldest quarter were dropped for being more than 90% correlated with it. Correlations within each gradient (temperature and urbanization) are much stronger than between them: maximum summer temperature with temperature annual range at $r = 0.84$ and percent impervious surface with tree canopy at $r = -0.90$, against $r = 0.26$ for maximum summer temperature with percent impervious surface (Fig. S1). This is why we interpret each association as a response to its gradient rather than to any single predictor.

**BayPass model settings**

BayPass v3.1 (cited in the main-text Methods) was run in two stages on the 24 field pools. The core model estimated the population covariance matrix Ω and calibrated $XtX_{st}$ statistics. The standard covariate model was then run in default importance-sampling mode using Ω and centered, scaled covariates (elevation, impervious surface, tree canopy, water-table depth, population density, maximum summer temperature, temperature annual range, precipitation of the warmest quarter). Both models used 10 pilot runs of 100 iterations each (the BayPass defaults), a 5,000-iteration burn-in, and 25,000 sampling iterations thinned every 25 (-nval 1000 -thin 25 -burnin 5000). $XtX_{st}$ significance thresholds were calibrated with a pseudo-observed dataset (POD) of 100,000 neutral SNPs simulated under the estimated Ω covariance and ($a_{\pi}$, $b_{\pi}$) parameters; the 99th and 99.9th percentiles of the POD $XtX_{st}$ null distribution ($XtX_{st}$ = 71.1 and 106.6, respectively) were the minor- and major-outlier thresholds, and Bayes factor thresholds of 10 and 30 dB (Bayes factors of 10 and 1,000) defined minor and major outliers for environmental association. Across the genome, 1,510 SNPs were significant outliers on both $XtX_{st}$ and a Bayes factor for at least one predictor. Because outlier positions under the permissive chain definition occur at roughly one per 5 kb genome-wide, the spatial criterion removed only 14% of these, leaving the 1,299 candidate SNPs (1.4% of positions retained), and the dual-outlier requirement is the operative filter.

**Annotation liftover, gene-content surveys and candidate-SNP attribution**

The *Cx. quinquefasciatus* JHB2020 annotation was projected onto the *Cx. pipiens* reference with Liftoff v1.6.3 (-copies -polish -infer_genes -infer_transcripts; Shumate and Salzberg 2021), and the polished GFF3 was used in all annotation-based analyses. Protein-coding genes overlapping each candidate interval were extracted with rtracklayer (Lawrence et al. 2009) and GenomicRanges (Lawrence et al. 2013) and classified into candidate families by regex matching on the GFF3 description field (cytochrome P450, glutathione S-transferase, carboxylesterase, ABC transporter, heat shock protein, voltage-gated ion channel, chemosensory receptor, acetylcholinesterase, cuticle/chitin, circadian and reproductive families). Genes of interest were queried against SwissProt by BLASTP with default settings to confirm orthology. At CpL2a this confirmed CQUJHB014704 as the *Yan* ortholog (ets DNA-binding protein pokkuri), CQUJHB002678 as *takeout* (the *takeout*-like FOXO target validated by Sim et al. (2015), CPIJ014239, corresponds instead to CQUJHB008894, an unannotated member of the same cluster whose protein is identical to it except at the final residue), and three ABC-family genes at approximately 14.3 Mb (CQUJHB009840 and CQUJHB007929, multidrug resistance-associated protein 1 and 1-like; CQUJHB002428, canalicular multispecific organic anion transporter).

To localize each region's BayPass signal, the clustered candidate SNPs (Methods, *Genome–environment association with BayPass*) were attributed to protein-coding genes of the lifted annotation. For every gene overlapping a candidate region we counted candidate SNPs within the gene span extended by a 2 kb flank on each side and, separately, within the per-gene union of CDS exons across all isoforms, and normalized each by the corresponding length (candidate SNPs per kb of gene plus flank, and per kb of CDS). We also recorded the maximum single-SNP $XtX_{st}$ within the gene-plus-flank interval and, among the gene-body candidate SNPs, the fraction whose most strongly associated predictor was the locus's nominal predictor (maximum summer temperature for CpL1a and CpL2a, percent impervious surface for CpL3a) and the fraction assigned to that predictor's broader category (temperature = maximum summer temperature + temperature annual range; urbanization = percent impervious surface + tree canopy + population density). The 2 kb flank captures proximal regulatory sequence without crossing into adjacent genes at the assembly's gene density (approximately 1 gene per 25 kb); distal regulatory elements are not recoverable by proximity-based attribution. The analysis was applied as a targeted localization at the six CpL1a CYP6 paralogs ranked against all genes in the CpL1a interval, as a discovery ranking of all genes in the 500 kb CpL2a core by gene-body candidate-SNP density, and as an exploratory ranking across the CpL3a inversion span, where the inversion rather than any gene is the unit of selection and gene-level results are context only.

**Individual-level ddRADseq genotyping**

Pooled sequencing cannot by itself distinguish a pool of admixed individuals from a pool of unmixed ones. We therefore drew on an independent individual-level ddRADseq dataset. Adult *Culex* were collected from 15 sites across the Maryland and Washington, DC portion of the study region between June 2019 and October 2020, together with laboratory colony material that included individuals from the source populations of all three reference pools: Vero Beach, FL (QUI), Orono, ME (PIP) and Calumet, IL (MOL).

Libraries were prepared from 616 individuals following Parchman et al. (2012) and Gompert et al. (2014), as described in Bell et al. (2024): genomic DNA was digested with EcoRI and MseI, ligated to adaptors carrying 8–10 bp individual barcodes, amplified with standard Illumina primers, size-selected to 300–400 bp on a BluePippin (Sage Science) and sequenced as 100 bp single-end reads on an Illumina NovaSeq SP at the North Carolina State University Genomic Sciences Laboratory. PhiX reads were removed with bowtie2 (Langmead and Salzberg 2012); remaining reads were demultiplexed with custom Perl scripts and aligned with BWA-MEM (Li 2013) to the *Cx. quinquefasciatus* JHB2020 assembly (GCA_015732765.1), an established practice for *Cx. pipiens* reads (Yurchenko et al. 2020; Noreuil and Fritz 2021; Bell et al. 2024). Variants were called with bcftools (Danecek et al. 2021), excluding indels, with the posterior probability that a site is invariant set to 0.05 and the prior on SNPs to 0.001. Forty-three low-coverage individuals were removed, leaving 573. SNPs were retained where total coverage was at least twice the number of individuals, mapping quality was at least 30, alternate-allele frequency fell between 0.05 and 0.95, base-quality, mapping-quality and read-position rank-sum statistics were below 2.5 in absolute value, and data were present for at least 95% of individuals; variants with coverage more than two standard deviations above the mean were discarded. This yielded 7,258 SNPs at a median depth of 8× per individual. Genotype probabilities were estimated in entropy version 1 (Gompert et al. 2014; Shastry et al. 2021) under a K = 2 admixture model, as two independent MCMC chains of 20,000 steps with a 5,000-step burn-in and thinning every 10 steps. Posterior mean genotypes, computed from the posterior genotype probabilities, were averaged across chains and ordinated by principal component analysis of the 553 field and reference individuals, excluding two colony populations from outside the study region (Fig. S4).

**Donor-private ancestry panels and the per-population donor-ancestry test**

The test asks whether locus-specific donor ancestry exceeds a population's genome-wide donor ancestry. Because allele frequencies in an admixed population are a linear combination of its sources' frequencies at every SNP (Patterson et al. 2012; Peter 2022), donor-ancestry proportions under a pure mixture of unmixed individuals are constant across the genome within a pool; under within-individual locus-specific introgression with environmentally structured selection, the focal locus departs from the genome-wide value and tracks the environmental predictor across pools.

For each reference lineage, a lineage-private panel was defined as the SNPs at which that reference carried the alternate allele at frequency ≥ 0.95 while both other references carried it at ≤ 0.05, with reference frequencies computed from the VCF under the same depth filter (≥ 10) used elsewhere. Panels were drawn from all SNPs within an interval rather than from the BayPass candidate set, and so are selected on reference frequencies alone; overlap with the candidate set is minimal (3 of 123 QUI-private SNPs at CpL1a, none of the 3,762 MOL-private SNPs there, and none of the 6 QUI-private or 30 MOL-private SNPs at CpL2a), so unlike the candidate SNPs these markers are not ascertained on environmental association. For each pool *i* and focal locus, $\alpha_{locus,i}$ was the mean pool alternate-allele frequency across panel SNPs within the focal interval and $\alpha_{genome,i}$ the same quantity across panel SNPs outside the union of the three candidate regions. The per-population difference $\Delta_i = \alpha_{locus,i} - \alpha_{genome,i}$ estimates locus-specific donor ancestry net of genome-wide donor ancestry, and has expectation zero at every locus under pure mixture provided that the panel marks donor ancestry equally well inside the locus and across the genome. Because this test admits a SNP to a panel only where every reference pool has depth ≥ 10, its panels are slightly smaller than those counted in Table S4: 118 QUI-private SNPs at CpL1a, 3,107 MOL-private SNPs at CpL3a and, for the non-QUI deficit at CpL2a, 49 QUI-private SNPs across the 2.5 Mb haplotype block (Table S4 counts 6 within the 500 kb core); the genome-wide panels outside the three loci hold 5,396 QUI-private and 112,641 MOL-private SNPs.

Donor identity was QUI at CpL1a and MOL at CpL3a. At CpL2a the PIP and MOL references are not separated at the candidate SNPs that distinguish QUI from PIP (mean $|\Delta p|$ = 0.11 across those 27 SNPs) and so cannot serve as discriminating donors, so we used the QUI-private panel with the deficit estimator 1 − $\alpha_{QUI}$ (equivalently $\alpha_{PIP}$ + $\alpha_{MOL}$, the combined non-QUI ancestry), testing whether warmer pools carry more non-QUI ancestry at CpL2a without committing to a PIP-versus-MOL donor identity. For each locus and panel we tested a marginal hypothesis, that the median of Δ across the 24 field pools exceeds zero (one-sided Wilcoxon signed-rank test), and a joint hypothesis, that Δ regresses positively on the relevant predictor (maximum summer temperature for CpL1a and CpL2a, percent impervious surface for CpL3a; one-sided linear regression). One-sided empirical-percentile P values for both statistics were computed against the null distribution of 10,000 width-matched random regions described next. Because a panel can mark donor ancestry more faithfully inside an inversion, where suppressed recombination preserves fixed differences between arrangements, than across the freely recombining genome, $\alpha_{locus}$ was also regressed on $\alpha_{genome}$ and the locus's predictor together, on the linear and on the log scale, and each term was read with the other in the model.

**Genome-wide null distribution for locus-level diagnostic statistics**

For each candidate locus, 10,000 random regions of the same width as the focal interval (2.5 Mb for CpL1a and CpL2a; 11.62 Mb for the CpL3a inversion span) were sampled uniformly along the three autosomal contigs, weighted by chromosome size, with rejection of any region overlapping the union of the three candidate intervals. At each region we computed five statistics on the field-pool allele-frequency matrix restricted to that region's SNPs: (i) PC1 % variance; (ii) the absolute Pearson correlation between pool scores on PC1 and pool mean $F_{ST}$ to the donor reference; (iii) the across-pool variance in mean $F_{ST}$ to the donor reference; (iv) the median across pools of the donor-private panel deviation Δ; and (v) the slope of Δ on the relevant predictor. The donor reference was QUI for CpL1a, QUI for CpL2a (read on the QUI-versus-non-QUI axis) and MOL for CpL3a. Mean per-population $F_{ST}$ to a reference was the per-SNP estimator of Reynolds et al. (1983), $(p_{pool} - p_{ref})^2 / (\bar{p}(1 - \bar{p}))$, averaged across SNPs. One-sided empirical-percentile P values for each focal-locus statistic were read from these distributions.

**Differentiation-matched null distribution**

The width-matched null holds region size constant but not the differentiation on which candidate regions were selected. We therefore built a second null whose comparison regions were additionally matched on mean $XtX_{st}$. Per-SNP calibrated $XtX_{st}$ values from the BayPass core model were joined to the allele-frequency matrix by marker index; 5,000 random regions of each focal width were drawn from the three chromosomes in proportion to length, excluding the candidate intervals and 8 Mb on either side of each, beyond the distance at which flank SNPs reach baseline correlation with the candidates at every locus (main text Fig. 4), and retained when their mean $XtX_{st}$ fell within a tolerance of the focal region's, widening from ±10% until at least 200 regions were retained. The same statistics were computed in each.

Two limitations apply. First, no random 2.5 Mb region attains CpL1a's mean $XtX_{st}$ of 50.7; the most differentiated available reach 36.7, against a width-only median of 23.0, so at CpL1a the comparison set is the most differentiated available rather than a matched one. CpL2a and CpL3a match within ±25% and ±10% respectively. Because a 40 Mb stretch of chromosome 1 (74–113 Mb) is the only broad region of the genome as differentiated as the candidate loci, nearly all retained regions lie there (all 291 at CpL1a, all 541 at CpL3a and 232 of 239 at CpL2a), so the comparison is with the most differentiated unlinked sequence available rather than with a sample of the whole genome. Second, the null distribution of the Δ–environment correlation is not centered on zero, because Δ subtracts a genome-wide mean that is itself environmentally structured: regions with small panels, whose ancestry is imprecise, return systematically shifted correlations, and within retained regions the correlation rises with panel size ($r = 0.14$, 0.44 and 0.41 at the three loci). The sign and size of that shift depend on the panel and the window width, so it must be measured for each panel and locus rather than assumed (Supplementary Results, *Locus statistics against a differentiation-matched null*). We therefore report each focal region's rank among retained regions rather than a calibrated percentile.

**Haplotype-block diagnostics**

For each candidate interval we tested whether the region behaves as a single segregating haplotype block using three pre-specified statistics on the per-population allele-frequency matrix restricted to region SNPs: PC1 % variance, the absolute correlation between PC1 scores and mean per-population $F_{ST}$ to the donor reference, and the across-pool variance in that $F_{ST}$. Per-population alternate-allele frequencies were extracted from the PoolSNP VCF with VariantAnnotation (Obenchain et al. 2014; vcfR v1.16.0, Knaus and Grünwald 2017, for the genome-wide null regions); SNP × pool cells with depth < 10 were masked, SNPs with mean across-pool frequency outside [0.05, 0.95] were dropped, and rows with remaining missingness were excluded. PCA was run on the 24 field pools by singular value decomposition (prcomp, centered, unscaled), with reference pools excluded. Donor references were QUI for CpL1a, QUI for CpL2a (with the diagnostic read on the QUI-versus-non-QUI axis, for the reason given above) and MOL for CpL3a. Each statistic was compared with the genome-wide null of 10,000 width-matched regions, with one-sided P values in the direction predicted by the haplotype-block hypothesis.

**Windowed and local PCA scan**

To test whether the candidate regions, and CpL3a in particular, could be detected without reference to the genome scan or to any environmental variable, we repeated the haplotype-block PCA in non-overlapping windows along the three chromosomes. This is the pooled counterpart of the windowed PCA with which Cebrián-Camisón et al. (2026) detected inversions in individually sequenced *Cx. pipiens*. SNPs were filtered as for the haplotype-block diagnostics (depth ≥ 10 in every field pool; mean field-pool frequency within [0.05, 0.95]), and windows retaining fewer than 200 SNPs were dropped, leaving 1,066 windows of 500 kb and 534 of 1 Mb. In each window, every SNP was centered on its field-pool mean and PCA was run on the 24 field pools. Windows were ranked genome-wide by the percentage of variance on PC1, a window counting as near a candidate locus if it lay within 2 Mb of the candidate interval. We recorded the percentage of variance on PC1 and the PC1 score of each pool, oriented so that positive scores indicate lower mean $F_{ST}$ to MOL. To find runs of windows sharing one local structure, we compared windows by local PCA (Li and Ralph 2019). Each window was summarized by its top two principal components, with eigenvalues scaled to sum to one, and the distance between two windows was the Frobenius norm of the difference between these rank-two summaries. Classical multidimensional scaling reduced the distances among all windows to two axes. An inversion whose frequency varies among pools should appear as a contiguous block of windows sharing an extreme value on one axis. Because each point is a pool, the scan cannot assign karyotypes and cannot detect an inversion that segregates at a similar frequency in every pool.

**Linkage disequilibrium analysis**

To test whether the candidate SNPs at each locus co-segregate as a block, and whether block-scale linkage extends beyond them, we computed pairwise allele-frequency correlations ($r^2$) across the 24 field pools for five SNP sets per locus: (i) within-cluster BayPass candidate SNPs; (ii) all within-cluster SNPs passing filters, subsampled to 5,000 where more numerous; (iii) pairs between a within-cluster candidate SNP and one of up to 3,000 randomly selected non-cluster SNPs within 3 Mb of the cluster, which measures the spatial extent and boundary sharpness of the block; (iv) pairs between the candidate SNPs at this locus and those at the other two loci, which measures environmentally driven covariation independent of physical linkage; and (v) a chromosomal baseline of 2,000 randomly selected non-cluster SNPs on the same chromosome. Values were binned by pairwise genomic distance from < 1 kb to > 20 Mb.

**Callable-position read-depth analysis**

A position was defined as callable if it passed PoolSNP's depth filters: per-population depth ≥ 10 across all 27 pools (min-cov 10) and not exceeding the per-population, per-chromosome 99th percentile (max-cov 0.99). Restricting depth analysis to callable positions puts the read-depth and allele-frequency analyses on the same genomic substrate. Per-position callable depths were extracted from the genome-wide mpileup in a two-pass streaming pipeline (pass 1, per-population and per-chromosome 99th-percentile cutoffs; pass 2, per-population depth at positions passing both filters), intersected with the lifted CDS annotation using BEDTools (Quinlan and Hall 2010), and summed within genes to give total depth for 13,741 genes × 24 field pools; the genome-wide sum of callable depth per pool served as a sequencing-effort denominator. Genome-wide, 155.1 M positions (29.1% of the assembly) were callable.

A baseline model (M0) regressed per-gene total depth on z-scored maximum summer temperature (bio05_z in the model formulas) with log genome-wide callable depth as an offset and a random intercept for gene:

total_depth ~ bio05_z + offset(log(genome_callable_depth)) + (1 | gene_id)

The model was fit in glmmTMB v1.1.14 (Brooks et al. 2017); Pearson dispersion under a Poisson family (201.5) far exceeded one, so the model was refit under a negative-binomial (nbinom2) family. The offset absorbs sequencing-effort differences and the gene intercept absorbs gene-level depth heterogeneity from length, GC content and mappability. Because the bio05_z fixed effect is fit across all 13,741 genes, any component of among-pool mapping efficiency that covaries with maximum summer temperature (for example reduced mapping of QUI-derived reads at the warm sites where QUI ancestry is elevated) is absorbed into M0; per-pool mapping rates in fact do not covary with maximum summer temperature (β = $-2.1 \times 10^{-3}$, P = 0.59). A focal-interaction model (M1) added a focal-gene indicator and its interaction with bio05_z:

total_depth ~ bio05_z * is_focal + offset(log(genome_callable_depth)) + (1 | gene_id)

with is_focal = TRUE for the six CpL1a CYP6 paralogs (CQUJHB001163, CQUJHB008986, CQUJHB009148, CQUJHB017007, CQUJHB009005, CQUJHB007871) in the CpL1a model and for the CpL2a *Yan* ortholog (CQUJHB014704) in the CpL2a model. The interaction tests whether the focal genes deviate from the genome-wide depth–temperature slope, with the remaining approximately 13,740 genes as an in-model null. Reference pools were excluded from both models.

To measure cluster depth in the reference pools, the same callable-depth products were re-aggregated with all 27 pools retained; no new alignment or pileup was involved. A position is callable only if it has depth ≥ 10 in every one of the 27 pools, so the 21,361,387 callable positions that fall within the coding sequence of the 13,741 genes are the same positions in the field and reference pools alike. The reference ratios are therefore directly comparable with the field ratios, and because any position at which one lineage drops out is excluded for every pool, the comparison reflects copy number rather than allele dropout. Depth was expressed exactly as for the field pools: callable-position depth per gene divided by the gene's callable-position count, then by the pool median across genes, so that 1.0 is a gene of typical depth for its pool; the cluster ratio is the mean over the six paralogs. Genome-wide callable depth confirms comparable sequencing of the references (4.91, 4.52 and 5.32 billion callable-position reads for QUI, PIP and MOL, against 3.77 to 5.17 billion across field pools). Implied duplication frequencies follow from a diploid baseline, on which a ratio of 1 + f corresponds to a duplication on a fraction f of chromosomes; we report these as descriptions of each colony rather than as wild-population parameters.

**Coding consequences, phylogenetic placement, microsynteny and consensus sequences of the CpL1a CYP6 paralogs**

The coding consequence of every CDS candidate SNP at the six paralogs was predicted with predictCoding in VariantAnnotation (Obenchain et al. 2014), using transcript models built from the lifted annotation with GenomicFeatures (Lawrence et al. 2013). Each biallelic SNP was classified as synonymous, nonsynonymous or nonsense against the transcript in which it fell. Pool-seq data are unphased, so consequences were assigned to single substitutions; where two candidate SNPs fell in the same codon (codon 503 of the CYP6AA7-like paralog) each was scored independently against the reference codon. Because idCulPipi1 represents the *Cx. pipiens* background, the alternate allele is the candidate introgressed state, and a nonsynonymous SNP was classified as QUI-diagnostic when its alternate allele was near-fixed in QUI and absent from MOL.

For phylogenetic placement, coding sequences from the JHB2020 and lifted annotations were translated, queried against UniProtKB by BLASTP, and combined with a curated panel of 14 representative mosquito P450 proteins. Proteins were aligned with MAFFT v7.526 (L-INS-i; Katoh and Standley 2013); alignments made with G-INS-i or trimmed with trimAl -automated1 (Capella-Gutiérrez et al. 2009) gave identical topologies. Trees were inferred with IQ-TREE 3.1.1 (-st AA -m MFP -alrt 1000 -B 1000). Top matches from P450Atlas (Gront et al. 2025; v1.3.0, accessed April 10, 2026) were recorded as descriptive “-like” labels. For microsynteny, chromosome 1 of the *Cx. pipiens* (OZ004311.1) and JHB2020 (CM027410.1) assemblies were aligned with minimap2 v2.30 (-x asm10; Li 2018); because they aligned in opposite orientation, JHB2020 coordinates were reversed for local comparison, and alignment links were drawn from primary-filtered hits (length ≥ 1,000 bp, MAPQ ≥ 5).

Whole-gene nucleotide and concatenated CDS consensus sequences for each paralog were generated from the filtered BAMs of all 33 sequenced pools (the 27 analyzed here and six further reference and colony pools) by extracting the reference interval, calling variants with bcftools mpileup/call, filtering to depth ≥ 5 and QUAL ≥ 20, masking low-depth positions and applying bcftools consensus. CDS consensus sequences were translated with seqkit v2.13.0 (Shen et al. 2024) and aligned per gene with MAFFT (--localpair --maxiterate 1000; Katoh and Standley 2013). Pairwise percent identity was computed site by site, excluding gaps and ambiguous characters, and linear mixed models of percent identity on maximum summer temperature, reference and their interaction were fit with random intercepts for gene and sample.

**Allele-frequency clines and reference structure at CpL1a and CpL2a**

For the candidate SNPs within each cluster, a per-SNP binomial GLM of allele count on maximum summer temperature was fit across the 24 field pools (cbind(alt, ref) ~ temperature, logit link), with per-pool allele counts taken from the PoolSNP read counts. At CpL1a each SNP was polarized to the QUI–MOL axis, the slope signed so that a positive coefficient is an increase in the QUI-like allele with temperature, and the set restricted to SNPs effectively fixed between QUI and MOL ($|\Delta p| \geq 0.95$), the filter under which the polarized axis is well defined. At CpL2a the same procedure was applied on the QUI–PIP axis at a fixation threshold of $|\Delta p| \geq 0.70$, lowered because the references are less differentiated there. To place PIP on the QUI–MOL axis at the CpL1a candidate SNPs, pairwise $F_{ST}$ among the three reference pairs restricted to these SNPs was computed in poolfstat (pooldata.subset followed by computeFST), with block-jackknife standard errors from approximately 10 equal blocks; the CpL2a computation was identical.

**Indel calling and per-base depth profiling at CpL2a**

To assess PIP–MOL divergence in variant classes the PoolSNP pipeline does not capture, indels were called jointly from the three reference BAMs across chr2:11–16 Mb with bcftools mpileup (v1.9; -q 10 -Q 20 -d 500, FORMAT/AD,FORMAT/DP) and bcftools call (-mv, –

ploidy 1), the haploid model being appropriate for near-inbred colonies. Calls were filtered to indels within chr2:13.5–14.0 Mb, and PIP–MOL fixed-different indels were those at which one reference carried the reference allele (GT = 0) and the other the alternate (GT = 1). Per-base depth was extracted with samtools depth -a (-q 10 -Q 20) across the same interval, and zero-depth runs > 100 bp were identified per reference as mapping failures.

**Confirmation of inversion breakpoints**

DELLY was run on the main-pipeline alignments of the reference pools (BWA-MEM2 v2.2.1, -k 19 -M; Picard MarkDuplicates v3.4.0, REMOVE_DUPLICATES=true), taken before the proper-pair filter, since reads spanning an inversion breakpoint are by definition not in proper-pair orientation. Insert-size distributions were nearly identical across the three pools (mean 220–246 bp; 99th percentile 712–781 bp), so the same parameters were used for all. DELLY v1.7.3 (Rausch et al. 2012) was run in inversion-only mode (-t INV, -q 20) on each pool, retaining PASS calls with paired-end support ≥ 5, median MAPQ ≥ 30 and size between 50 kb and 20 Mb. Because Pool-seq does not resolve haplotypes, inversion frequencies cannot be estimated directly from these calls (Kapun et al. 2014), so they were used only to locate breakpoints.

The DELLY breakpoints (~149.4 and ~161.0 Mb) agree closely with inversion 3q characterized by Cebrián-Camisón et al. (2026) through windowed PCA in a Spanish *Cx. pipiens* population (~149–161 Mb on the same idCulPipi1 assembly), where all *molestus* specimens carried the inverted homokaryotype.

**Chemosensory gene content of the CpL3a inversion**

Because the JHB2020 annotation assigns gene families by description string and under-annotates insect chemosensory receptors, the CpL3a gene-content survey was repeated against a curated chemosensory reference assembled from Bell et al. (2024) and the manually curated chemosensory annotation, differentially expressed gene sets and sequence-divergent gene sets of Menna et al. (2026), lifted onto idCulPipi1 with Liftoff v1.6.3. The merged reference comprises 486 gene symbols, of which 482 carry lifted autosomal coordinates and a family assignment: 164 ionotropic receptors, 146 odorant receptors, 111 odorant-binding proteins and 61 gustatory receptors. Where a symbol lifted to multiple copies the leftmost was retained, so positional counts are per symbol and conservative; no symbol inside CpL3a lifted to more than one copy.

Counts within the inversion span between the DELLY breakpoints (149,402,438–161,021,387) were compared against the same 10,000 width-matched random regions used for the other locus-level diagnostics, giving one-sided upper-tail empirical P values for four statistics: total chemosensory genes; distinct tandem arrays (gene spans merged by single linkage at 100 kb, well above the approximately 25 kb mean intergenic spacing); per-family counts; and genes carrying a candidate SNP within the gene span ± 2 kb. Because candidate SNPs are concentrated at CpL1a and most random regions contain none, the between-region version of the last statistic mostly measures whether a region carries candidate SNPs at all. We therefore added a conditional test holding the 69 candidate SNPs inside the inversion fixed and randomizing only their positions, under a uniform null and under a circular shift of the observed position vector wrapping at the inversion boundaries, which preserves clustering and spacing and is the more conservative (10,000 permutations each). Both assume callable-site density is approximately even across the span.

**Geographic and metropolitan-area controls**

Each predictor was regressed on latitude and longitude to quantify its collinearity with position along the corridor. Sites were assigned to the Baltimore or Washington metropolitan area by great-circle distance to the nearer city center (Baltimore 39.2904° N, 76.6122° W; Washington 38.9072° N, 77.0369° W), giving 11 and 13 sites respectively (Table S1). Locus-level environmental models were refitted with a metropolitan-area term and with both gradients entered together, and separately within each area. Spatial autocorrelation of each predictor was measured as Moran's I on row-standardized inverse great-circle-distance weights. Spatial structure was modeled as a covariate rather than permuted, because no permutation of maximum summer temperature across the 24 sites reproduces its spatial autocorrelation (observed Moran's I = +0.494, against −0.210 to +0.224 across 2,000 unconstrained permutations), so a permutation null would be anti-conservative.

**Targeted analysis of insecticide-resistance target sites**

Because the genome scan retains only SNPs that are both differentiation outliers and environmentally associated, a site under selection from an unmeasured pressure cannot enter the candidate set. We therefore assembled a panel of canonical insecticide-resistance target and detoxification genes by matching gene descriptions in the lifted annotation: the voltage-gated sodium channel (*Vgsc*/*para*), acetylcholinesterase (*Ace-1*), GABA-receptor subunits (*Rdl*), the carboxylesterases and esterases of the Ester locus, and the glutathione S-transferases, to which we added the six CpL1a CYP6 paralogs and, as a labeled negative control, the sodium channel 60E gene. The panel comprises 63 genes and 99.0 kb of coding sequence, and allele counts were read for every coding position in it directly from the allele-count table, independently of the scan. Coding consequences were called against the lifted annotation, and per-pool frequency was the alternate-allele count divided by depth, set to missing below ten reads. Because a projected annotation cannot be assumed to preserve published residue numbering, a matching residue change was not by itself treated as a site call: the *Vgsc* candidate was reconciled to housefly numbering by global pairwise alignment (BLOSUM62) of the canonical *Culex para* translation to *Musca domestica* Vssc1 (UniProt Q94615), and only that gene was so reconciled, so the residue-match flags for *Ace-1* and *Rdl* in Table S3 are candidates rather than confirmed sites. Environmental associations were ordinary least-squares fits of per-pool frequency on each predictor. Two correlations involving *kdr* were tested against empirical nulls rather than against a standard significance test, because *kdr* is a strongly differentiated site and strongly differentiated sites correlate with ancestry as a matter of course. For the correlation between *kdr* frequency and genome-wide donor ancestry, the null was the same correlation computed at comparison SNPs matched to *kdr* on $XtX_{st}$ and on BayPass posterior mean allele frequency. For the correlation between *kdr* frequency and read depth at the CpL1a CYP6 cluster, the null was the same correlation computed at every other run of six adjacent genes in the genome and, for the region-level version, at the differentiation-matched 2.5 Mb windows described above. In every case the focal and null statistics were computed identically, and the empirical P value is the two-sided fraction of null values at least as extreme as the focal one.

### Supplementary Results

*Genome-wide context and environmental design*

#### Genome-wide diversity and differentiation from the reference pools

Per-pool diversity was indexed as the mean of $2p(1 - p)$ across all 11,629,890 SNPs, a relative index across pools of comparable coverage rather than a coverage-corrected estimate of nucleotide diversity. Mean expected heterozygosity across the 24 field pools is 0.2343 and varies little (0.2309 to 0.2381), but it rises with maximum summer temperature (adjusted $R^2$ = 0.45, $P < 0.001$) and is unrelated to impervious surface ($P = 0.66$). Genome-wide differentiation from each reference declines with maximum summer temperature (adjusted $R^2$ = 0.362 against PIP, 0.146 against QUI, 0.588 against MOL). Adding diversity to those models separates them: the decline against PIP does not survive (temperature $P = 0.72$) and is attributable to the diversity gradient, whereas the temperature terms against MOL and QUI remain significant, the decline against MOL persisting ($P = 0.0055$) and the decline against QUI reversing to an increase ($P < 0.001$), as Fig. S5B shows. Because of these diversity effects, the main text measures ancestry with lineage-private panels rather than $F_{ST}$. The reference pools differ markedly in diversity, with expected heterozygosity of 0.227 for PIP, 0.151 for MOL and 0.079 for QUI.

#### Background composition estimated from lineage-private panels

Genome-wide differentiation orders the references PIP 0.175 < MOL 0.268 < QUI 0.542, the inverse of their heterozygosities. Because $F_{ST}$ is deflated by within-population diversity, the most diverse reference shows the lowest $F_{ST}$ to any field pool for algebraic reasons, so this ordering cannot by itself identify the background composition. Re-estimated from lineage-private panels lying outside the three candidate intervals, mean $\alpha$ over the 24 field pools is 0.366 for

PIP, 0.158 for MOL and 0.108 for QUI at thresholds of ≥ 0.95 / ≤ 0.05; 0.350, 0.183 and 0.122 at ≥ 0.90 / ≤ 0.10; and 0.336, 0.202 and 0.136 at ≥ 0.85 / ≤ 0.15. PIP is the largest component at every threshold. These α values are indices of lineage-private allele frequency rather than ancestry proportions and are not expected to sum to one, and the PIP panel is much the smallest (440 SNPs outside the candidate intervals against 5,884 QUI-private and 116,508 MOL-private), so PIP ancestry is the least precisely estimated despite being the largest component. That asymmetry is expected: the reference with the highest heterozygosity is least often fixed for an allele absent from both others.

**Reference-pool provenance and its effect on the private panels**

All three reference pools derive from laboratory colonies, held for approximately 160 generations (MOL) or 20 to 30 generations (PIP and QUI) before extraction. Three consequences bear on the private panels: drift distorts a colony's allele-frequency spectrum; the references differ substantially in diversity; and because the reference assembly was built from a Swedish member of the PIP lineage, mapping efficiency declines with divergence from PIP, leaving QUI most exposed to allele dropout. These causes are entangled, but their joint effect on each panel is predictable in direction. PIP, the most diverse reference, has the fewest alleles fixed for a state absent from both others (444 SNPs). MOL's long colony history pushes alleles toward fixation beyond what its intermediate diversity predicts, while QUI dropout at the PIP-like assembly makes positions where QUI fails to map appear MOL-private, together producing 123,749 MOL-private SNPs. QUI, though the least diverse, loses private alleles to that same dropout and is intermediate (6,017). The > 250-fold range in panel size is thus expected from the biology and the sequencing design, and we do not compare panel sizes or absolute ancestry magnitudes between donors. This restriction does not touch the quantities the results rest on: α and Δ are each computed within a single donor's panel, so a donor's Δ–environment slope, its median Δ and its rank against the null are unaffected. What it rules out is reading, for example, a genome-wide QUI α of 0.108 and a MOL α of 0.158 as showing that the corridor carries more MOL than QUI ancestry, because the two panels are attenuated by different and unknown amounts. None of the three effects is expected to correlate with the environmental predictors across sites, so the direction and significance of each ancestry–environment relationship is robust to them; reference-mapping bias in particular acts against the CpL1a result, since it suppresses detection of the QUI alleles whose increase is reported.

**Temperature is collinear with position along the corridor**

Regressed on latitude and longitude, maximum summer temperature gives $R^2 = 0.858$, leaving a residual SD of 0.24 °C against a raw SD of 0.63 °C; it correlates with longitude at $r = -0.867$ and with great-circle distance from the Vero Beach QUI source population at $r = -0.752$. Temperature annual range is more collinear still ($R^2 = 0.903$), whereas percent impervious surface ($R^2 = 0.000$) and tree canopy ($R^2 = 0.021$) show no spatial trend. Subtracting the genome-wide background does not remove this exposure, because the background is itself environmentally structured: genome-wide QUI-like ancestry correlates with maximum summer temperature at $r = +0.688$ and genome-wide MOL-like ancestry with impervious surface at $r = +0.661$.

The two-metropolitan design is what separates the gradients. Among the Washington sites, maximum summer temperature and percent impervious surface correlate at $r = +0.821$ ($P = 0.0006$); among the Baltimore sites they are uncorrelated ($r = -0.330$, $P = 0.32$). A within-Washington analysis accordingly cannot separate them (the CpL1a temperature term falls from $P = 0.0025$ alone to $P = 0.52$ once impervious surface is added), but the full corridor can: with a metropolitan-area term and both gradients entered together, temperature retains its effect at CpL1a ($P = 0.0051$) and CpL2a ($P = 0.0050$) while impervious surface does not ($P = 0.11$ and 0.82), and impervious surface retains its effect at CpL3a ($P = 0.0100$) while temperature does not ($P = 0.93$). Fitted within each area separately, the temperature effect is present in

Washington, where maximum summer temperature spans 2.03 °C (CpL1a $P = 0.0025$, adjusted $R^2 = 0.54$; CpL2a $P = 0.0015$, adjusted $R^2 = 0.58$), and absent in Baltimore, where it spans only 0.88 °C ($P = 0.85$ and 0.44), reflecting the absence of a thermal gradient there rather than of an effect. Because maximum summer temperature is 86% predicted by coordinates corridor-wide and 88% within Washington, temperature can be separated from urbanization but not from other quantities varying along the same spatial gradient.

The stricter control of latitude and longitude as continuous covariates is reported here in full. Under that control no temperature association reaches conventional significance: the CpL1a and CpL2a ancestry associations fall to $P = 0.096$ and $P = 0.099$, on Δ and on raw ancestry alike, while the CpL3a impervious-surface association is unaffected ($P = 0.0053$), because impervious surface carries no spatial trend. The coordinate-controlled null itself is not informative, however. With maximum summer temperature 86% predicted by the coordinates, the variance inflation factor on the temperature term is 7.0 and its standard error inflates 2.65-fold, so a coefficient significant at $P = 0.005$ is expected to land near $P = 0.3$ with its point estimate unchanged, which is what we observe. Conditioning on a covariate that reproduces 86% of the predictor removes the power to test it rather than the effect. We therefore treat the coordinate control as uninformative in this design rather than as evidence against a thermal association, and neither control can establish a thermal mechanism: what the design supports is that the temperature-gradient loci respond to something tracking the thermal gradient and not to urbanization.

Metropolitan assignment is part of the analysis rather than a descriptive convenience. Four sites lie within 10 km of equal distance to the two centers, SERC (3.15 km nearer Washington), Brighton (4.02 km), Jerald (6.59 km) and Pax (6.93 km), and all fall to Washington under the rule. Reassigning all four to Baltimore, the most adverse perturbation available, changes the within-area correlations between maximum summer temperature and percent impervious surface only from +0.821 to +0.860 within Washington and from −0.330 to −0.322

within Baltimore, so the identifying contrast is a property of the two-city design rather than of how the mid-corridor sites are placed. Latitude is not the rule and does not substitute for it: the corridor runs southwest to northeast, and splitting at 39.00, 39.10 or 39.20° N gives northern-group correlations of −0.487, −0.802 and −0.882, none of which is the reported value.

**Genome-wide *molestus*-like ancestry follows impervious surface independently of position**

Impervious surface has no spatial trend along the corridor ($R^2 = 0.000$ on latitude and longitude; $r = -0.12$ with distance to central Washington), so its genome-wide association with *molestus*-like ancestry is not exposed to the positional collinearity that limits maximum summer temperature. Genome-wide MOL-like ancestry, measured outside the three candidate intervals, rose with impervious surface alone (adjusted $R^2 = 0.41$, $P < 0.001$) and kept that effect with latitude and longitude in the model ($P < 0.001$), with distance to central Washington ($P < 0.001$), and with maximum summer temperature and metropolitan area ($P < 0.001$). It correlated with impervious surface within Washington ($r = 0.90$, $P < 0.001$) and, more weakly, within Baltimore ($r = 0.55$, $P = 0.08$), and across the 20 pools outside the four within 3 km of central Washington ($r = 0.45$, $P = 0.046$). Genome-wide QUI-like ancestry showed no association with impervious surface ($P = 0.31$ alone, 0.16 to 0.59 with the same controls). PIP-like ancestry fell with impervious surface within Washington ($r = -0.76$, $P = 0.002$), as MOL-like ancestry rose there, but not detectably across the corridor ($P = 0.22$). Including the panel SNPs inside the three candidate intervals, which these genome-wide estimates exclude, barely changes per-pool ancestry (with against without them, $r = 0.98$ for QUI-like, 0.997 for PIP-like, and 0.998 for MOL-like ancestry) and changes the direction or significance of none of the main-text associations with temperature, impervious surface, or both, with or without a metropolitan-area term. The largest change is a steeper QUI-like temperature slope (+0.0055 against +0.0042 per °C, $P < 0.001$ in both), contributed by CpL1a, where QUI-like ancestry rises steeply with temperature;

MOL-like ancestry follows impervious surface as closely with the intervals as without them (adjusted $R^2 = 0.44$ against 0.41, $P < 0.001$ in both).

**Environmental predictor assignment across candidate SNPs**

Each candidate SNP was assigned to the predictor with which it showed the strongest association. Across the 1,299 candidates genome-wide, maximum summer temperature was assigned most often, at 64.4%, followed by precipitation of the warmest quarter at 9.8%, temperature annual range at 8.5%, percent impervious surface at 5.6% and tree canopy at 4.8%. The precipitation signal is almost entirely absent from the three candidate regions and falls instead among the 466 unclustered candidates lying outside them (Fig. 3). Within the three regions the composition differs sharply and is the basis for calling two of them temperature loci and one an urbanization locus. At CpL1a, 84.4% of the 654 candidate SNPs were assigned to maximum summer temperature; at CpL2a the composition is nearly the same (81.8% maximum summer temperature and 12.7% temperature annual range, or 94.5% temperature). At CpL3a the assignment falls on the urbanization predictors: tree canopy 36.2%, elevation 17.4% and percent impervious surface 15.9%. Tree canopy and percent impervious surface are near-collinear across the 24 sites ($r = -0.90$) and together account for 52.2% of CpL3a candidate SNPs, which is why percent impervious surface represents that gradient.

*CpL1a: the CYP6 cluster on chromosome 1*

**Donor-private panels give opposing ancestry gradients at CpL1a and CpL2a**

Across all 73,073 SNPs in the CpL1a interval, 123 were QUI-private, 3,762 MOL-private and one PIP-private. QUI-like ancestry rose with maximum summer temperature, from a mean of 0.088 across the six coolest sites to 0.234 across the six warmest (+0.088 per °C, adjusted $R^2 = 0.64$, $P < 0.001$), while MOL-like ancestry fell from 0.117 to 0.089 (−0.014 per °C, adjusted

$R^2$ = 0.15, P = 0.033). Across the 10,866 SNPs of the CpL2a interval the pattern reversed: 6 QUI-private SNPs gave a decline from 0.325 to 0.146 (−0.108 per °C, adjusted $R^2$ = 0.50, P < 0.001) and 30 MOL-private SNPs an increase from 0.085 to 0.174 (+0.060 per °C, adjusted $R^2$ = 0.66, P < 0.001). At CpL3a, 3,449 MOL-private SNPs increased with percent impervious surface (adjusted $R^2$ = 0.37, P = 0.001), while 4 QUI-private and 3 PIP-private SNPs were too few to analyze. Within each locus the panels share no SNPs, so their agreement rules out an artifact specific to one marker set; it is not a second independent line of evidence, because ancestry within an interval sums to a fixed total and a rise in one panel alongside a fall in another is one event seen two ways. The α values are attenuated by different amounts for each panel, summing to only about 0.63 genome-wide, so the PIP component cannot be recovered by subtraction and is not reported. PIP-private variation is nearly absent from all three intervals (1, 0 and 3 SNPs), and the panels are strongly unequal genome-wide (6,017 QUI-private, 444 PIP-private, 123,749 MOL-private), so donor attribution is resolvable on a QUI-versus-MOL axis but not a QUI-versus-PIP one.

**Reference structure at the CpL1a candidate SNPs, and why the QUI-versus-PIP contrast is unavailable**

Across the 544 candidate SNPs effectively fixed between QUI and MOL, the median binomial slope on maximum summer temperature was 1.01 on the log-odds scale, with 543 of 544 significant at P < 0.01 and none negative. Pairwise $F_{ST}$ among reference pools at these SNPs placed QUI and MOL at maximal differentiation (0.998, 95% CI 0.997–0.999) with PIP intermediate and nearer QUI ($F_{ST}$ to MOL 0.614, 0.584–0.644; to QUI 0.429, 0.396–0.461). The temperature-associated cline therefore represents movement from a MOL-like toward a QUI-like allelic state rather than away from the local PIP background. Polarizing the candidate SNPs on QUI versus PIP instead leaves almost no informative markers: 3 of the 654 candidates are near-fixed between QUI and PIP at |Δp| ≥ 0.95 and 24 at ≥ 0.70, against 544 and 579 for QUI versus

MOL. At the 544 QUI-versus-MOL near-fixed SNPs, PIP carries the QUI-type allele at a median frequency of 0.580, and the median field-pool frequency rises from 0.087 at the coolest site to 0.469 at the warmest, remaining below the PIP reference throughout. Of the 58 candidate SNPs in coding sequence at the six paralogs, 13 were nonsynonymous and 45 synonymous, with no nonsense variants; PIP carries the QUI-type allele at the 13 nonsynonymous sites at 0.56 to 0.82 (median 0.73), so these substitutions mark alleles absent from MOL and shared between QUI and PIP.

**Candidate-SNP attribution within the CpL1a interval**

The six CYP6 paralogs rank first through sixth among the 79 protein-coding genes of the interval on both gene-body and coding-sequence candidate-SNP density (Table S5). On raw counts two genes exceed them: a sodium/calcium exchanger (CQUJHB003101) with 228 gene-body candidate SNPs and a Na/K-transporting ATPase (CQUJHB014796) with 78, together 306 of the region's 654 candidates and holding the interval's two highest single-SNP $XtX_{st}$ values (330.0 and 308.7, against 296.7 for the highest CYP6 paralog). Their gene bodies span 252.6 kb of largely intronic sequence, whereas the six paralogs carry 166 gene-body candidate SNPs within 10.7 kb. Attribution is therefore reported on density rather than count.

**The CpL1a CYP6 cluster: phylogenetic placement, microsynteny and sequence similarity**

The lifted annotation identified six cytochrome P450 loci (CQUJHB009005, 008986, 009148, 017007, 001163 and 007871) forming a compact approximately 28.4 kb cluster near the center of the interval, assigned by P450Atlas as CYP6AA7-like, CYP6AA8-like, CYP6AA9-like, CYP6BZ2-like, CYP6CC2-like and CYP6BB4-like. In a 26-protein phylogeny with a curated panel of mosquito P450 references, the six focal pairs were distributed across multiple CYP6 subgroups rather than forming a single clade (Fig. S8), so the locus is a multi-gene paralog cluster rather than a tandem array of one repeated gene. The cluster is structurally conserved between *Cx. pipiens* and *Cx. quinquefasciatus*, with the six genes collinear and in the same order in both assemblies (Fig. S9).

Consensus-sequence comparisons show that sequence composition also shifts with temperature. At the nucleotide level, field pools were more similar to QUI ($\beta = 3.401$, $P < 0.001$) and PIP ($\beta = 4.966$, $P < 0.001$) than to MOL, and allowing temperature slopes to vary by reference improved fit ($\chi^2 = 16.66$, df = 2, $P < 0.001$), with steeper slopes for QUI (0.853) and PIP (0.762) than for MOL (−0.569). The amino-acid analysis recovered a similar, more asymmetric pattern (interaction $\chi^2 = 25.56$, df = 2, $P < 0.001$). Warmer populations thus accumulate more QUI-like focal genes at both levels.

**Implied copy number and per-paralog depth slopes**

Pooled across the six paralogs, the cluster depth ratio exceeds 1.0 in every field pool (1.06 to 1.64) and rises with maximum summer temperature (+0.181 in ratio per °C, adjusted $R^2 = 0.57$, $P < 0.001$), from fitted values of 1.11 at the coolest site to 1.48 at the warmest. On a diploid baseline these correspond to implied copy numbers of approximately 2.2 and 3.0: a second copy on roughly 11% of chromosomes at the coolest sites and on roughly 48% at the warmest. The cluster is thus close to unduplicated at the cool end and duplicated on about half of chromosomes at the warm end, and the expansion is modest in absolute terms, of order two to three copies rather than the high-copy amplifications described at some metabolic-resistance loci. Fitted separately, all six paralogs show positive depth–temperature slopes, each significant at $P < 0.05$: CQUJHB009148 (+0.237 per °C, adjusted $R^2 = 0.57$, $P < 0.001$), CQUJHB007871 (+0.208, 0.57, $P < 0.001$), CQUJHB001163 (+0.213, 0.47, $P < 0.001$), CQUJHB008986 (+0.149, 0.46, $P < 0.001$), CQUJHB017007 (+0.130, 0.22, $P = 0.013$) and CQUJHB009005 (+0.126, 0.28, $P = 0.0047$). Cool-site ratios range from 0.99 to 1.28 and warm-site ratios from 1.21 to 1.60, so the expansion is distributed across the cluster rather than driven by one paralog.

**Whether the CpL1a depth cline could be a mismapping artifact**

Because the reference assembly is a *Cx. pipiens* f. *pipiens* individual and the six paralogs are recently duplicated and mutually similar, two artifacts could in principle generate a depth cline: misassignment of reads among the paralogs, or recruitment of reads into the cluster from homologous sequence elsewhere. The first is excluded by conservation. Misassignment moves reads between paralogs and cannot change the cluster total, so it predicts trade-off; what we observe is co-elevation. All fifteen pairwise correlations among the six paralogs' depth ratios across the 24 pools are positive (+0.56 to +0.84, mean +0.73), and the unweighted sum of the six per-gene ratios rises from 7.06 at the six coolest sites to 8.88 at the six warmest (+1.06 per °C, $P < 0.001$). The cluster is also an outlier against the genome: fitting the depth-ratio slope on maximum summer temperature separately for each of the 13,741 genes, the six paralogs rank at the 99.72nd, 99.75th, 99.86th, 99.98th, 99.99th and 100th percentiles, the same in-genome null as the mixed model in the main text evaluated per gene.

Recruitment from outside the cluster leaves no trace we can find. Reads are finite, so recruitment requires donor sequences to lose depth. Among the 13,735 non-focal genes the slope distribution is centered (mean −0.00001, median +0.00015 in ratio per °C), and 34 genes gain at or above the smallest focal slope of +0.126 against 22 that lose at that magnitude, summing to +5.11 and −3.11 ratio units. No set of genes loses depth in the amount and pattern recruitment would require; this is suggestive rather than conclusive, because no genome-wide P450 annotation is available with which to test a named homolog set. The direction of mapping bias favors the observed signal rather than creating it: dropout of divergent QUI sequence lowers depth where QUI ancestry is high, whereas cluster depth increases with QUI-like ancestry at the locus (+10.9 ratio units per unit $\alpha$). The reference ratios settle the point more directly. Divergence from the Swedish PIP-lineage assembly runs PIP < MOL < QUI, so mismapping scaling with divergence predicts cluster depth in that order; the observed order is

MOL 0.999 < PIP 1.514 < QUI 1.579, with MOL 0.515 below PIP rather than above it. The prediction fails at the one comparison that discriminates divergence from copy number.

Cluster depth is more closely associated with QUI-like ancestry at the locus (adjusted $R^2$ = 0.70, $P < 0.001$) than with maximum summer temperature (adjusted $R^2$ = 0.56, $P < 0.001$). Because QUI-like ancestry at CpL1a itself tracks maximum summer temperature (adjusted $R^2$ = 0.64), we report partial correlations: depth against QUI-like ancestry controlling for maximum summer temperature is +0.604 ($P$ = 0.0023), and depth against maximum summer temperature controlling for ancestry is +0.241 ($P$ = 0.27). The ancestry association survives the temperature control while the temperature association does not, which is what the claim that the extra copies travel with the incoming sequence requires; the relationship with MOL-like ancestry is not detectable ($P$ = 0.11). With the two predictors correlated at $r$ = 0.81 the partition is not sharp, and we do not read the non-significant temperature partial as evidence that temperature is irrelevant.

**Read depth in the reference pools, and the haplotype that carries the extra copies**

Re-aggregating the callable-depth products with the reference pools retained gives cluster ratios of 0.999 in MOL, 1.514 in PIP and 1.579 in QUI, against a field range of 1.060 to 1.637. Collapsed paralogs in the assembly would inflate the cluster in every pool alike, and MOL instead sits at background, so the elevation is a genuine difference in copy number between lineages. The unduplicated state is MOL; both PIP and QUI carry the cluster at roughly 1.5 times background, a duplication on about half of their chromosomes. Because the high-copy state is present in PIP as well as QUI, depth alone cannot say which of the two a warming population is acquiring; the attribution to QUI rests on the QUI-private alleles at the locus, which the PIP reference lacks, and not on depth.

The references also calibrate the field cline, once depth is set against the haplotype that carries the extra copies rather than against panel ancestry, which is an index and not a proportion. At the 544 candidate SNPs nearly fixed between QUI and MOL, the mean frequency of the QUI-state allele is 0.998 in QUI, 0.582 in PIP and 0.000 in MOL, and 0.121 to 0.543 among the field pools. Field cluster depth is linear in that frequency (ratio = 1.020 + 0.979 × frequency, adjusted $R^2 = 0.80$, $P < 0.001$), and the line passes through two of the three references at their own frequencies, predicting 1.020 for MOL against 0.999 observed and 1.590 for PIP against 1.514, both within the 95% prediction intervals. At fixation it reaches 2.00, a doubling of the cluster on the QUI-state haplotype: two copies of the array on each chromosome carrying it against one on the MOL-state haplotype. The QUI reference falls below the line (1.579 against a predicted 1.997, 95% prediction interval 1.787–2.208), the direction expected if reads from the lineage most divergent from the assembly map less completely; QUI also has the lowest mapping rate of the three references (0.42, against 0.48 for PIP and MOL). With haplotype frequency in the model, neither temperature ($P = 0.72$) nor QUI-like ancestry ($P = 0.64$) improves the fit, and QUI-like ancestry at the locus tracks haplotype frequency closely ($r = 0.96$). The depth cline is therefore the haplotype-frequency cline seen through copy number: the extra copies travel with the QUI-state haplotype, and the coolest field pools sit near the MOL state because they carry that haplotype at only 12–22%. Depth places the duplication on a haplotype that QUI and PIP share and MOL lacks; it cannot by itself say from which lineage the field pools acquired it.

*CpL2a: the Yan-centered interval on chromosome 2*

**Gene content of the CpL2a haplotype block**

BLASTP against SwissProt confirmed the gene at the center of the signal, CQUJHB014704, as the ortholog of *Yan* (ets DNA-binding protein pokkuri), and identified within the surrounding approximately 2.5 Mb haplotype block CQUJHB002678 as *takeout* and three ABC-family transporters at approximately 14.3 Mb as multidrug resistance-associated protein 1

(ABCC1/MRP1) paralogs (CQUJHB009840, CQUJHB007929) and a canalicular multispecific organic anion transporter (CQUJHB002428). Genes within the 500 kb core carrying candidate SNPs are listed in Table S7.

**Stability of the six-SNP QUI-private panel at CpL2a**

The QUI-private panel within the CpL2a interval contains six SNPs, three of them at consecutive positions (13,688,362–13,688,364), so it carries fewer independent signals than its size suggests. Fitted individually, four of the six decline with temperature (the three consecutive positions at −0.168, −0.165 and −0.167 per °C, all $P < 0.001$; and 13,995,002 at −0.145, $P = 0.0022$), while the remaining two sit near zero frequency and are flat (13,762,068, $P = 0.88$; 13,798,651, $P = 0.79$). The signal therefore rests on two independent positions. Three checks bound what that supports. Dropping each SNP in turn leaves the panel-mean slope between −0.096 and −0.130, all with $P < 0.001$. A bootstrap over the six SNPs (1,000 resamples) gives a 95% interval of −0.163 to −0.049 with 99.8% of resamples negative. Most informatively, relaxing the panel definition grows the panel without moving the estimate: at ≥ 0.90 / ≤ 0.10 it holds 53 SNPs and the Δ slope is −0.117 per °C (adjusted $R^2 = 0.65$, $P < 0.001$), and at ≥ 0.85 / ≤ 0.15 it holds 217 SNPs with a slope of −0.122 (adjusted $R^2 = 0.63$, $P < 0.001$), against −0.112 at the strict threshold; a thirty-six-fold increase in panel size changes the slope by under 10%. Two estimators appear in these analyses: the panel means in the main text and the leave-one-out and bootstrap analyses are slopes on raw α, whereas the threshold analysis is on Δ, which subtracts each pool's genome-wide panel background. At the strict threshold they give −0.108 and −0.112 per °C, a difference immaterial to any conclusion.

**PIP–MOL sequence divergence at CpL2a beyond SNPs**

The candidate SNPs do not separate PIP from MOL, but the interval does. Across all 10,866 SNPs called in the 500 kb interval, per-SNP Reynolds $F_{ST}$ between PIP and MOL averaged 0.78, above the chromosome 2 average of 0.52, and the interval contained 39 opposite-fixed SNPs (PIP ≥ 0.95 and MOL ≤ 0.05 or vice versa), none of them among the 110 candidates. Mean |Δp| between PIP and MOL was 0.27 across the interval, against 0.21 at the candidate SNPs, 0.19 across chromosome 2, and 0.23 genome-wide. Haploid genotype calling on the reference BAMs across chr2:13.5–14.0 Mb identified 1,118 indels, of which 558 (49.9%) were fixed-different between PIP and MOL; sizes ranged from 1 to 41 bp, 78% in non-homopolymer context, and at sites where QUI carried a genotype it matched PIP at 67% of fixed-different indels (323 of 479), consistent with the non-QUI framing used for this locus. Per-base depth profiling revealed asymmetric mapping failures: within the interval PIP had zero depth at 58,541 bp (11.7%) in runs > 100 bp, against 95,614 bp (19.1%) for MOL and 133,984 bp (26.8%) for QUI, and 54,792 bp of the MOL zero-depth bases fell where PIP had nonzero depth against only 17,719 bp the reverse, a 3:1 asymmetry indicating that MOL carries structural differences at CpL2a that prevent reads from mapping. These represent sequence the SNP-calling pipeline cannot interrogate.

*CpL3a: the molestus inversion on chromosome 3*

**Inversion breakpoint coordinates**

The approximately 11.6 Mb extent of the CpL3a block, its uniform across-locus differentiation profile, and the independent identification of an inversion in this region of *Cx. pipiens* (inversion 3q; Cebrián-Camisón et al. 2026) together implicate a segregating inversion. DELLY's highest-confidence inversion calls were in MOL (paired-end support 6–10 across three concordant calls), with lower-confidence support in PIP (a single call at comparable coordinates) and none in QUI within the candidate window. Cebrián-Camisón et al. (2026) characterized inversion 3q at ~149–161 Mb on the same idCulPipi1 assembly through windowed PCA and linkage disequilibrium analysis, and found that all *molestus* specimens carried the inverted homokaryotype. Their approximate breakpoints agree with our DELLY calls (~149.4 and ~161.0 Mb) to within ~1 Mb at both ends.

**A genome-wide windowed PCA recovers the inversion without prior coordinates**

Scanned without reference to the candidate intervals, the genome contained one extended region with a distinct local structure, and it was CpL3a (Fig. S10). Ranked by PC1 variance alone, the 25 most structured of the 1,066 windows all lie within 2 Mb of a candidate locus (eight at or beside CpL1a, two at CpL2a, and fifteen at CpL3a; main text Fig. 3B), and the first window elsewhere ranks 26th (chromosome 1, 118.75 Mb); in 1 Mb windows the 18 most structured of 534 lie within 2 Mb of a candidate locus. Across the 23 windows of 500 kb from 149.5 to 161.0 Mb, PC1 explained 22–32% of among-pool variance (median 28.4%, against a genome-wide median of 10.8%). On the second local-PCA axis these windows took values of 0.56–0.70, as did the window spanning the proximal breakpoint (149.0–149.5 Mb). Values fell through intermediate levels (0.26–0.48) within about 1.5 Mb on either side, and no window elsewhere in the genome exceeded 0.31. The block therefore matches the DELLY breakpoints (~149.4 and ~161.0 Mb) to within one window, and 1 Mb windows gave the same boundaries. Other windows exceeded the 95th percentile of PC1 variance, including those at CpL1a and CpL2a, but none shared the local structure of CpL3a. Within the block, each pool held a stable position along PC1 from window to window, and pools with more impervious surface sat further toward the *molestus*-like end (median r = 0.64 between PC1 score and percent impervious surface across the block). Outside it, the ordering of pools changed between adjacent windows. A stable, environmentally ordered frequency for each population across the whole block is the form an inversion takes in pooled data, in place of the discrete karyotype clusters that individual genotypes give.

**The CpL3a inversion shows no localized gene-level candidate-SNP concentration**

Beyond the summary in the main text, gene-body candidate-SNP densities were uniformly low across the 11.6 Mb span, and the genes that do carry coding candidate SNPs (Table S8) are spread across functions with plausible relevance to an urban-associated haplotype: two facilitated glucose transporters, an ABCA3-class transporter, a 60E-type voltage-gated sodium channel, and a gustatory and an odorant receptor. None is nominated as the target of selection.

**The CpL3a inversion is not enriched for chemosensory receptor genes**

The inversion span contains five curated chemosensory genes, one ionotropic receptor (*Ir174*), one gustatory receptor (*Gr74*) and three odorant receptors (*Or42*, *Or43*, *Or132*), in four tandem arrays. Neither count is exceptional against the width-matched null: five genes against a null median of six (P = 0.634) and four arrays against a median of four (P = 0.502), and no family is enriched (ionotropic receptors 1 against a median of 2, P = 0.756; odorant receptors 3 against 1, P = 0.288; gustatory receptors 1 against 0, P = 0.388; odorant-binding proteins 0 against 1, P = 1.000). The curated annotation did not materially change the interval's content: the lifted JHB2020 annotation recovers four chemosensory genes and the curated annotation five, and the two genes named in the main text are recovered by both under different nomenclatures (GR2a, CQUJHB002662, is *Gr74*; Or67d, CQUJHB008968, is *Or43*). None of the five appears in the host-preference windows of Bell et al. (2024) or in the differentially expressed or sequence-divergent sets of Menna et al. (2026), so none carries independent functional evidence of a role in host preference.

The sparse candidate-SNP signal within the inversion is nevertheless not placed independently of these genes: two of the five carry a candidate SNP within ± 2 kb (*Gr74*, maximum $XtX_{st}$ 81.8; *Or43*, 76.8), whereas randomizing the positions of the 69 candidate SNPs across the inversion yields a median of zero genes hit (uniform null P = 0.023; circular-shift null

P = 0.021). This is a weak result resting on two genes, and the top predictor at these two SNPs is maximum summer temperature at *Gr74* and precipitation of the warmest quarter at *Or43*, neither the urbanization gradient along which the block is structured. Chromosome 3 does carry a substantial chemosensory array nearby, thirty-two genes (twenty-seven ionotropic receptors) spanning 161.22–163.95 Mb, but it begins 203 kb distal to the 161.02 Mb breakpoint. Loci close to inversion breakpoints can remain in linkage disequilibrium with the inversion, but in these data the allele-frequency correlation between candidate SNPs and flanking SNPs falls to the chromosomal baseline within the first distance bin outside the breakpoints (main text Fig. 4), so across the 24 pools the array does not covary with the inversion haplotype.

**Robustness of the CpL3a MOL-like signal to MOL fixation**

Because the MOL-private panel is the largest of the three and carries the CpL3a result, we asked whether “MOL-private” is partly an artifact of the MOL reference being fixed for alleles that are simply common across the corridor, by recomputing the corridor-wide median Δ at CpL3a while progressively excluding panel SNPs with high mean field frequency. The median Δ is +0.080 using all 3,449 panel SNPs and does not weaken as the exclusion tightens: +0.080 excluding SNPs with a corridor mean above 0.80, +0.086 above 0.60, +0.091 above 0.50 and +0.084 above 0.40, the last still retaining 3,085 SNPs. Excluding more aggressively reduces the estimate (+0.062 at 0.30, +0.038 at 0.20), but those thresholds cut below the locus median α of 0.233 and so remove the SNPs carrying the signal being measured rather than testing it.

**The CpL3a urban association against genome-wide *molestus*-like ancestry**

Regressed on the same pool’s genome-wide MOL-like ancestry, the inversion’s MOL-like ancestry rose with a slope of 2.36 (SE 0.46; adjusted $R^2$ = 0.52), against the slope of one that Δ assumes. With genome-wide ancestry and impervious surface in one model, genome-wide ancestry predicted the inversion’s ancestry (P = 0.006) and impervious surface did not (P =

0.21). On the log scale the slope on genome-wide ancestry is 1.00 and impervious surface again adds no significant effect (P = 0.13), so the inversion's MOL-like ancestry is close to a constant multiple of the genome-wide level. The same model at the temperature loci gives the reverse: maximum summer temperature predicts CpL1a ancestry beyond the background (P = 0.0014; P = 0.0023 on the log scale) and the background does not (P = 0.10), and at CpL2a the corresponding values are P = 0.0016 (0.0022) and P = 0.98.

*Cross-locus comparisons*

**Haplotype-block diagnostics at the three candidate loci**

All three loci behave as coherent allele-frequency units against the genome-wide null of 10,000 width-matched regions. At CpL1a (108.5–111.0 Mb; 68,818 region SNPs), PC1 explained 41.2% of among-pool variance (permutation P < 0.001), pool scores on PC1 were near-perfectly correlated with each pool's mean $F_{ST}$ to QUI (|r| = 0.995; P < 0.001), and the across-pool variance in that $F_{ST}$ was elevated (0.041; P < 0.001): the region varies across pools along a single QUI-ancestry axis. At CpL3a (149.4–161.0 Mb; 234,538 SNPs) all three diagnostics were likewise supported, with PC1 at 28.1% (P < 0.001), |r| = 0.995 to MOL (P < 0.001) and an elevated $F_{ST}$ variance (0.010; P = 0.026), so the approximately 11.6 Mb interval of elevated $XtX_{st}$ reflects allele-frequency variation in one block of MOL-like ancestry rather than independent selection at the loci it contains. At CpL2a (12.5–15.0 Mb; 56,287 SNPs), read on the QUI-versus-non-QUI axis, the first two diagnostics were strongly supported (PC1 25.6%, P < 0.001; |r| = 0.990 to QUI, P = 0.004) while the across-pool $F_{ST}$ variance was elevated but not significant (0.0068; P = 0.068): the region segregates as a coherent unit whose primary axis indexes QUI-like versus non-QUI-like ancestry, with more modest donor-ancestry differentiation across pools than at CpL1a.

**Linkage architecture at the three candidate loci**

At CpL1a the candidate SNPs were strongly intercorrelated (median $r^2 = 0.529$), well above the chromosomal baseline (approximately 0.02), declining only modestly across the 2.5 Mb cluster (from approximately 0.73 among the closest pairs to approximately 0.39 among the most distant); pairs with SNPs outside the cluster decayed from approximately 0.09–0.11 just outside it to baseline by 5–8 Mb, while linkage across all SNPs in the interval sat only just above baseline (median 0.053), so the region does not behave as a single recombinational block. At CpL2a the candidate SNPs were correlated at median $r^2 = 0.444$, and the association extended well beyond them: interval-to-flank correlation decayed smoothly from approximately 0.28 immediately outside the core, through approximately 0.18 at 20–100 kb, 0.10 at 250–500 kb, 0.06 at 750 kb–1 Mb and 0.04 at 1–1.5 Mb, reaching the baseline (approximately 0.02) only at 2–3 Mb, which places a recombinationally coherent haplotype of approximately 1–1.5 Mb on each side of the 500 kb core. At CpL3a the candidate SNPs showed uniformly high correlation (median $r^2$ approximately 0.30) that did not decay across the entire 11.6 Mb interval, while flanking pairs collapsed to baseline (approximately 0.03) immediately outside the candidate boundaries. The contrast between CpL1a and CpL2a is therefore one of amplitude beside the selected core rather than of reach: CpL1a's flank correlation extends further while being weaker throughout. Comparisons are made against the same-chromosome baseline rather than against the cross-locus control (set iv), because that control is composed of the candidate SNPs of the other two loci and is inflated by shared environmental ascertainment (median $r^2 = 0.261$ at CpL1a against a baseline of 0.023).

**Locus statistics against a differentiation-matched null**

Against the differentiation-matched null, the Δ–environment correlation at each focal locus exceeds that of every retained comparison region: +0.804 against a maximum of +0.502 among 291 regions at CpL1a, +0.722 against +0.658 among 239 at CpL2a, and +0.528 against

+0.504 among 541 at CpL3a. Because the null is not centered on zero this is reported as a rank. The among-pool variance in donor-likeness is unaffected by that centering problem: at CpL3a it is 4.2 times the median of matched regions ($P < 0.001$) and at CpL1a 9.3 times ($P < 0.001$); at CpL2a it is 2.7 times and not significant ($P = 0.20$), and no claim is made for it. The focal intervals are also enriched for donor-private markers relative to their SNP content: CpL1a holds 2.04% of all QUI-private SNPs within 0.63% of all SNPs (3.3-fold), and CpL3a 2.79% of MOL-private SNPs within 2.11% of SNPs (1.3-fold); CpL2a is depleted for MOL-private markers by a comparable factor, consistent with its signal being a QUI deficit rather than a MOL excess.

Matching comparison regions on panel size as well shows that the small-panel shift must be measured per panel and locus. At CpL2a on the MOL-private panel (30 SNPs; focal correlation +0.722), restricting to the 30 matched regions whose panel size lies within a factor of two of the focal panel gives a null shifted positive (mean $r = +0.221$), and none of the 30 reaches the focal value. On the six-SNP QUI-private panel (focal correlation −0.731; none of 456 matched regions reaches it in absolute value), the 283 regions with comparably small panels give a null shifted positive (mean $r = +0.137$) and none reaches the focal value ($P < 0.01$), so at this panel the small-panel bias runs against the reported direction rather than producing it. At CpL1a only 9 of 291 matched regions carry a panel within a factor of two of the focal 123, because the interval is itself enriched for QUI-private markers, and we report that enrichment rather than a null. At CpL3a the null remains negatively shifted after panel-size matching (mean $r = -0.275$ across 317 regions), so the rank statement is the defensible summary there.

**Statistics supporting the main-text Results**

These values support analyses reported in the main-text Results. Genome-wide, with temperature and percent impervious surface both in the model, QUI-like ancestry retained its temperature effect ($P < 0.001$) and showed no effect of impervious surface ($P = 0.83$); with a metropolitan-area term in the model, the temperature trends in QUI-like and MOL-like ancestry persisted ($P = 0.016$ and 0.002), whereas the PIP-like decline did not ($P = 0.17$). At CpL1a, pairs joining a candidate SNP to a SNP outside the interval stayed weakly correlated at every distance (median $r^2$ 0.04–0.11) and were indistinguishable from the baseline by 5–8 Mb. The per-population Δ tracked summer temperature at CpL1a (adjusted $R^2 = 0.62$; one-sided $P < 0.001$; permutation $P < 0.001$ against width-matched randomly selected regions); in the two-gradient model, temperature retained its effect ($\beta = 0.078$, $P < 0.001$) while impervious surface contributed nothing ($\beta = 4.0 \times 10^{-4}$, $P = 0.16$) and did not improve the model ($F = 2.16$, $P = 0.16$). At CpL2a, mean |Δp| between PIP and MOL across the candidate SNPs is 0.21, below the interval-wide value of 0.27; Δ tracked temperature (adjusted $R^2 = 0.55$; one-sided $P < 0.001$; permutation $P < 0.001$), and in the joint model temperature retained its effect ($\beta = +0.057$, $P < 0.001$) while impervious surface contributed nothing ($P = 0.84$). At CpL3a, the median Δ was 0.081 ($P < 0.001$); Δ tracked percent impervious surface (adjusted $R^2 = 0.245$; one-sided $P = 0.004$; permutation $P < 0.001$), and in the two-gradient model impervious surface retained its effect ($\beta = 6.4 \times 10^{-4}$, $P = 0.021$) while temperature contributed nothing ($\beta = 0.017$, $P = 0.20$) and did not improve the model ($F = 1.75$, $P = 0.20$). When the genome-wide ancestry level and the relevant predictor were entered together, temperature predicted locus ancestry beyond the genome-wide level at CpL1a and CpL2a ($P = 0.001$ and 0.002) and the genome-wide level added nothing ($P = 0.10$ and 0.98), whereas at CpL3a genome-wide *molestus*-like ancestry predicted the inversion's ancestry ($P = 0.006$) and impervious surface added no significant effect ($P = 0.21$). At CpL1a, averaged over the 24 populations Δ was small and not significantly positive, as expected for a locus elevated only at the warmest sites, and adding a metropolitan-area term left the result unchanged (temperature $P = 0.005$, impervious surface $P = 0.11$). The CYP6 cluster read-depth ratio was roughly 31% higher in the warmest populations than in the coolest. At CpL2a, interval-to-flank pairs began at median $r^2 \approx 0.28$, roughly three times the corresponding CpL1a value, and decayed to baseline by 2–3 Mb, against 5–8 Mb at CpL1a.

*Insecticide resistance outside the candidate set*

**A characterized resistance allele outside the candidate set**

A leucine-to-phenylalanine substitution at chromosome 2 position 191,086,281 in the voltage-gated sodium channel gene *para* (CQUJHB005371) falls at residue 1035 of the 2,147-residue canonical transcript, which aligns to residue 1014 of *Musca domestica* Vssc1 at 80% amino-acid identity; both residues are leucine in an identical 17-residue context (LATVVIGNLVVLNLFLA) that occurs once in each protein. The reference codon is TTA and the variant TTT (c.3105A>T on transcript CQUJHB005371.R8259), and no other variant segregates in that codon. The corridor therefore carries *kdr* L1014F under the standard numbering, and not the serine or histidine substitutions at the same residue reported in other populations of the *Cx. pipiens* assemblage. Across the 24 field pools the resistant allele averaged 0.344 (range 0.080–0.821; Fig. S7; Table S3), unassociated with maximum summer temperature ($P = 0.32$) or percent impervious surface ($P = 0.16$); the highest frequency fell at the rural SERC site (0.821, 0.5% impervious surface). This position could not have been recovered by the genome scan: its $XtX_{st}$ of 128.2 exceeds the major-outlier threshold of 106.6, but no covariate returned Bayes-factor support (best BF = −1.9 dB), so it falls in the $XtX_{st}$-only class the dual-outlier criterion excludes. That class is close to null expectation genome-wide (13,270 observed major outliers against 11,630 expected under the pseudo-observed null, a ratio of 1.14), so it is not a reservoir of undetected loci, but functionally characterized sites can sit within it. Among the references the resistant allele was near-fixed in MOL (0.912) and effectively absent from PIP (0.000) and QUI (0.026); with one colony per lineage we do not assign *kdr* a donor on this evidence.

Two correlations that might be expected to follow do not survive testing. Across the field pools *kdr* frequency correlates with genome-wide PIP-like ancestry (Spearman $\rho = -0.52$), but against SNPs matched to *kdr* on differentiation and allele frequency that correlation is weaker than the median matched SNP achieves (null median $|\rho| = 0.574$; empirical $P = 0.72$; MOL and

QUI panels P = 0.88 and 0.97). The reason is general: $XtX_{st}$ measures the magnitude of a SNP's deviation from the expected population covariance, not its direction, so strongly differentiated SNPs track the dominant ancestry axis as a matter of course (a random SNP correlates with PIP-like ancestry at $|\rho| = 0.165$, indistinguishable from the chance expectation of 0.163, rising to 0.323 in the top $XtX_{st}$ decile and 0.574 among SNPs as differentiated as *kdr*). We therefore make no ancestry claim for *kdr*. Its covariation with the CpL1a CYP6 cluster depends on the direction the question is asked. Against 13,553 contiguous six-gene runs elsewhere in the genome, the cluster is unusual in how strongly its depth tracks *kdr*, raw ($\rho$ = +0.47, P = 0.022) and after removing maximum summer temperature and genome-wide PIP background (partial r = +0.56, P = 0.006), unchanged when the null is restricted to the most variable one per cent of runs (P = 0.007). The symmetric test does not agree: against matched SNPs, *kdr* is unremarkable in how it tracks cluster depth (raw P = 0.89; partial P = 0.053), and an equally differentiated random 2.5 Mb window tracks *kdr* as well as CpL1a does (P = 0.19). We report the covariation as suggestive of target-site and metabolic resistance accumulating together under a pressure on neither environmental gradient, and not as evidence bearing on the interpretation of CpL1a. No other canonical target-site substitution was found; in particular *Ace-1* G119S was absent, the three glycine–serine variants in the acetylcholinesterase genes lying outside the G119 context. Across all 63 genes we recovered 2,039 nonsynonymous variants (Table S3), mostly in carboxylesterases of the Ester locus (1,263 across 26 genes) and glutathione S-transferases (237 across 20); no environmental association among them survives correction for multiple testing, and they are reported as a resource rather than as candidates.

**Supplementary Figures**

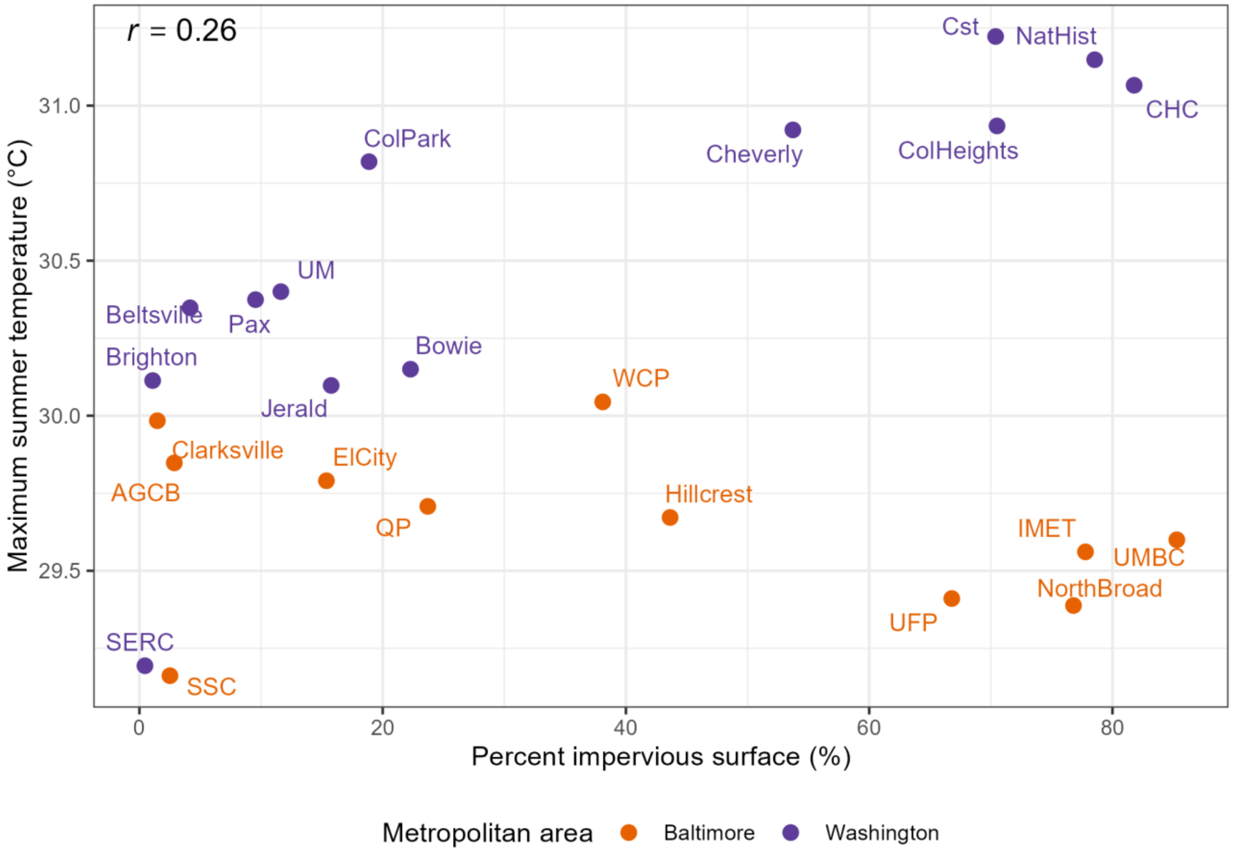


**Figure S1. The two environmental gradients are largely independent across the 24 field sites.** Maximum summer temperature (°C) plotted against percent impervious surface (%) for each of the 24 collection sites, colored by metropolitan area. The Pearson correlation is weak ($r$ = 0.26), because temperature varies chiefly between the cooler Baltimore and the warmer Washington, DC metropolitan areas while impervious surface varies within each of them. This near-independence allows allele-frequency signals tracking the temperature gradient to be distinguished from those tracking the urbanization gradient.

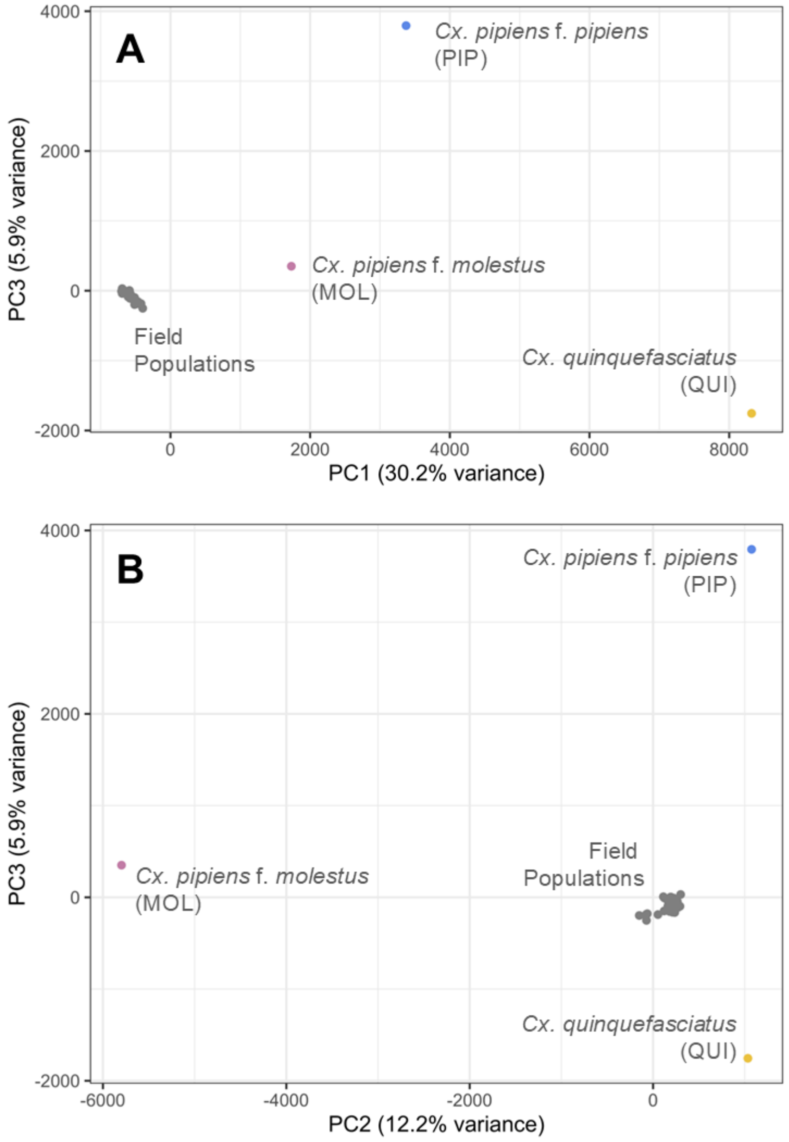


**Figure S2. Additional principal component axes.** (A) PC1 versus PC3 and (B) PC2 versus PC3 for all 27 pools (24 field and 3 reference). Colors follow Figure 2: PIP (blue), QUI (yellow), MOL (pink), field pools (grey). PC3 explains 5.9% of the variance and separates PIP from QUI along an axis orthogonal to their separation on PC1 and PC2. The 24 field pools remain clustered near the origin in both projections, consistent with the tight grouping in the PC1-versus-PC2 space (Fig. 2).

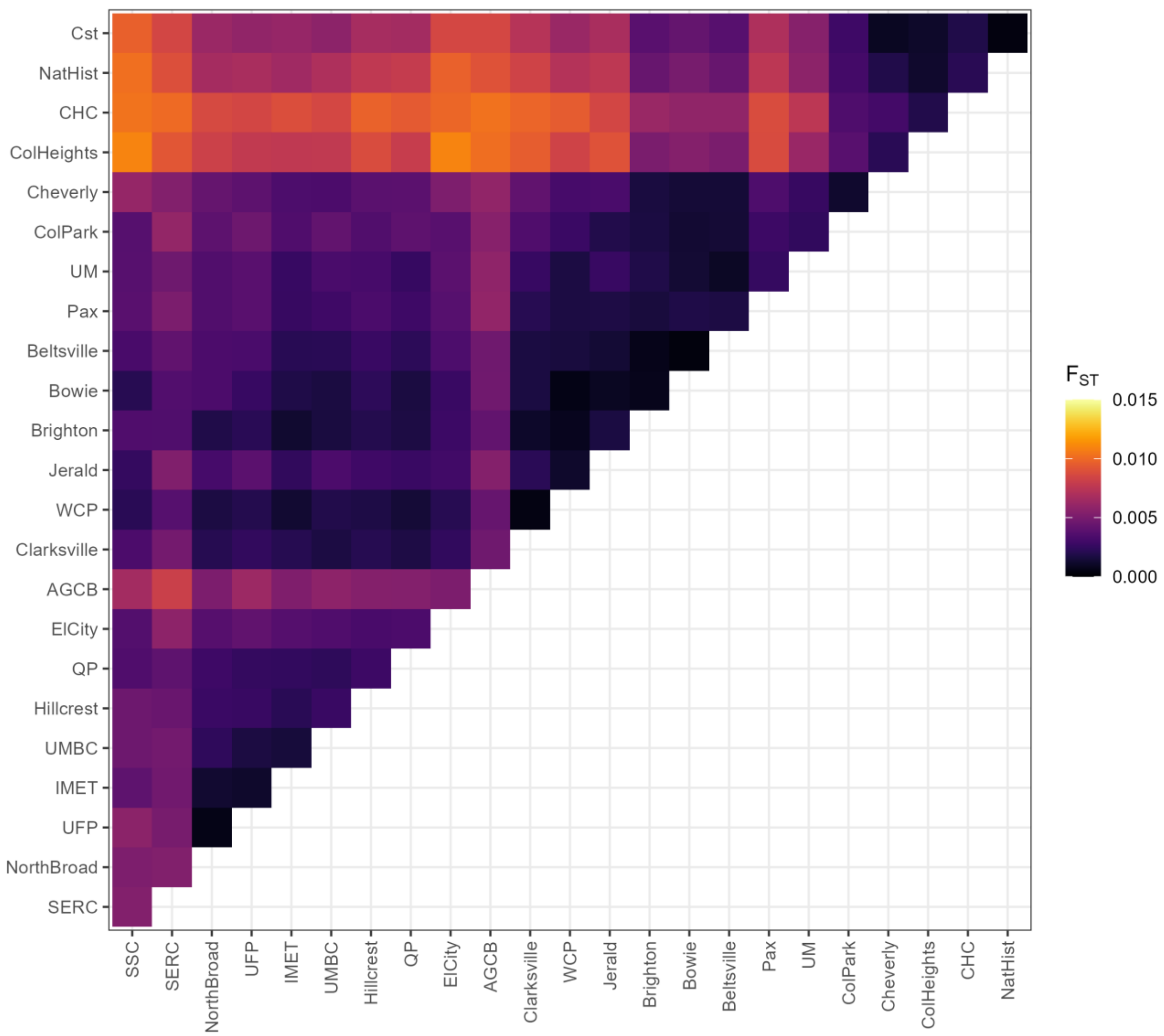


**Figure S3. Genome-wide differentiation among the 24 field pools is low.** Pairwise $F_{ST}$ between all 24 field-collected *Culex pipiens* pools, estimated genome-wide in poolfstat using the analysis-of-variance estimator, with each pool treated as 80 chromosomes (40 diploid individuals per pool). Pools are arranged by maximum summer temperature. Across all 276 pairwise comparisons, $F_{ST}$ was low (mean 0.004, range 0.0003–0.0110), indicating that the Baltimore–Washington field pools share a largely undifferentiated regional genomic background.

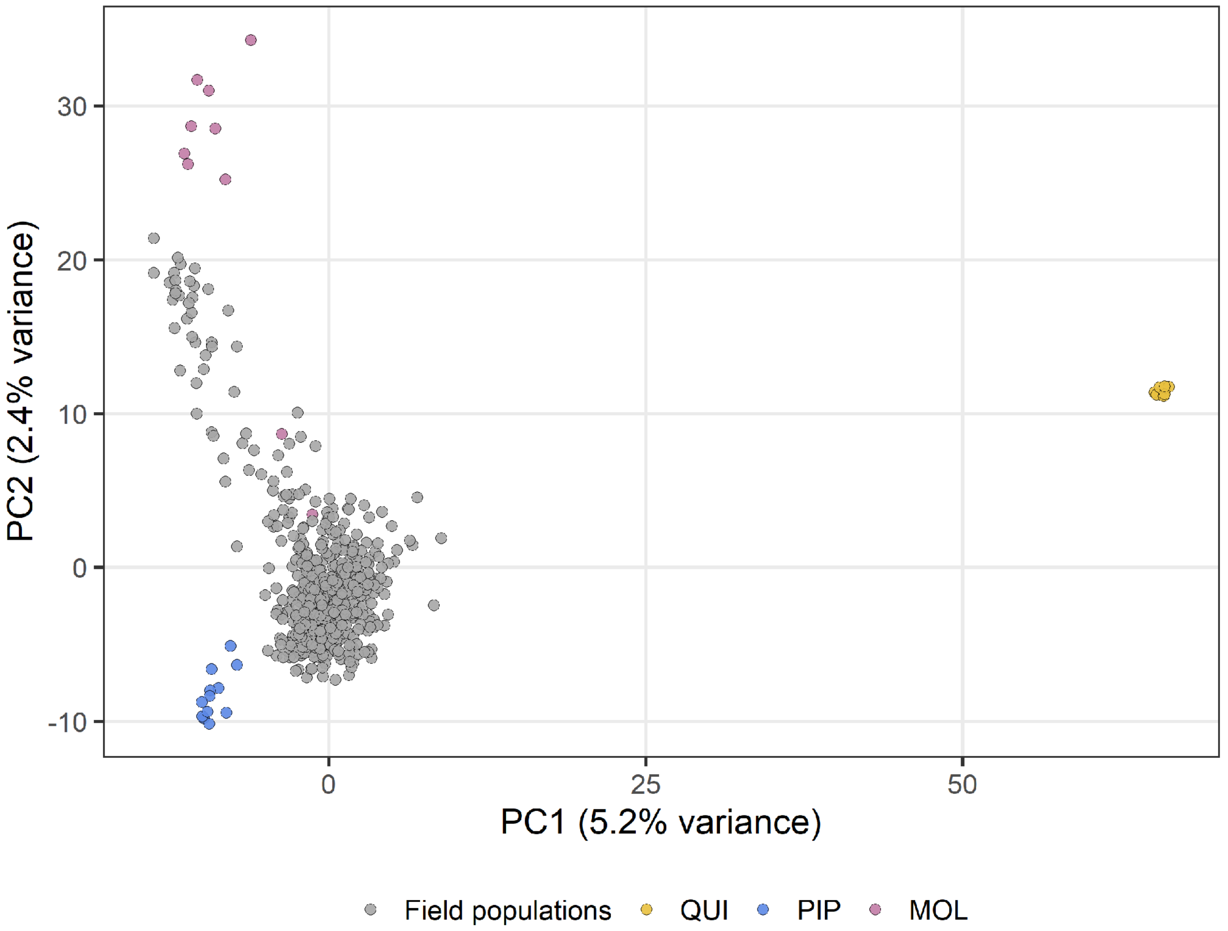


**Figure S4. Individual-level ddRADseq ordination of corridor mosquitoes and the three reference lineages.** Principal component analysis of chain-averaged posterior mean genotypes (computed from the entropy posterior genotype probabilities) at 7,258 SNPs for 553 individuals: 521 wild-caught mosquitoes from 15 sites across the Maryland and Washington, DC portion of the corridor (grey), together with individuals drawn from the same source populations as the three reference pools, QUI (yellow), PIP (blue) and MOL (pink). PC1 explains 5.2% of the variance and PC2 2.4%. Field individuals form a single continuous cloud spanning the PIP and MOL references, with no discrete subgroups and no individual intermediate between that cloud and the well-separated QUI reference.

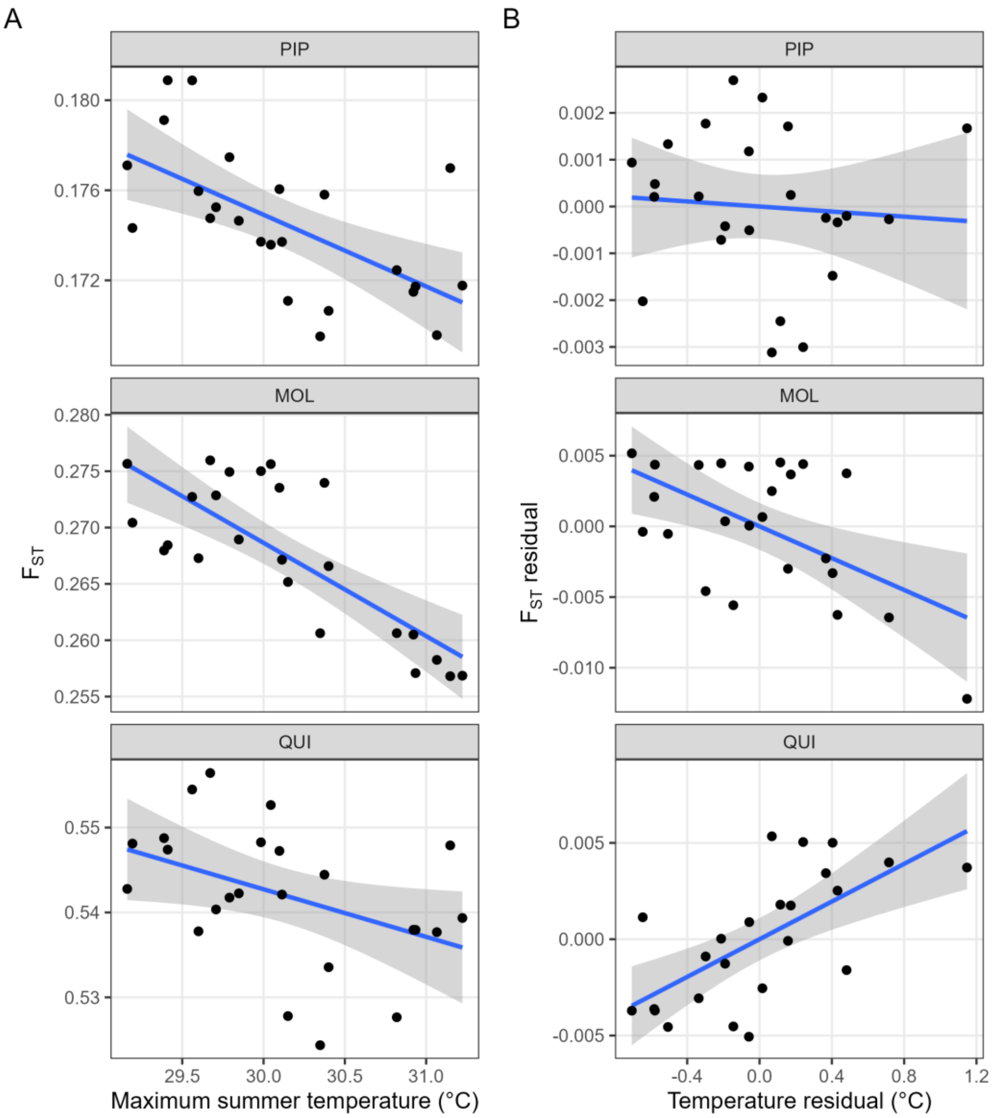


**Figure S5. Genome-wide differentiation of the field pools from each reference declines with summer temperature, but part of the pattern is a diversity artifact.** (A) Genome-wide $F_{ST}$ between each of the 24 field pools and each reference pool, PIP, MOL and QUI (separate panels), estimated in poolfstat with the analysis-of-variance estimator, plotted against site maximum summer temperature (°C). Each point is one field pool; blue lines are ordinary-least-squares fits with 95% confidence bands. Note that the $F_{ST}$ scale differs among panels: the field pools are least differentiated from PIP ($F_{ST} \approx 0.17$), intermediate from MOL ($\approx 0.27$), and most differentiated from QUI ($\approx 0.54$). In separate univariate linear models, $F_{ST}$ declined significantly with maximum summer temperature against all three references (vs. PIP, adjusted $R^2 = 0.36$, $P = 0.001$; vs. MOL, adjusted $R^2 = 0.59$, $P < 0.001$; vs. QUI, adjusted $R^2 = 0.15$, $P = 0.04$), with the relationship strongest and tightest against MOL and weakest against QUI. (B) Partial-residual (added-variable) plots after controlling for genome-wide expected heterozygosity. Once the diversity gradient is removed, the decline against PIP disappears ($P = 0.72$), confirming it as a diversity artifact; the decline against MOL persists ($P = 0.006$); and the decline against QUI reverses to a significant increase ($P < 0.001$), indicating that warmer, more diverse populations are slightly more differentiated from QUI after accounting for their higher heterozygosity.

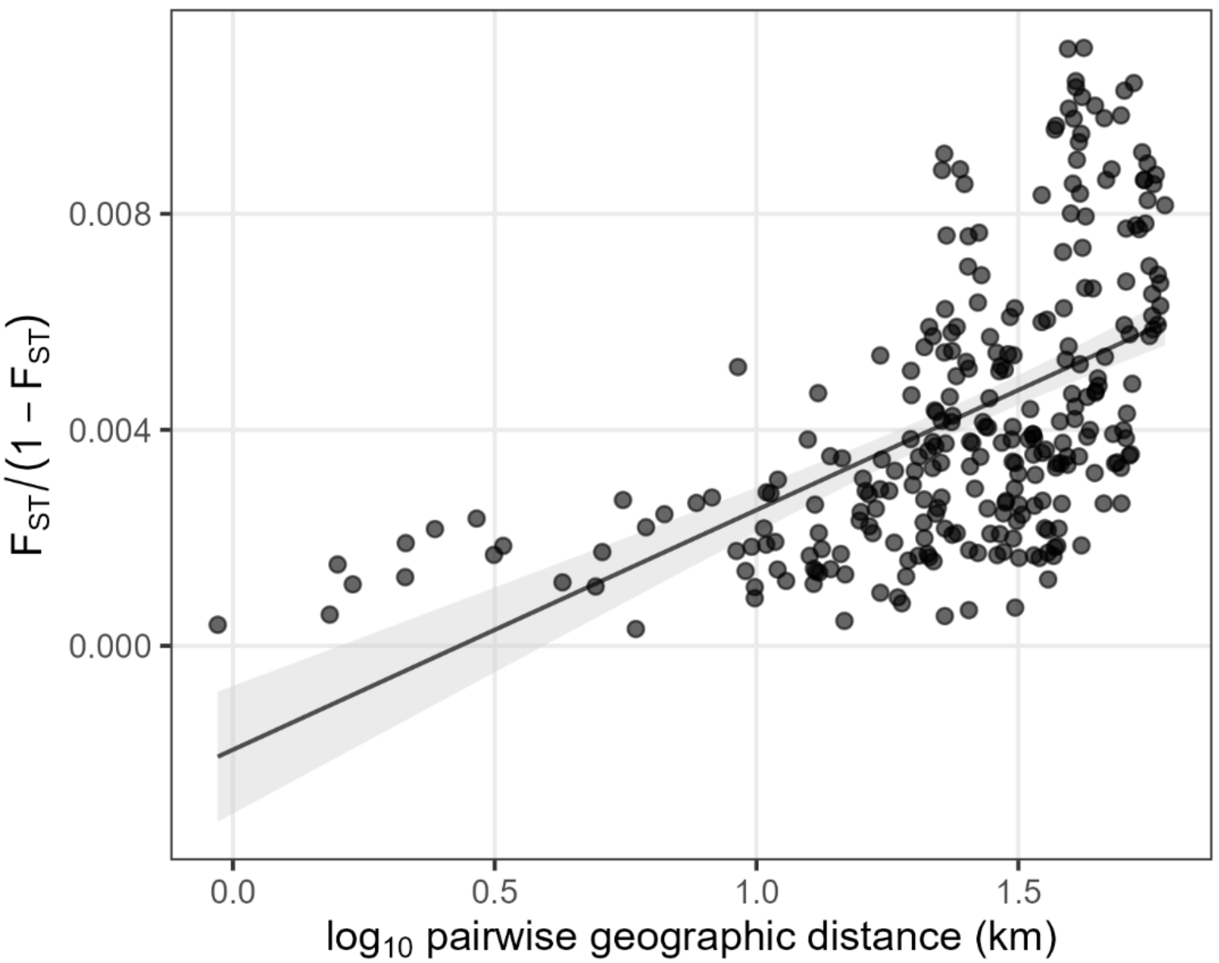


**Figure S6. Genome-wide differentiation among field pools increases with geographic distance (isolation by distance).** Linearized $F_{ST} /(1 - F_{ST})$ (Rousset 1997) versus $\log_{10}$ pairwise geodesic distance (km) for all 276 pairs of the 24 field pools, over a distance range of 0.9–60 km; pairwise $F_{ST}$ values are the genome-wide poolfstat estimates. Each point is one pool pair (semi-transparent to show overplotting). The line is the ordinary-least-squares fit and the shaded band its 95% confidence interval. The relationship is significantly positive (slope = $4.4 \times 10^{-3}$, SE = $4.2 \times 10^{-4}$; Pearson $r = 0.538$, one-sided $P < 0.001$; adjusted $R^2 = 0.29$) and was confirmed by a Mantel permutation test ($r = 0.538$, $P < 0.001$, 9,999 permutations). Geographic distance accounts for roughly a third of the among-pair variation in differentiation, consistent with active but distance-limited gene flow across the corridor.

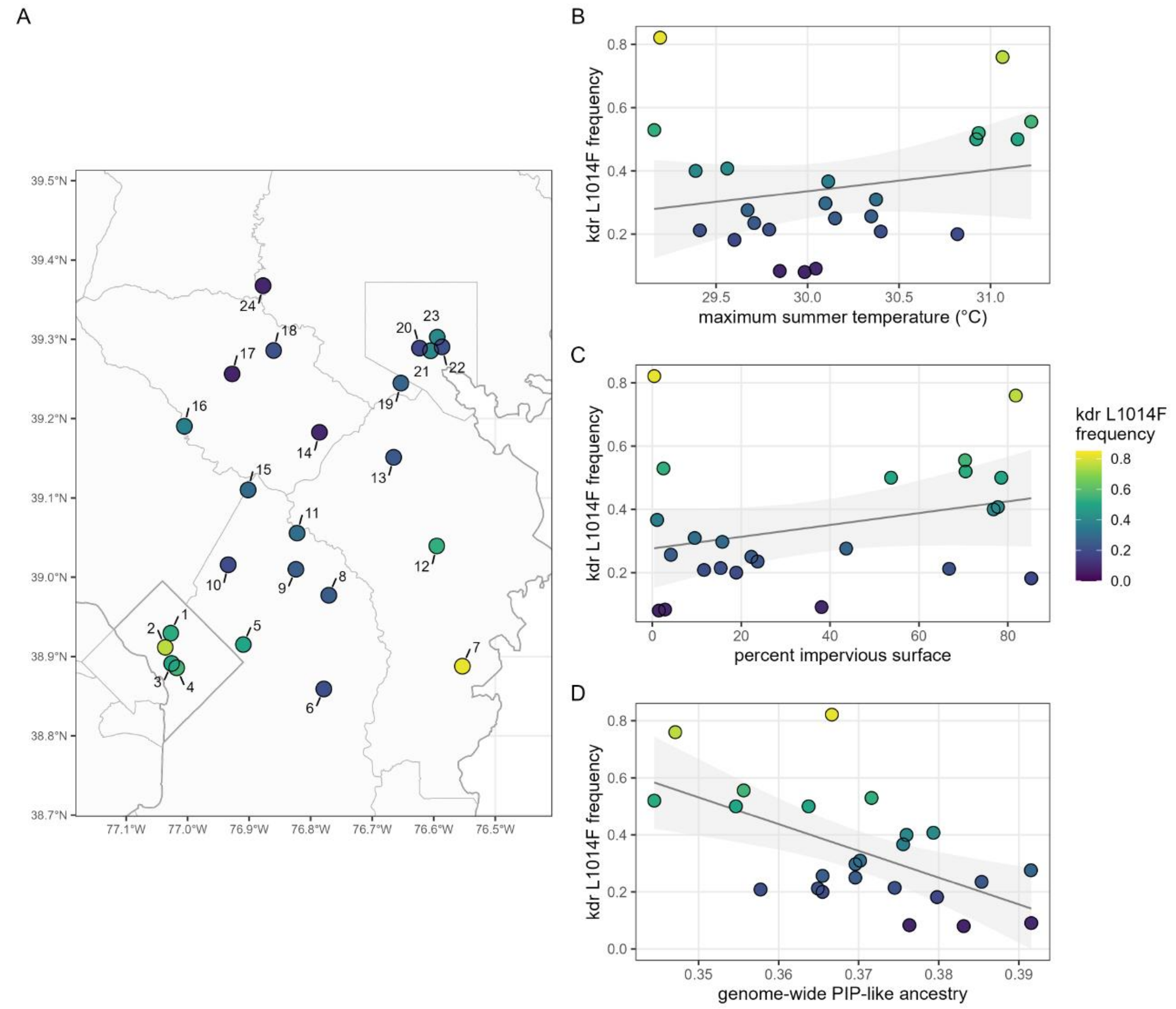


**Figure S7. *kdr* (*Vgsc* L1014F) frequency across the 24 field pools.** (A) Per-pool frequency of the resistant (phenylalanine) allele at chromosome 2 position 191,086,281, plotted at each collection site; point fill gives allele frequency and site numbers follow Figure 1. (B) Frequency against maximum summer temperature and (C) against percent impervious surface, with least-squares fits shown (neither significant). (D) Frequency against genome-wide PIP-like ancestry; this correlation does not survive a matched null, as set out in Supplementary Results. Reference-pool frequencies were 0.912 (MOL), 0.026 (QUI) and 0.000 (PIP).

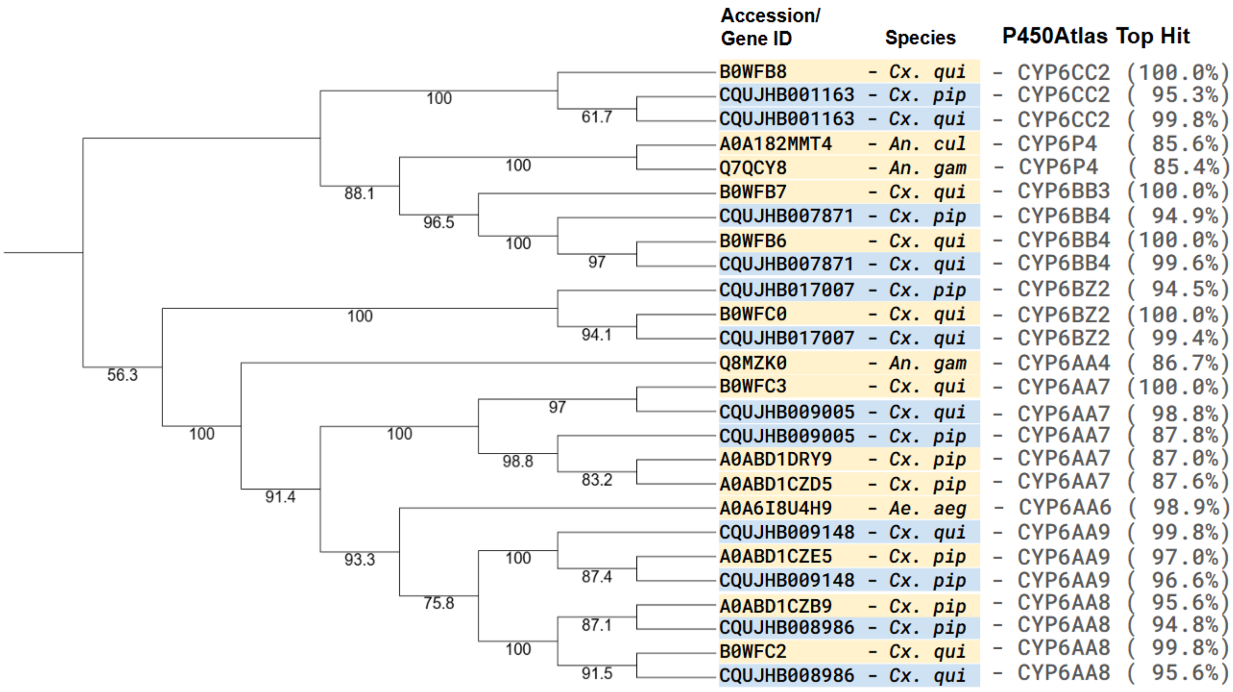


**Figure S8. The six CpL1a CYP6 paralogs belong to multiple CYP6 subgroups, not a single repeated gene.** Maximum-likelihood amino-acid phylogeny of 26 cytochrome P450 proteins: the six focal CpL1a cluster genes, each represented by its copy in both the *Cx. pipiens* (idCulPipi1, lifted annotation) and *Cx. quinquefasciatus* (JHB2020) assemblies (12 sequences, blue), together with a curated panel of 14 representative mosquito P450 references (yellow) spanning *Cx. pipiens*, *Cx. quinquefasciatus*, *Anopheles gambiae*, *Anopheles culicifacies*, and *Aedes aegypti*. Tips are labeled with UniProtKB or gene accession, source species (*Cx. qui*, *Cx. quinquefasciatus*; *Cx. pip*, *Cx. pipiens*; *An. gam*, *An. gambiae*; *An. cul*, *An. culicifacies*; *Ae. aeg*, *Ae. aegypti*), and the P450Atlas top match with percent identity, the basis for the descriptive "-like" labels used in the text. Trees were inferred in IQ-TREE 3.1.1 with the amino-acid substitution model selected by ModelFinder; node labels are SH-aLRT support. Each focal pair groups tightly with its cross-assembly counterpart but embeds within a distinct CYP6 subgroup (CYP6AA, CYP6BB, CYP6BZ, CYP6CC) rather than forming a single clade, identifying the locus as a multi-gene CYP6 paralog cluster rather than a tandem array of one repeated gene.

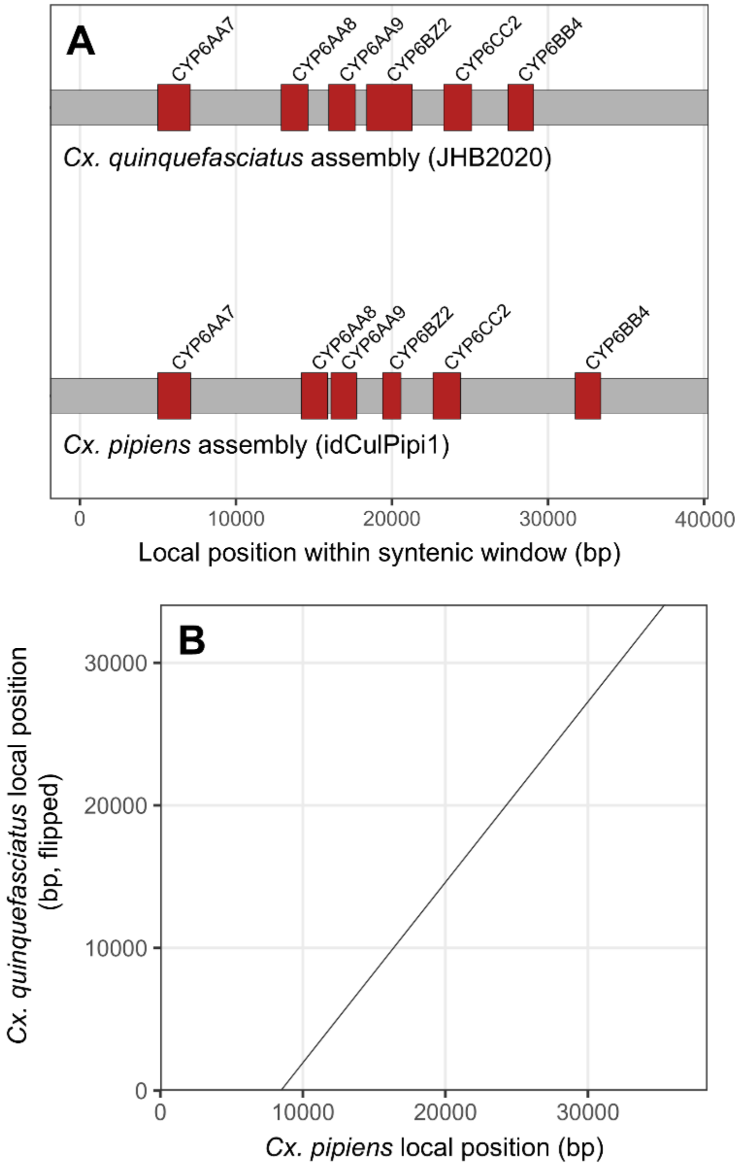


**Figure S9. The CpL1a CYP6 cluster is structurally conserved between the *Cx. pipiens* and *Cx. quinquefasciatus* assemblies.** (A) Gene-order schematic for the six focal CYP6 paralogs within the syntenic chromosome-1 window of the *Cx. quinquefasciatus* JHB2020 assembly (top) and the *Cx. pipiens* idCulPipi1 reference (bottom). Red boxes are gene models, labeled by their P450Atlas "-like" assignment; the gray bar is the chromosomal interval. The six genes occur in the same order in both assemblies (CYP6AA7, CYP6AA8, CYP6AA9, CYP6BZ2, CYP6CC2, CYP6BB4). (B) Dot-plot of the minimap2 alignment between the two windows (primary-filtered hits, alignment length ≥ 1 kb, MAPQ ≥ 5), with *Cx. pipiens* local position on the x-axis and *Cx. quinquefasciatus* local position on the y-axis. The two assemblies aligned in opposite orientation, so *Cx. quinquefasciatus* coordinates were reversed ("flipped") for comparison; the single clean positive diagonal confirms a colinear, co-oriented alignment with no internal rearrangement across the cluster.

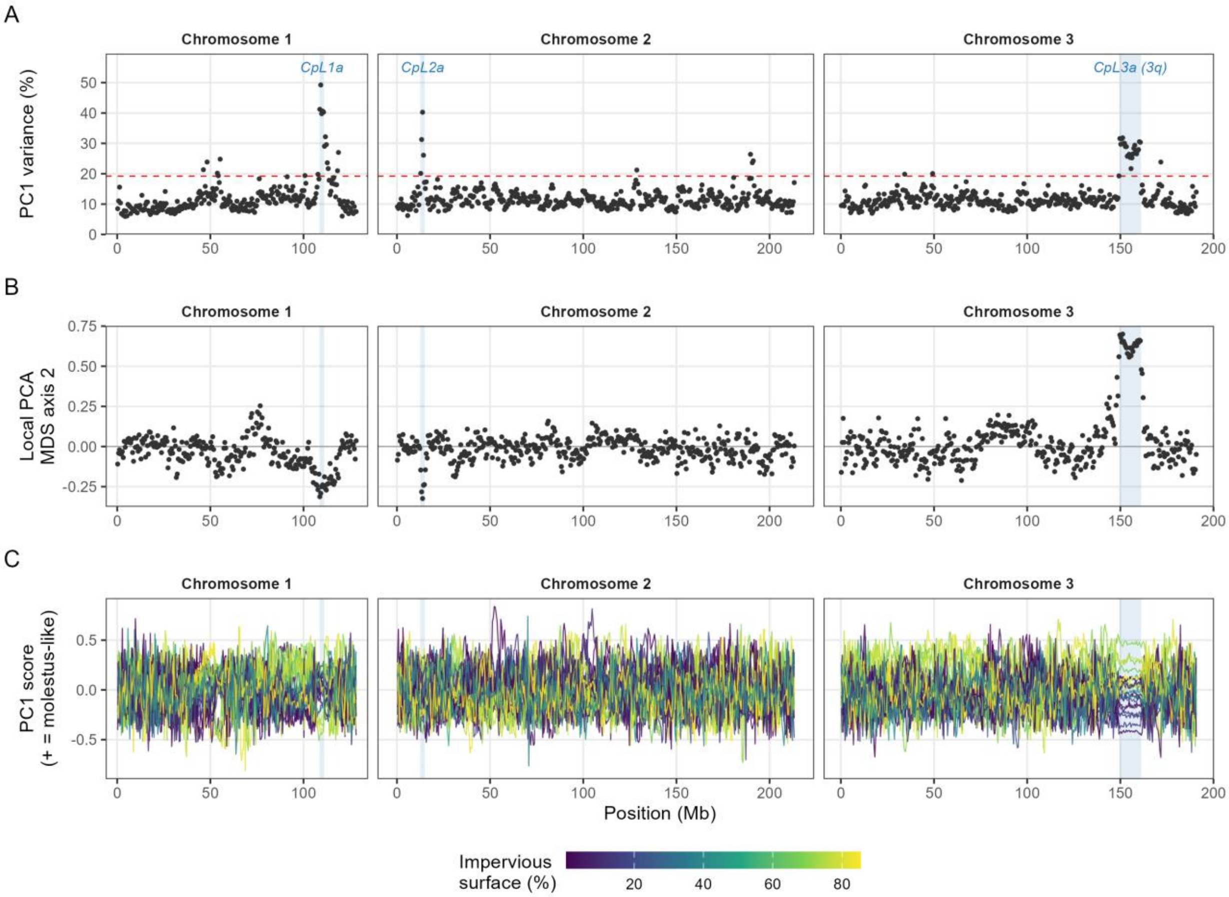


**Figure S10. A genome-wide windowed PCA of field-pool allele frequencies recovers the CpL3a inversion without prior coordinates.** Principal component analysis of the 24 field pools in 1,066 non-overlapping 500 kb windows (Supplementary Methods), the pooled counterpart of the windowed PCA used by Cebrián-Camisón et al. (2026) to detect inversions in individually sequenced *Cx. pipiens*. (A) Percentage of among-pool variance explained by PC1 in each window; the dashed red line marks the 95th percentile across windows. (B) Position of each window on the second axis of a multidimensional scaling of distances between the local PCA structures of windows (Li and Ralph 2019); windows sharing a pattern of pool similarity take similar values. (C) PC1 score of each field pool in each window, oriented so that positive values are more *molestus*-like (lower $F_{ST}$ to MOL), colored by percent impervious surface at the collection site. Blue bands mark the three candidate loci; CpL3a coincides with inversion 3q of Cebrián-Camisón et al. (2026). CpL3a is the only extended block in the genome with both elevated PC1 variance and a shared, distinct local structure, and within it pools hold stable positions along PC1, whereas outside it their ordering varies from window to window. Because pools average over individuals, the scan detects an inversion only where its frequency varies among populations, and it cannot assign karyotypes.

**Supplementary Tables**

Tables S3, S5, S7 and S8 are provided as separate worksheets in Supplementary_Data.xlsx; their legends are given below in sequence.

**Table S1. Collection sites and environmental predictors for the 24 field pools.** Each pool was collected at a distinct site across the Baltimore–Washington corridor, listed by site number and short name with coordinates in decimal degrees (°N, °W). The metropolitan area column gives the assignment of each site to whichever city center is nearer in great-circle distance; this assignment enters every locus-level environmental model as a fixed effect. The eight environmental predictors used in the genome scan are grouped by gradient. Temperature gradient: bio05, maximum temperature of the warmest month (°C); bio07, temperature annual range (°C). Urbanization gradient: PIS, percent impervious surface (%); Tree, tree canopy cover (%); PD, human population density (natural log of persons $km^{-2}$). Context variables: Elev, elevation (m); WTD, water-table depth (m); bio18, precipitation of the warmest quarter (mm). Within each gradient, predictors are more strongly correlated with one another than across gradients, so we interpret each association as a response to its gradient rather than to any single predictor. All values are reported at the precision at which they enter the models; unrounded values are deposited with the analysis code.

| # | Site | Coords (°N, °W) | Metro | bio05 | bio07 | PIS | Tree | PD | Elev | WTD | bio18 |
|---|---|---|---|---|---|---|---|---|---|---|---|
| 1 | ColHeights | 38.92955, -77.02741 | Washington | 30.9 | 33.5 | 70.5 | 7.1 | 9.4 | 52 | 6.6 | 310 |
| 2 | CHC | 38.91161, -77.03632 | Washington | 31.1 | 33.5 | 81.8 | 2.7 | 9.6 | 33 | 3.3 | 304 |
| 3 | NatHist | 38.89126, -77.02606 | Washington | 31.1 | 33.5 | 78.5 | 2.8 | 5.4 | 8 | 1.0 | 296 |
| 4 | Cst | 38.88587, -77.01776 | Washington | 31.2 | 33.5 | 70.4 | 5.1 | 5.1 | 7 | 1.3 | 293 |
| 5 | Cheverly | 38.91516, -76.90947 | Washington | 30.9 | 33.1 | 53.7 | 21.9 | 3.6 | 19 | 3.1 | 303 |
| 6 | UM | 38.85905, -76.77863 | Washington | 30.4 | 32.1 | 11.6 | 35.8 | 4.6 | 28 | 9.5 | 325 |
| 7 | SERC | 38.88785, -76.55342 | Washington | 29.2 | 30.1 | 0.5 | 69.0 | 3.1 | 6 | 4.6 | 348 |
| 8 | Bowie | 38.97728, -76.77058 | Washington | 30.1 | 32.2 | 22.3 | 45.3 | 6.8 | 54 | 9.9 | 337 |
| 9 | Beltsville | 39.00999, -76.82407 | Washington | 30.3 | 32.6 | 4.2 | 52.9 | 2.2 | 51 | 6.6 | 327 |
| 10 | ColPark | 39.01578, -76.93413 | Washington | 30.8 | 33.4 | 18.9 | 34.2 | 6.9 | 38 | 4.3 | 306 |
| 11 | Pax | 39.05560, -76.82215 | Washington | 30.4 | 32.8 | 9.5 | 59.1 | 5.8 | 47 | 11.9 | 319 |
| 12 | SSC | 39.03942, -76.59501 | Baltimore | 29.2 | 30.6 | 2.5 | 76.9 | 4.7 | 31 | 15.4 | 341 |
| 13 | QP | 39.15120, -76.66517 | Baltimore | 29.7 | 31.8 | 23.7 | 41.5 | 5.2 | 37 | 3.2 | 323 |
| 14 | WCP | 39.18274, -76.78570 | Baltimore | 30.0 | 32.8 | 38.1 | 33.9 | 7.7 | 78 | 5.6 | 324 |
| 15 | Jerald | 39.11010, -76.90164 | Washington | 30.1 | 32.9 | 15.8 | 46.8 | 6.4 | 112 | 5.6 | 330 |
| 16 | Brighton | 39.19042, -77.00528 | Washington | 30.1 | 33.1 | 1.1 | 61.5 | 4.3 | 114 | 7.7 | 302 |
| 17 | Clarksville | 39.25637, -76.92767 | Baltimore | 30.0 | 33.0 | 1.5 | 19.5 | 3.9 | 112 | 5.9 | 308 |
| 18 | ElCity | 39.28590, -76.86023 | Baltimore | 29.8 | 32.9 | 15.4 | 53.3 | 6.7 | 126 | 4.6 | 311 |
| 19 | Hillcrest | 39.24479, -76.65343 | Baltimore | 29.7 | 32.0 | 43.6 | 18.0 | 7.7 | 33 | 16.9 | 302 |

| # | Site | Coords (°N, °W) | Metro | bio05 | bio07 | PIS | Tree | PD | Elev | WTD | bio18 |
|---|---|---|---|---|---|---|---|---|---|---|---|
| 20 | UMBC | 39.28889, -76.62303 | Baltimore | 29.6 | 31.9 | 85.3 | 2.2 | 8.1 | 15 | 3.1 | 299 |
| 21 | IMET | 39.28555, -76.60513 | Baltimore | 29.6 | 31.7 | 77.8 | 0.8 | 8.0 | 5 | 1.3 | 298 |
| 22 | UFP | 39.29039, -76.58647 | Baltimore | 29.4 | 31.5 | 66.8 | 2.1 | 8.6 | 13 | 7.9 | 300 |
| 23 | NorthBroad | 39.30278, -76.59423 | Baltimore | 29.4 | 31.7 | 76.8 | 3.3 | 8.4 | 32 | 6.1 | 304 |
| 24 | AGCB | 39.36769, -76.87747 | Baltimore | 29.8 | 33.1 | 2.9 | 59.8 | 3.9 | 115 | 18.8 | 290 |

**Table S2. Genome-wide pairwise $F_{ST}$ between each field pool and the three reference pools.** $F_{ST}$ computed genome-wide in poolfstat using the analysis-of-variance estimator, with each pool treated as 80 chromosomes (40 diploid individuals), for the 24 field pools against *Cx. pipiens* f. *pipiens* (PIP), *Cx. p.* f. *molestus* (MOL), and *Cx. quinquefasciatus* (QUI). Site numbers match Figs. 1–2. Every pool is least differentiated from PIP and most differentiated from QUI, with MOL intermediate, consistent with a predominantly *Cx. pipiens*-like genomic background across the corridor.

| Site Name | Site Number | PIP | MOL | QUI |
|---|---|---|---|---|
| ColHeights | 1 | 0.172 | 0.257 | 0.538 |
| CHC | 2 | 0.170 | 0.258 | 0.538 |
| NatHist | 3 | 0.177 | 0.257 | 0.548 |
| Cst | 4 | 0.172 | 0.257 | 0.539 |
| Cheverly | 5 | 0.171 | 0.260 | 0.538 |
| UM | 6 | 0.171 | 0.267 | 0.534 |
| SERC | 7 | 0.174 | 0.270 | 0.548 |
| Bowie | 8 | 0.171 | 0.265 | 0.528 |
| Beltsville | 9 | 0.170 | 0.261 | 0.524 |
| ColPark | 10 | 0.172 | 0.261 | 0.528 |
| Pax | 11 | 0.176 | 0.274 | 0.544 |
| SSC | 12 | 0.177 | 0.276 | 0.543 |
| QP | 13 | 0.175 | 0.273 | 0.540 |
| WCP | 14 | 0.174 | 0.276 | 0.553 |
| Jerald | 15 | 0.176 | 0.274 | 0.547 |
| Brighton | 16 | 0.174 | 0.267 | 0.542 |
| Clarksville | 17 | 0.174 | 0.275 | 0.548 |
| ElCity | 18 | 0.177 | 0.275 | 0.542 |
| Hillcrest | 19 | 0.175 | 0.276 | 0.556 |
| UMBC | 20 | 0.176 | 0.267 | 0.538 |
| IMET | 21 | 0.181 | 0.273 | 0.554 |
| UFP | 22 | 0.181 | 0.268 | 0.547 |
| NorthBroad | 23 | 0.179 | 0.268 | 0.549 |
| AGCB | 24 | 0.175 | 0.269 | 0.542 |

**Table S3 (Supplementary_Data.xlsx, worksheet "Table S3"). Nonsynonymous variation across canonical insecticide-resistance target and detoxification genes.** The panel of 63 genes was surveyed independently of the genome scan; the table lists the 60 that carry at least one nonsynonymous variant (the other three, an acetylcholinesterase and two esterase B1 genes, carry none). For each gene, the table reports the number of nonsynonymous variants segregating across the 24 field pools, the maximum mean allele frequency among those variants, the number of canonical resistance-associated sites (residue-match flags for *Ace-1* and *Rdl* are candidates based on projected annotation rather than confirmed sites; see Supplementary Methods), and the minimum univariate P value for association with maximum summer temperature and percent impervious surface (PIS) across the 24 field pools. The six CpL1a CYP6 paralogs are included for completeness, and the sodium channel 60E gene is a labeled negative control rather than a resistance target. No environmental association survives correction for multiple testing. The per-variant detail underlying these gene-level summaries is deposited with the analysis code (Data Availability).

**Table S4. Donor-private panel sizes by region.** Counts of SNPs private to each reference lineage, at a frequency of at least 0.95 in that reference and no more than 0.05 in both others, within each of the three candidate intervals and across the remaining genomic background. The second column gives the number of positions in the region at which all three references have a called frequency, which is the pool from which each panel is drawn. Two features of this table bear on what each locus can support. First, the asymmetry among donors is expected rather than anomalous: a reference with higher heterozygosity is less often fixed for an allele absent from both others, so the most diverse reference, PIP, yields the smallest panel in every region. Second, the counts set the limits of donor attribution directly. CpL2a contains no PIP-private SNPs at all and CpL1a contains a single one, which is why ancestry at those loci is reported as QUI measured against both alternatives rather than as a three-way partition, and why the per-population test at CpL2a uses the combined non-QUI fraction. At CpL3a the QUI and PIP panels are likewise too small to support an estimate, and only the MOL panel is used.

| Region | SNPs called in all three references | QUI-private | PIP-private | MOL-private |
|---|---|---|---|---|
| CpL1a | 73,073 | 123 | 1 | 3,762 |
| CpL2a | 10,866 | 6 | 0 | 30 |
| CpL3a | 245,296 | 4 | 3 | 3,449 |
| Genomic background | 11,300,655 | 5,884 | 440 | 116,508 |
| *Genome-wide total* | *11,629,890* | *6,017* | *444* | *123,749* |

**Table S5 (Supplementary_Data.xlsx, worksheet "Table S5"). Candidate-SNP density among the protein-coding genes of the CpL1a interval.** For the 15 highest-ranked of the 79 protein-coding genes in the chromosome 1 candidate interval (108.5–111.0 Mb), the table reports the number and density of BayPass candidate SNPs over the gene body and over CDS only, the maximum per-SNP $XtX_{st}$ in the region, and the lifted gene description. Gene-body counts and densities are computed over the gene span extended by a ±2 kb flank to capture proximal regulatory sequence, so gene-body density is candidate SNPs per kb of gene length + 4 kb; CDS density is candidate SNPs in the isoform-union CDS per kb of CDS. Genes are ranked by CDS SNP density; the six CYP6 paralogs occupy ranks 1–6 on that metric and on gene-body density alike. Raw counts rank differently, and the two genes that exceed the CYP6 paralogs on count are discussed in Supplementary Results.

**Table S6. Nonsynonymous coding consequences of the temperature-associated CYP6 candidate SNPs at CpL1a.** The 13 focal coding-sequence SNPs predicted to cause a nonsynonymous (missense) change across the six CpL1a CYP6 paralogs. For each substitution the table gives the genomic position, BayPass differentiation ($XtX_{st}$) and the most strongly associated environmental predictor (bio05, maximum summer temperature; PIS, percent impervious surface), the reference and alternate alleles, the alternate-allele frequency in each reference pool (QUI, PIP, MOL), and the resulting codon change, amino-acid change, and protein position. Consequences were predicted with predictCoding (VariantAnnotation) against transcript models from the lifted idCulPipi1 annotation. Because idCulPipi1 represents the *Cx. pipiens* genomic background, the alternate allele is the candidate introgressed state, and an alternate allele near-fixed in QUI and absent from MOL is diagnostic for the *Cx. quinquefasciatus* (QUI) donor. Reference and alternate alleles are shown on each gene's coding strand, so that the alleles, codon, and amino acid are internally consistent within each row. All six paralogs are transcribed on the minus strand, so these coding-strand alleles are the reverse complement of the forward-strand alleles recorded at the listed genomic positions; for example, the C→A substitution shown at position 109074507 is recorded as G→T on the forward (+) strand in a VCF or genome browser. Of the 13 substitutions, 11 are QUI-diagnostic and associated with the summer-temperature gradient, one (Q411H, CYP6BB4-like) is QUI-diagnostic but tracks percent impervious surface, and one (N469S, CYP6AA7-like) segregates at comparable frequency in QUI and PIP and is therefore not donor-diagnostic.

| Gene | Position on Chr1 | $XtX_{st}$ | Predictor | Ref. Allele | Alt. Allele | QUI Alt. Freq. | PIP Alt. Freq. | MOL Alt. Freq. | Reference Codon | Variant Codon | Reference Amino Acid | Variant Amino Acid | Protein Position |
|---|---|---|---|---|---|---|---|---|---|---|---|---|---|
| CYP6AA7-like | 109074507 | 186.7 | bio05 | C | A | 1.00 | 0.81 | 0.00 | ACG | AAG | T | K | 503* |
| | 109074609 | 236.0 | bio05 | A | G | 0.56 | 0.56 | 0.00 | AAC | AGC | N | S | 469 |
| | 109076033 | 197.6 | bio05 | G | A | 1.00 | 0.68 | 0.00 | GTT | ATT | V | I | 18 |
| CYP6AA8-like | 109083729 | 240.0 | bio05 | G | A | 1.00 | 0.73 | 0.00 | AGC | AAC | S | N | 327 |
| CYP6AA9-like | 109085707 | 233.8 | bio05 | A | G | 1.00 | 0.72 | 0.00 | ACA | GCA | T | A | 335 |
| | 109085731 | 198.3 | bio05 | C | A | 1.00 | 0.73 | 0.00 | CAC | AAC | H | N | 327 |
| | 109086213 | 264.5 | bio05 | T | C | 1.00 | 0.82 | 0.00 | GTG | GCG | V | A | 166 |
| CYP6BZ2-like | 109088676 | 217.7 | bio05 | T | G | 0.97 | 0.74 | 0.00 | AAT | AAG | N | K | 256 |
| | 109088955 | 188.9 | bio05 | A | T | 1.00 | 0.78 | 0.00 | GAA | GAT | E | D | 163 |

| Gene | Position on Chr1 | $XtX_{st}$ | Predictor | Ref. Allele | Alt. Allele | QUI Alt. Freq. | PIP Alt. Freq. | MOL Alt. Freq. | Reference Codon | Variant Codon | Reference Amino Acid | Variant Amino Acid | Protein Position |
|---|---|---|---|---|---|---|---|---|---|---|---|---|---|
| CYP6CC2-like | 109092779 | 204.5 | bio05 | G | T | 1.00 | 0.72 | 0.00 | TTG | TTT | L | F | 142 |
| CYP6BB4-like | 109101089 | 198.2 | PIS | A | C | 1.00 | 0.76 | 0.00 | CAA | CAC | Q | H | 411 |
| | 109101372 | 266.9 | bio05 | G | C | 1.00 | 0.74 | 0.00 | GAA | CAA | E | Q | 335 |
| | 109102092 | 120.0 | bio05 | G | T | 1.00 | 0.63 | 0.00 | GGC | TGC | G | C | 95 |

*Codon 503 contains a second focal CDS SNP: a synonymous change at the adjacent nucleotide (position 109074506). Because pooled data are unphased, the two SNPs are scored independently against the reference codon, so this T503K call assumes the reference base at the partner site. If both introgressed alleles lie on the same haplotype, the combined codon (AAC) encodes asparagine (T503N).

**Table S7 (Supplementary_Data.xlsx, worksheet "Table S7"). Genes carrying candidate SNPs across the CpL2a interval.** Protein-coding genes within the 500 kb CpL2a interval that carry at least one candidate SNP in the gene body, ordered by gene-body candidate-SNP count. For each gene: the number of candidate SNPs falling within the gene body (gene span extended by 2 kb on each side) and within coding sequence, the maximum single-SNP BayPass differentiation (Max $XtX_{st}$) in the gene-plus-flank interval, and the gene description from the lifted idCulPipi1 annotation.

**Table S8 (Supplementary_Data.xlsx, worksheet "Table S8"). Genes carrying coding candidate SNPs across the CpL3a inversion span.** Protein-coding genes within the approximately 11.6 Mb CpL3a inversion that carry at least one candidate SNP in coding sequence, ordered by gene-body candidate-SNP count. For each gene: the number of candidate SNPs falling within the gene body (gene span extended by 2 kb on each side) and within coding sequence, the maximum single-SNP BayPass differentiation (Max $XtX_{st}$) in the gene-plus-flank interval, and the gene description from the lifted idCulPipi1 annotation. Coding candidate-SNP counts are low, no gene carrying more than two, and are spread across genes of varied function, in contrast to the localized peaks at CpL1a and CpL2a where single genes carried tens of candidate SNPs. Because the inversion rather than any individual gene is the unit of selection under suppressed recombination, these genes are reported as context only and none is nominated as a target of selection.

**Table S9. Reference-pool collection localities and coordinates.** Collection coordinates (latitude, longitude in decimal degrees) for the three lineage-reference pools used to polarize donor ancestry: QUI (*Cx. quinquefasciatus*; Vero Beach, FL), the candidate donor for the temperature-associated CpL1a signal; PIP (*Cx. pipiens* f. *pipiens*; Orono, ME), representing the temperate *Cx. pipiens* background; and MOL (*Cx. p.* f. *molestus*; Calumet, IL), the candidate donor for the urbanization-associated CpL3a inversion. Each reference pool was constructed from 40 specimens, as for the field pools (Methods).

| Population | Collection Coordinates (Latitude, Longitude) |
|---|---|
| QUI | 27.568669, -80.417803 |
| MOL | 41.650225, -87.600140 |
| PIP | 44.892258, -68.671079 |
